# Discovery and visualization of uncharacterized drug-protein adducts using mass spectrometry

**DOI:** 10.1101/2021.06.24.449838

**Authors:** Michael Riffle, Michael R. Hoopmann, Daniel Jaschob, Guo Zhong, Robert L. Moritz, Michael J. MacCoss, Trisha N. Davis, Nina Isoherranen, Alex Zelter

## Abstract

Drugs are often metabolized to reactive intermediates that form protein adducts. Adducts can inhibit protein activity, elicit immune responses, and cause life threatening adverse drug reactions. The masses of reactive metabolites are frequently unknown, rendering traditional mass spectrometry-based proteomics incapable of adduct identification. Here, we present Magnum, an open-mass search algorithm optimized for adduct identification, and Limelight, a web-based data processing package for analysis and visualization of data from all existing algorithms. Limelight incorporates tools for sample comparisons and xenobiotic-adduct discovery. We validate our tools with two drug/protein combinations and apply our workflow to identify novel xenobiotic-protein adducts in CYP3A4. Our new methods and software enable accurate identification of xenobiotic-protein adducts with no prior knowledge of adduct masses or protein targets. Magnum outperforms existing tools in xenobiotic-protein adduct discovery, while Limelight fulfills a major need in the rapidly developing field of open-mass searching, which until now lacked comprehensive data visualization tools.

## Main

Humans are constantly exposed to chemicals from their environment causing adverse effects and premature aging^1^. Protein adducts result from covalent modification by xenobiotics, or their metabolites, and can also cause unintended toxicities and adverse drug reactions^2–4^. Adduct identification improves understanding of the mechanisms of drug induced toxicities and enables design of structural modifications to prevent them. Current identification methods, using radiolabeled compounds plus trapping agents, are labor-intensive and require extensive experimentation^5–7^ to gain specific information about the adducts’ chemical nature or the proteins modified. Chromatographic, immunological and mass spectrometry-based methods for detecting xenobiotic-protein adducts have been developed, but cannot sensitively measure the presence, abundance and localization of protein adducts across a range of proteins without prior knowledge^2,8,9^.

Traditional database search algorithms^10^ search MS/MS spectra against protein sequences to identify the peptides and proteins that produced them. Known modifications can be identified if their masses are predefined. This is not possible for xenobiotic-protein adducts if their chemical composition is unknown. “Open-mass” search strategies^11–19^ solve this issue by allowing observed peptide masses to differ from identified peptide masses, returning mass differences as modifications of the identified peptides. This has allowed peptide spectrum matches (PSMs) to be made from a large proportion of previously unassigned spectra in shotgun proteomics data^11–19^. However, few algorithms or workflows have been designed to identify xenobiotic-protein adducts and there are no graphical software tools to analyze and visualize data from open-mass searches.

Here we present Magnum, a purpose-built xenobiotic-protein adduct discovery algorithm, and Limelight, a web-based open modification analysis platform that rapidly highlights protein adducts that result from a specific treatment in a background of unrelated modifications. Limelight can combine and compare data from different pipelines empowering users to find the best tool or combination of tools for their specific application.

We validate our new tools using two drug/protein combinations and compare results from Magnum to several previously published open-mass search tools. We apply our workflow to identify novel xenobiotic-protein adducts in the P450 enzyme CYP3A4 resulting from exposure to raloxifene. Our software and workflow enable rapid and accurate identification of novel xenobiotic-protein adducts with no prior knowledge of adduct masses or protein targets. These tools provide a highly accelerated and statistically rigorous workflow for the discovery and characterization of xenobiotic-protein adducts and can be incorporated into drug discovery pipelines and environmental toxicology screening.

## Results

### Development of Magnum

Magnum was developed to analyze MS/MS spectra to identify peptide sequences modified by adducts of unknown mass (Figure 1; Supplementary Note 1). The approach taken in Magnum is similar to cross-linked peptide identification in Kojak^20^. However, in Kojak search results contain two peptide sequences that sum together to produce the observed mass, whereas Magnum returns only one peptide sequence plus a modification mass equal to the mass unexplained by the predicted peptide.

**Figure 1:**
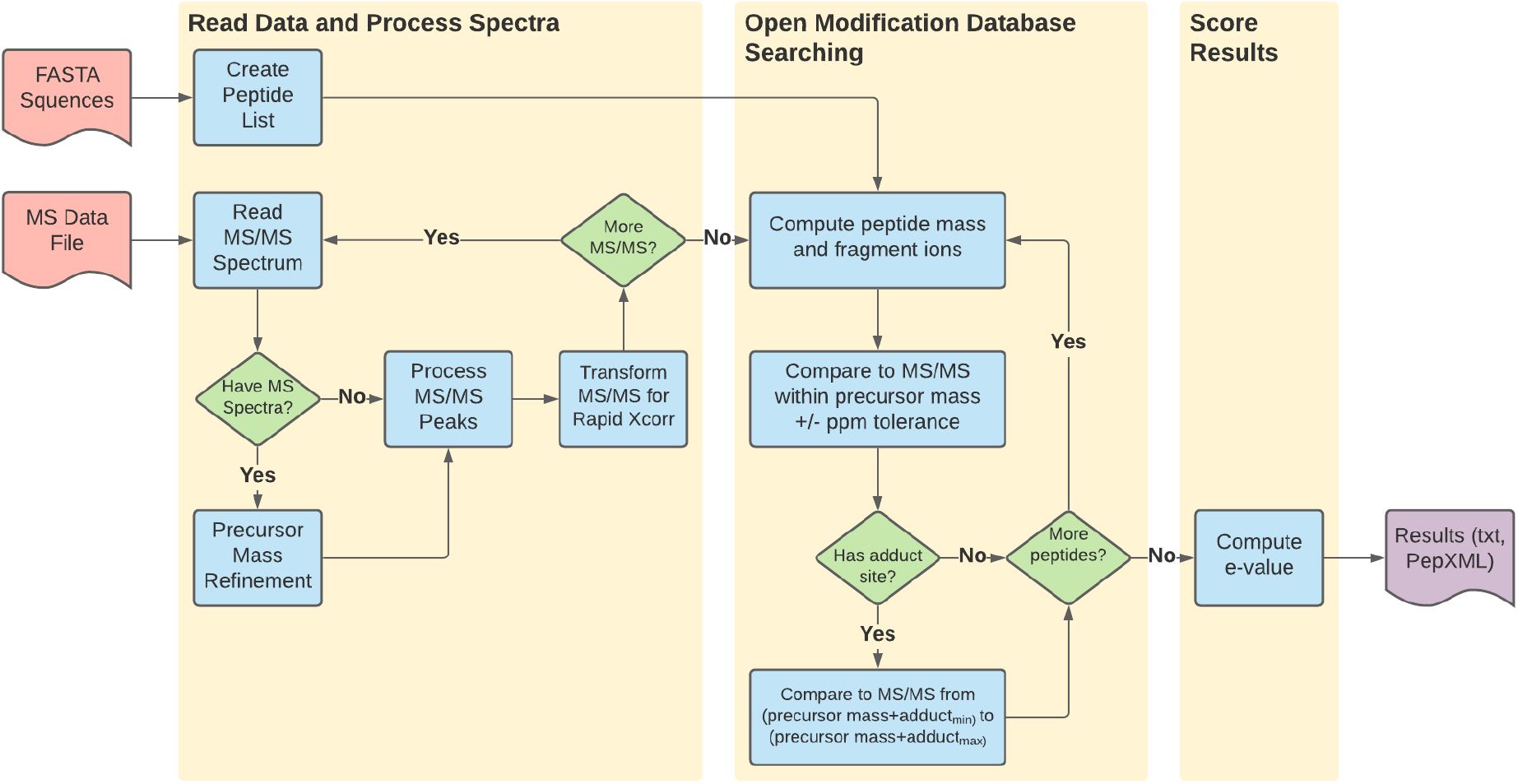
Magnum data processing workflow. Magnum architecture is divided into three primary categories: reading and processing spectra, open modification database searching, and scoring. Input files are a protein FASTA sequences file and spectral data in an open format (e.g. mzML). Output from Magnum is in both tab-delimited text and PepXML format.

Magnum incorporates several features that improve its ability to identify protein adducts. First, Magnum calculates and scores an adduct-modified MS/MS ion series. Many open search algorithms disregard adduct-modified fragment ions leading to a loss of sensitivity and preventing adduct localization^11,16–18^. Magnum handles both MS-labile and non-labile modifications by calculating and scoring a modified and unmodified ion series allowing adduct identification with or without adduct localization. Second, Magnum allows restricting adduct localization to specific amino acids during searching. This is useful if the reactivity of a xenobiotic is known or hypothesized based on its chemistry or through detection of thiol conjugates with reactive intermediates^2^. Increased search sensitivity and statistical power can be gained by restricting the open modification search space to specific amino acids^21,22^. Third, Magnum can flag MS/MS spectra containing diagnostic reporter ions. Some xenobiotics and endogenous post-translational modifications (PTMs) fragment predictably in the mass spectrometer producing reporter ions^23–25^, the masses of which are constant regardless of the peptide to which the adduct was attached. If present, Magnum records this information in the PSM. Fourth, Magnum considers user defined peptide modifications and open modifications. This allows separation of masses due to defined versus undefined modifications and is important if both types of modification exist on a single peptide.

The product of a Magnum analysis is a set of PSMs for as many input spectra as possible, plus a set of metrics that can be used as input in a variety of PSM-validation^26,27^ algorithms.

### Evaluation of open modification search tools for xenobiotic-protein adduct discovery

Open-mass searches have greater potential for false positive identifications than closed searches because discrepancies between theoretical and observed peptide masses are interpreted as open modifications. While enabling identification of undefined modifications, this can lead to incorrect identifications not otherwise possible. We created a gold standard (true positive) dataset to evaluate the accuracy and sensitivity of Magnum in identifying xenobiotic-protein adducts. We also compared Magnum, which was designed specifically for xenobiotic-protein adduct detection, to several previously published open search algorithms.

For our gold standard dataset, we acquired high resolution LC-MS/MS data of human serum albumin (HSA) exposed to the β-lactam antibiotics dicloxacillin and flucloxacillin. A significant portion of previous research identifying protein adducts from environmental exposures has focused on HSA^28^ as it is the most abundant protein in plasma and forms adducts with numerous xenobiotics^9^. The β-lactam antibiotics dicloxacillin and flucloxacillin were chosen as there are published MS characterizations of their clinically significant adducts^23,29,30^.

Our gold standard dataset consisted of 2,979 MS/MS spectra extracted from 307,652 scans acquired from two dicloxacillin- and two flucloxacillin-treated HSA samples. Each of the 2,979 spectra was definitively determined to result from a peptide containing one 469 Da (dicloxacillin) or 453 Da (flucloxacillin) adduct using the methods described in Supplementary Note 2. The four full datasets (307,652 MS/MS scans) were searched with 8 algorithms. Open modification masses associated with each PSM were extracted for the 2,979 gold standard spectra. A precision/recall analysis was performed where *precision* was defined as the fraction of all answers that are correct (i.e. number of correct answers/total number of answers) and *recall* as the fraction of total possible correct answers identified (i.e. number of correct answers/number scans tested [2,979]). An answer within ±3 Da of the known modification mass was considered correct, allowing for incorrect monoisotopic mass assignments^31^, which in open searching are compensated for by changes in open modification masses. Both Magnum and MSFragger^12^ had excellent precision: >0.9 at 1% FDR (false discovery rate) (Figure 2a). Recall of Magnum was better than all other algorithms. MSFragger, comet with a 500 Da wide precursor window, and MetaMorpheus^18^ performed next best. Recall of open-pFind^13^ was close to zero as it does not perform a truly unrestricted search, but rather a multinotch search^18^, allowing only delta masses present in UniMod^32^ to be used as modification masses. As dicloxacillin and flucloxacillin adducts are not present in Unimod, open-pFind cannot identify them and only returned results for 9 of the 2,979 scans (Supplementary Table 6).

**Figure 2:**
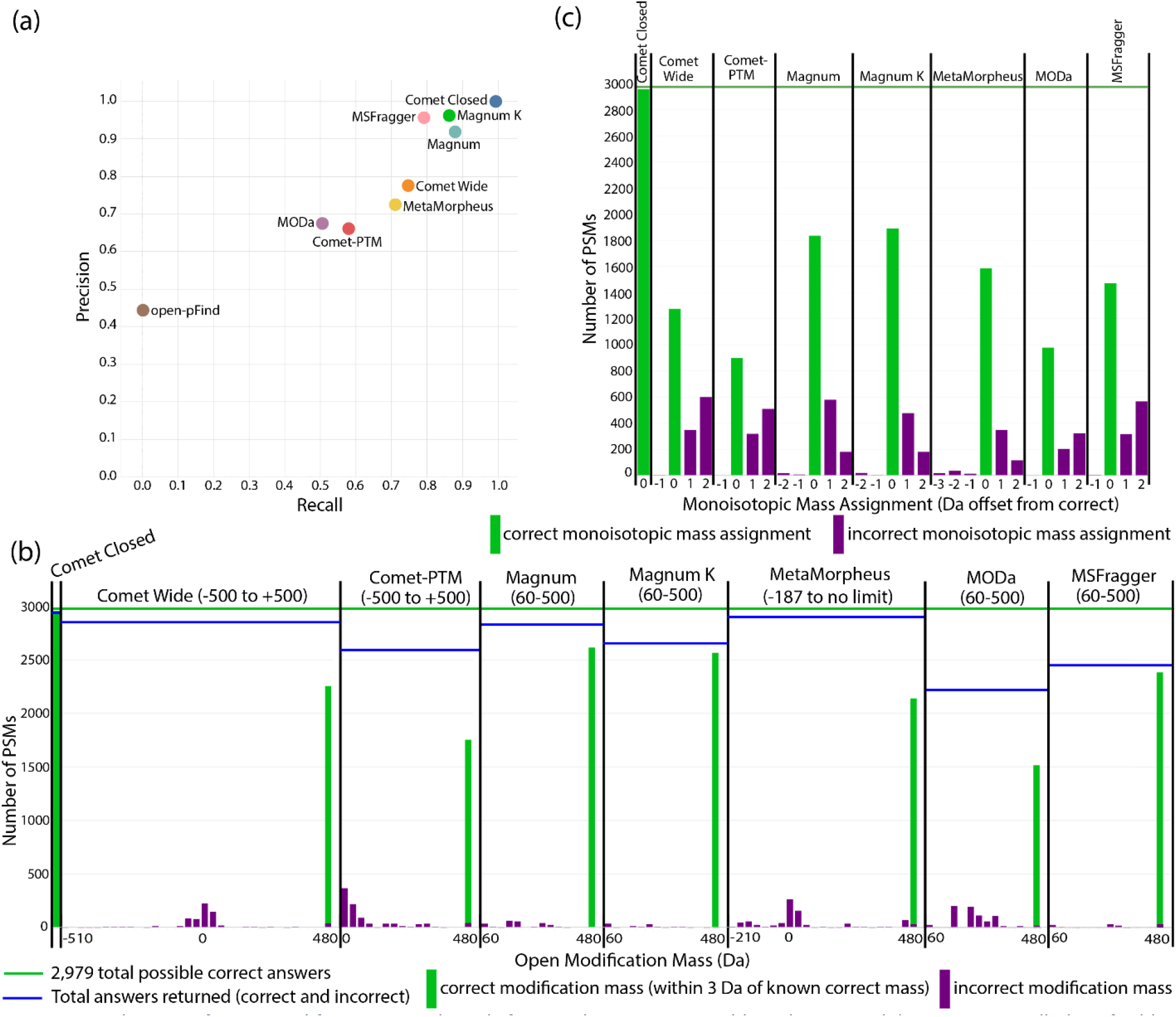
Evaluation of open modification search tools for xenobiotic-protein adduct discovery. (a) Precision recall plot of adduct masses reported at 1% FDR by 7 open search algorithms for 2,979 unique MS/MS spectra definitively determined to result from a peptide containing a single +469 Da (dicloxacillin) or +453 Da (flucloxacillin) modification. Results from closed comet searches using defined modifications of 469 or 453 were included as a positive control. An open modification mass returned by an algorithm is defined as correct if it is within ±3 Da of the known correct modification mass (469 or 453). Magnum was run allowing for open-masses on any amino acid (Magnum) or restricted to lysine only (Magnum K), the previously published residue modified by dicloxacillin and flucloxacillin adducts^23,29^; (b) Histograms showing the distribution of open-mass modifications reported by each algorithm for the 2,979 gold standard spectra at 1% FDR. Results are distributed between incorrect open-masses (purple bars, >3 Da from known correct modification mass) and correct masses (green bars, within ±3 Da of the known correct modification mass). The first and last mass bin are labeled on the x axis, which was cut off at the 480 Da bin due to sparseness in data beyond that point. The open-mass range searched by each algorithm is shown in parenthesis above each plot. Bins are 30 Da wide and all correct answers fall within the 450 Da bin. Incorrect masses within the 450 Da bin are shaded purple. (c) Histograms of monoisotopic masses assigned to each of the correct answers in (b) by each algorithm. In (c) these results are distributed between incorrect monoisotopic mass assignments (non-zero offset values, purple bars) and correct monoisotopic mass assignments (zero offset values, green bars). Bin width is 1 Da. Correct monoisotopic mass assignments have 0 Da offsets. All data in this figure is filtered at a 1% FDR. open-pFind results were excluded from (b) and (c) as only 9 out of 2,979 possible answers were returned, of which four were within 3 Da of the correct answer giving a precision of 0.44 and a recall of 0.001.

Except for open-pFind, algorithms having a low recall did so mainly due to returning incorrect modification masses rather than by not returning a result for that scan. All algorithms except open-pFind returned over 2200 total answers at 1% FDR (horizontal blue lines in Figure 2b). These were distributed between incorrect masses (purple bars in Figure 2b) and correct masses (green bars in Figure 2b).

An analysis of all open-mass modifications returned by each algorithm within ±3 Da of the known correct modification masses shows Magnum had the most accurate (62-63%) monoisotopic mass assignment of the algorithms tested (green bars, Figure 2c). MetaMorpheus and MSFragger were next best, and respectively assigned 53% and 49% of PSMs the correct monoisotopic mass. Like many pharmaceutical drugs, dicloxacillin and flucloxacillin contain chlorine, confounding monoisotopic mass assignment due to its unusual isotopic composition. To overcome these complexities, Magnum incorporates code that reduces peptide isotope distributions to a single monoisotopic mass (Supplementary Note 1). Overall, our analysis showed Magnum accurately and sensitively identifies xenobiotic-protein adducts.

### Development of Limelight

Mass spectrometry-based proteomics data is complex and open modification data even more so. Each search algorithm uses different scoring metrics and outputs results in different formats. PSM-validation tools^26,27^ are often used to assign statistical confidence to results, and add another layer of metrics to the data. If adduct data are to be assessable for answering specific questions, a simple graphical interface is needed. Limelight is a web application built to analyze, visualize, and share bottom-up MS proteomics data (Figure 3; Supplementary Note 3). It provides a generalized platform designed to support any MS database search pipeline. Data are displayed on a global level in protein, peptide, or modification centric views, which can be filtered on multiple criteria and that provide access to all underlying raw data and associated metrics from every part of the analysis. Annotated spectra, including ions modified by open-masses where applicable, are viewable using a built in spectrum viewer^33^. Data are unified and can be queried, viewed, and compared across multiple experiments and disparate software pipelines allowing the strengths of different algorithms to be leveraged.

**Figure 3:**
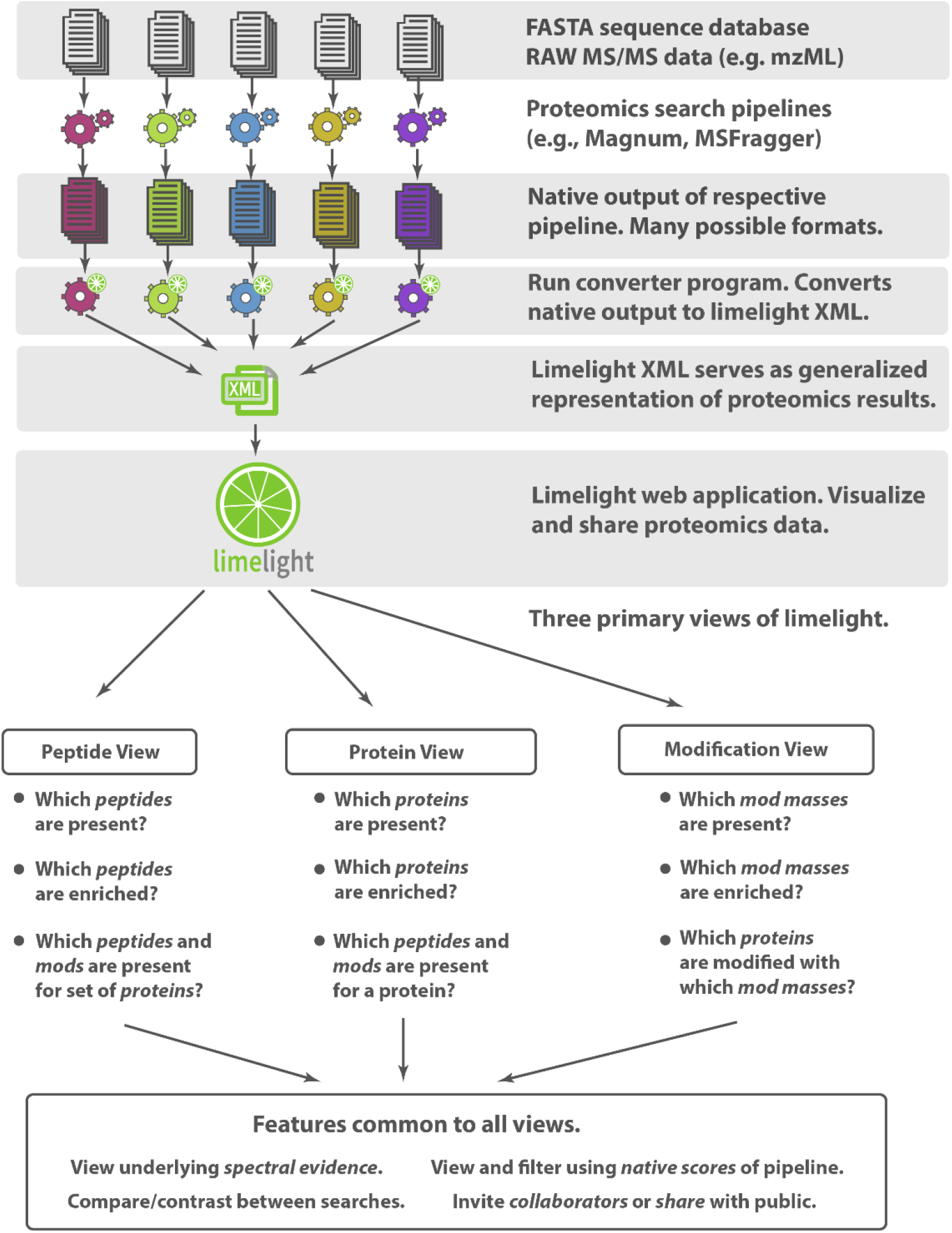
Limelight. A web-based application to interrogate, analyze and visualize mass spectrometry proteomics results, including peptide-adduct identifications, across multiple samples simultaneously. Data input is designed to support any search algorithm. Three primary views allow interaction with the data.

Limelight incorporates novel features designed for adduct and PTM analysis. Mass modifications identified in peptides may be viewed, analyzed, and visualized independently from the peptides or proteins in which they were identified and regardless of whether those modifications were localized or not. If desired, this allows interesting adduct masses to be identified and examined before segregation into protein, peptide, or residue level groups. To highlight exposure related adducts, we developed a suite of tools for visualizing and statistically comparing modification masses between different experiments within Limelight. These functions are described below.

### Development and validation of adduct discovery pipeline

Open search results contain many modification masses not related to exposures or treatment conditions. Numerous open-mass modifications are identified even in untreated purified HSA (Supplementary Figure 8, Supplementary Note 4). We therefore designed an experimental workflow to allow discovery of protein adducts resulting specifically from exposure to xenobiotics (Figure 4). Our strategy was to produce, analyze and compare both untreated and xenobiotic treated samples. We tested our pipeline using unexposed, dicloxacillin, and flucloxacillin exposed HSA.

**Figure 4:**
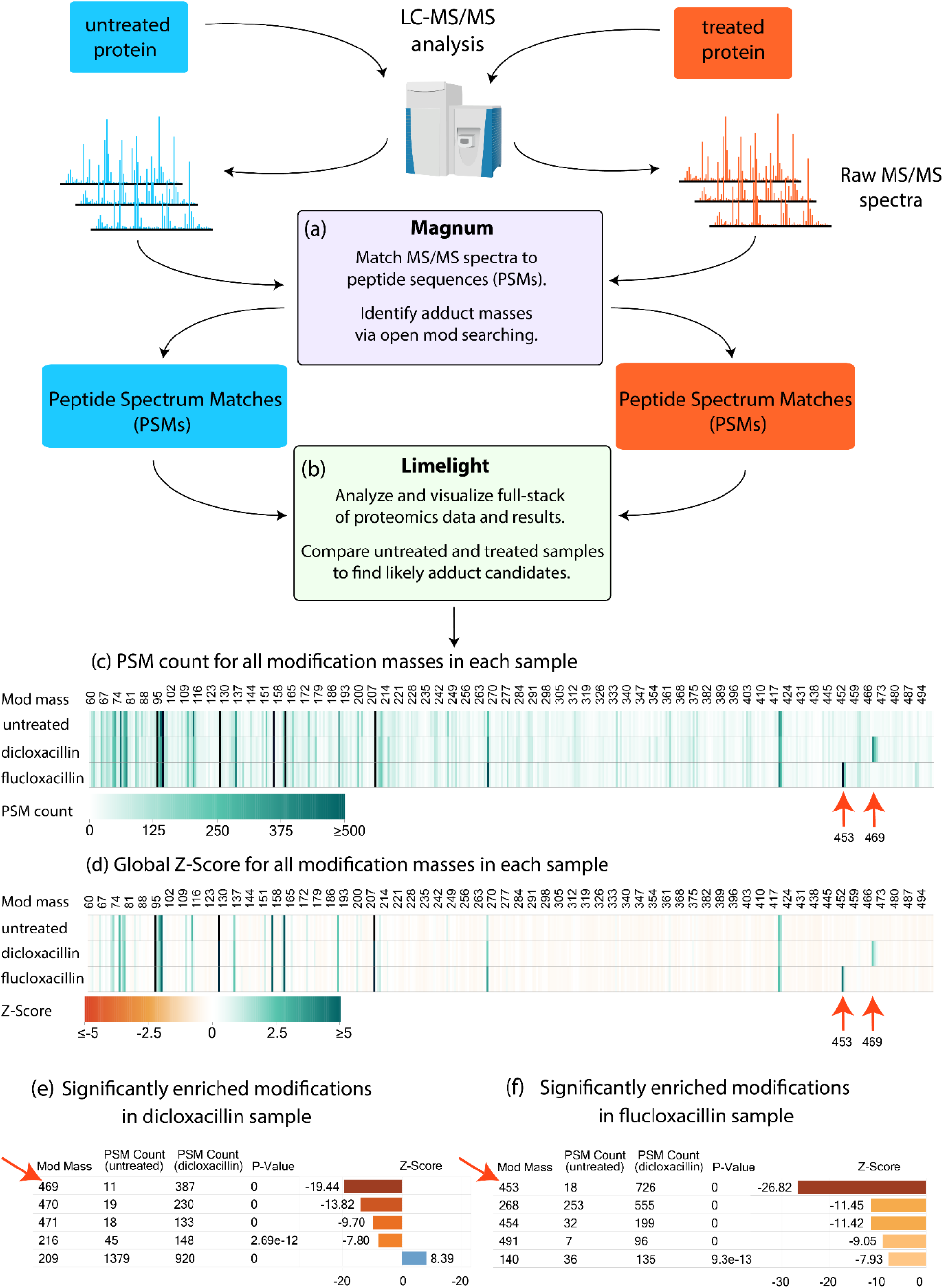
Adduct identification and visualization workflow using dicloxacillin or flucloxacillin exposed versus untreated human serum albumin (HSA) as an example. (a) Raw MS data was searched using Magnum and (b) resulting PSMs were analyzed by Limelight. Data visualization includes: (c) a heatmap of PSM count for each modification mass in each sample highlighting modification masses enriched in the treated samples (red arrows); and (d) a similar visualization to (c) but showing the calculated Global Z-Score for all modification masses across all samples; (e) results of statistical test of proportions on all modification masses automatically pinpoints dicloxacillin (469 Da) and (f) flucloxacillin (453 Da) adducts (red arrows). See Supplementary Note 2 for details. All results shown have a Percolator calculated PSM level q ≤ 0.01. Live views for (c) and (d) can be found on Limelight here: (f) https://limelight.yeastrc.org/limelight/go/OX0qQMTOak and here (g) https://limelight.yeastrc.org/limelight/go/x1h2QRXqrE.

Open-mass searching was performed using Magnum and PSMs were imported into Limelight for downstream analysis. Using Limelight, a two-tailed test of proportions comparing untreated with dicloxacillin or flucloxacillin treated HSA, easily picked out exposure specific modifications enriched in the treated samples (Figure 4e-f). Previously published studies determined that dicloxacillin and flucloxacillin produce 469 Da and 453 Da adduct modifications, respectively, on HSA lysine residues^23,29^. These studies relied on MS analysis of unadducted antibiotics, combined with trapping experiments using N-acetyl-cysteine and N-acetyl-lysine plus extensive targeted MS experiments. Using our untargeted method, comparisons of dicloxacillin or flucloxacillin treated versus untreated samples resulted in 469 Da or 453 Da, being the most significantly enriched mass for their respective treatment with no prior knowledge or experimentation. These data show a two-tailed test of proportions comparing treated versus untreated samples, is effective in highlighting exposure specific adducts from PSMs generated by Magnum.

Limelight was designed to support output from any proteomics pipeline, and we performed the same analysis using PSMs from 6 additional open-mass search algorithms (Supplementary Note 4). In all cases Magnum identified the most treatment related PSMs as well as resulting in the largest Z Score for the correct mass compared to other algorithms. Overall, Magnum identified 41 unique dicloxacillin- and 55 unique flucloxacillin-adducted HSA peptides of which 90% contained 1 or more reporter ions resulting from adduct fragmentation (Supplementary Table 7). Manual validation using Limelight’s built-in spectrum viewer further confirmed Magnum’s automated results and example annotated spectra are shown in Figure 5 (dicloxacillin) and Supplementary Figure 12 (flucloxacillin). These data demonstrate our software and workflow can correctly and automatically identify and highlight drug protein adducts and that Magnum is the most sensitive algorithm of those tested.

**Figure 5:**
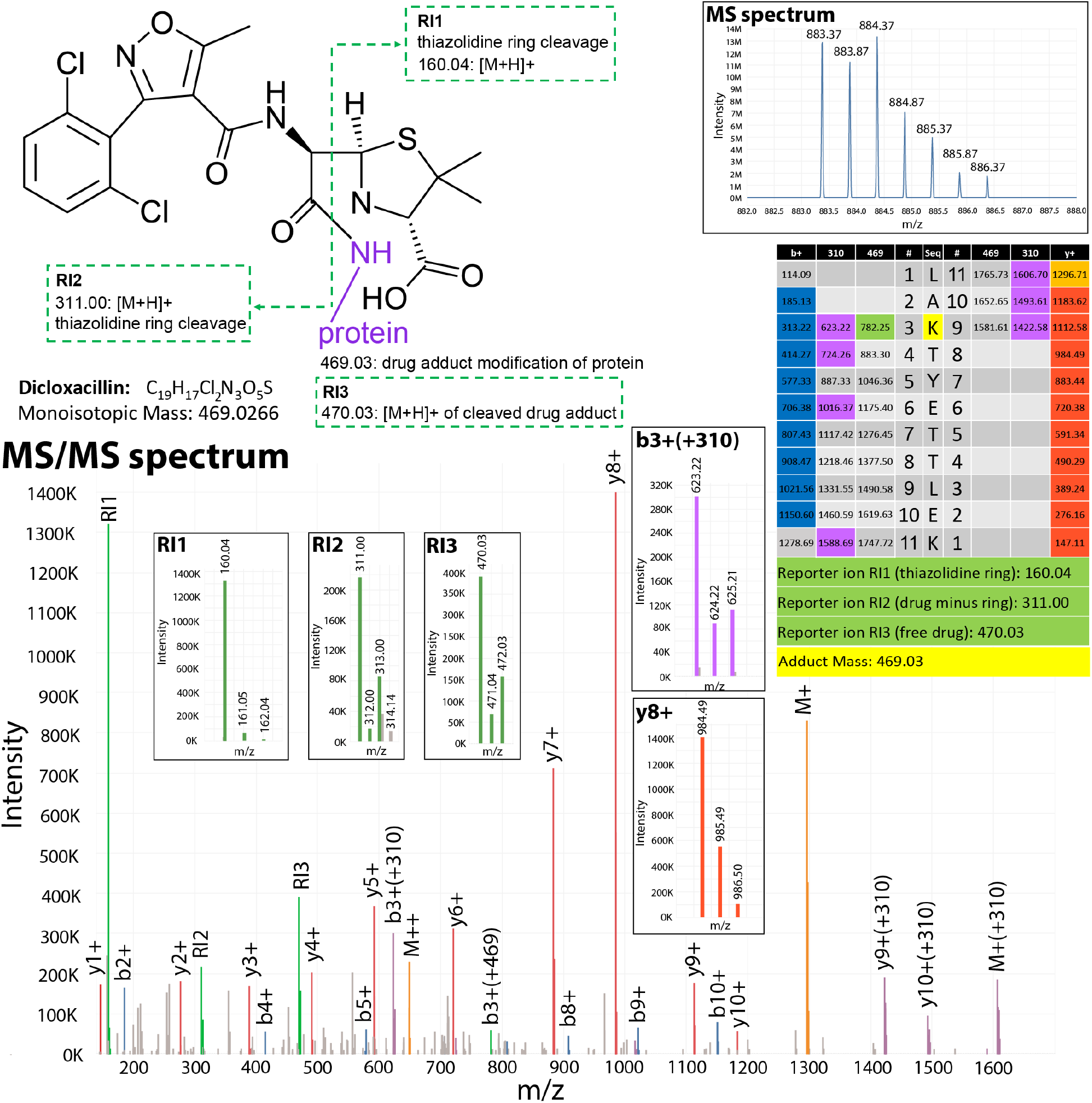
Annotated MS and MS/MS spectrum of dicloxacillin adducted HSA peptide identified by Magnum. The adduct (structure, top left) is covalently bound to the lysine primary amine (purple). Dicloxacillin is an MS-labile adduct. During peptide fragmentation the adduct itself is cleaved at the thiazolidine ring (green dotted line) releasing reporter ion 1 (RI1) and reporter ion 2 (RI2). The entire adduct can also be cleaved from the peptide releasing reporter ion 3 (RI3). The masses of these ions are independent of the peptide to which the drug is adducted as the ions are derived from adduct fragmentation. For this precursor ion, the MS spectrum indicated an observed mass of 883.37 m/z at charge 2 (M+2H^2+^). The observed precursor mass is thus 1,764.724 Da. The theoretical mass of peptide LAKTYETTLEK is 1,295.697 Da. The adduct modification mass, equal to the mass unexplained by the predicted peptide, is thus 469.03 Da. The peptide sequence and calculated ion series are displayed in the table (center right). The annotated MS/MS spectrum is shown below with inset zoomed panels depicting RI1, RI2, RI3, b3(+310) which is the b3+ ion plus the dicloxacillin adduct minus the thiazolidine ring and is the dominant drug modified ion series, and y8+. Note that the MS spectrum, RI2, RI3 and b3+(+310) all have clear chlorine isotope signatures due to the chlorine in dicloxacillin. R1 has no chlorine signature as the thiazolidine ring has no chlorine. Also note that the dominant b and y ion series are unmodified as the adduct is cleaved off the peptide during the fragmentation step. This prevents correct localization of the adduct by all open search algorithms tested (discussed in Supplementary Note 5). The original identification can be viewed on limelight here: https://limelight.yeastrc.org/limelight/d/pg/spectrum-viewer/ps/2327/psm/139200618.

### Magnum and Limelight identify novel raloxifene adducts in CYP3A4 and P450-reductase

Most xenobiotics that ultimately form adducts undergo metabolic activation by cytochrome P450 (CYP) enzymes to form short lived reactive intermediates that are difficult to predict and identify. Raloxifene, a drug commonly used to treat osteoporosis in postmenopausal women, is a mechanism-based inhibitor of CYP3A4^34^. Raloxifene metabolism by CYP3A4 produces several electrophilic species (Supplementary Figure 13), and a single 471 Da adduct on cysteine 239 was previously identified^5^. Past studies identifying adducts in CYP3A4^5–7^ required extensive experimentation involving multiple techniques including: (1) adduct trapping with GSH, N-acetyl cysteine, N-acetyl lysine and potassium cyanide to determine the masses of likely adducts; (2) radiolabeled drugs combined with LC-radiochromatography in-line with MS; (3) whole-protein MS combined with heavy and light labeled drugs; (4) targeted MS experiments using predicted masses of adducts and (5) closed MS/MS database searching using predefined adduct masses determined by methods 1-4.

We exposed CYP3A4 plus P450-reductase to raloxifene *in vitro* in the presence of NADPH. LC-MS/MS data of exposed and untreated samples were acquired, searched using Magnum and analyzed using Limelight. A two-tailed test of proportions identified 471 Da as the most significantly (p = 0) enriched modification mass in raloxifene treated samples (Supplementary Table 9). We searched these data using 6 additional open-mass search algorithms, however none proved as sensitive as Magnum and masses identified as enriched in treated samples by other algorithms were typically also represented by many PSMs in the untreated samples (Supplementary Note 6). For this experiment the advantages of Magnum, which was designed specifically for xenobiotic protein-adduct detection, were required to definitively distinguish raloxifene adducts in a background of other masses.

A fully unrestricted Magnum search (modifications allowed on any amino acid) found 146 PSMs containing a 471 Da modification mass in treated samples versus 14 in untreated samples (10:1 ratio) indicating that >90% of 471 Da identifications made by Magnum in raloxifene treated samples are exposure-specific adducts. Manual evaluation, using Limelight’s built-in spectrum viewer, of all 146 PSMs (Supplementary File raloxifene_Manual_Eval.xlsx) revealed that close to 90% of 471 Da adducts were on cysteine, tryptophan or tyrosine. This is a novel discovery as the single previously identified adduct was on a cysteine and raloxifene adducts have subsequently been presumed to occur on cysteines only. Cysteine, tryptophan and tyrosine contain nucleophilic moieties (the thiol group of cysteine, the aromatic nitrogen of tryptophan and the phenolic group of tyrosine) which can react with electrophilic sites of the reactive raloxifene metabolite to form adducts (Figure 6a, red arrows). We improved the search sensitivity by using Magnum’s ability to restrict open-mass modifications to specific types of residue: searches were performed restricting open-mass modifications to cysteine only, or, to cysteine (C), tryptophan (W), and tyrosine (Y) collectively. Cysteine-restricted searches yielded fewer (67) PSMs containing 471 Da adduct masses confirming the presence of adducts on residues other than cysteine. CWY-restricted searches resulted in more such PSMs in treated samples (157), and less in untreated samples (3), than any other search (a 52:1 ratio, Supplementary Figure 14).

**Figure 6:**
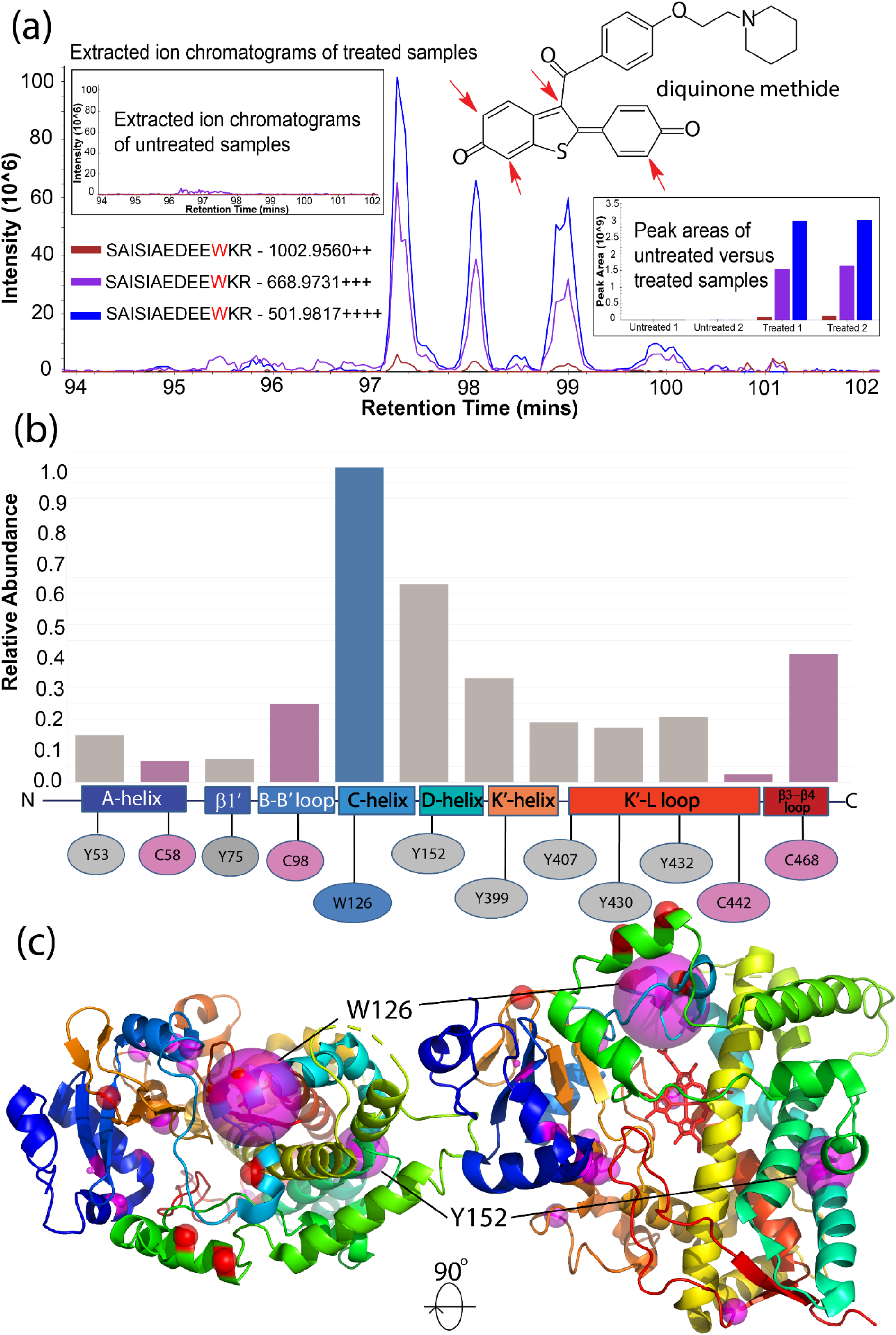
Identification of novel raloxifene adducts in CYP3A4. (a) Extracted ion chromatograms (XICs) of 2+, 3+ and 4+ precursor ions corresponding to CYP3A4 residue W126 (peptide SAISIAEDEEW[471]KR) elute as 4 distinct chromatographic peaks likely representing regioisomers resulting from the different positions^34,37^ in the raloxifene metabolite, diquinone methide, that are subject to nucleophilic attack (red arrows). No signal is observed for these ions in untreated samples (upper left inset). Integrated peak area for these ions shows signal in raloxifene treated samples only (lower left inset). Full skyline sessions showing quantification of all peptides are available on Panorama here: https://panoramaweb.org/CYP3A4-raloxifene.url and integrated peak areas for other peptides are shown in Supplementary Figure 15). (b) Magnum identifies multiple 471 Da protein adducts in CYP3A4 after exposure to raloxifene. Abundance of all modifications is shown based on the number of PSMs identified in each location relative to W126, which was observed in 121 PSMs. Adducted residues in CYP3A4 are mapped to defined regions^38^ of that protein. (c) Observed 471 Da modifications are shown on the structure of CYP3A4 (PDB: 1TQN) as magenta spheres. The size of the magenta spheres is proportional to the relative abundance reported in (b). Red spheres depict CYP3A4 metabolite exit channel 2e (Lys115, Asp123, Glu124, Pro231, Val235 and Lys390)^38^. Supplemental File 1TQN-raloxifene.pse is a pymol session of this image. All results shown have a Percolator calculated PSM level q≤0.01 and were identified by ≥3 PSMs. Raw data are available on Limelight at: https://limelight.yeastrc.org/limelight/go/0UjwIJNz45.

Extracted ion chromatograms (XICs) were produced and quantified in Skyline^35,36^ and showed treatment specific signal for all 471 Da modified peptides identified by Magnum as exclusively in treated samples (Figure 6a, Supplementary Figure 15). We searched an additional 6 untreated and 6 raloxifene treated CYP3A4 samples using CWY-restricted Magnum resulting in a total of 443 PSMs containing a 471 Da modification in CYP3A4 and 91 PSMs in P450-reductase, with just 7 total PSMs showing this mass in the untreated samples (a 76:1 ratio – Supplementary File raloxifene-MagnumCWY.xlsx). These PSMs mapped to 12 distinct amino acids in CYP3A4 (Figure 6b-c; Supplementary Table 12) and 11 in P450-reductase (Supplementary Figure 22) and represent novel raloxifene-protein adducts.

Computational studies have identified several substrate access and metabolite egress channels in CYP3A4^38–40^ but no experimental evidence currently exists to support metabolite egress through these channels after drug oxidation. The most prominent adduct identified in our study, W126, is located in the C-helix of CYP3A4, which flanks the exit channel 2e^38^. W126 adducts support egress of raloxifene metabolites from the CYP3A4 active site via the 2e channel. Interestingly, W126 was associated with 4 distinct chromatographically separated peaks (Figure 6a) that might represent regioisomers resulting from the different positions in the raloxifene metabolite subject to nucleophilic attack (Figure 6a, red arrows)^34,37^. Several other 471 Da modified peptides showed multiple distinguishable chromatographic peaks (Supplementary Note 6) and represent the first evidence for regioisomers in native P450 enzymes.

To a lesser extent raloxifene also forms adducts with the A-helix (Y53, C58) and β3 sheet (Y75) that line the 2a substrate access channel buried in the endoplasmic reticulum membrane, indicating this might be a minor pathway of metabolite egress. Finally, we discovered a cluster of adducts in the K’ helix (Y399), K’-L loop (Y407, Y430, Y432, C442) and D helix (Y152) that are thought to face the cytosol and are thus likely adducted after the raloxifene metabolites have dissociated from CYP3A4.

The identification of multiple novel raloxifene-protein adducts within both CYP3A4 and P450-reductase demonstrates our methodology and software provide greatly expanded power to better understand the structure-function and ligand interactions of catalytic membrane proteins such as cytochrome P450s. Past biochemical studies have typically identified single residues in CYP3A4 that can be modified by xenobiotics despite the expected exposure of multiple nucleophilic residues to reactive metabolites formed by CYP3A4^6,7,34,41^. Additionally, previous studies required extensive experimentation to define adducts while the methods and tools presented here required no prior knowledge or experimentation.

## Discussion

Open-mass search algorithms have become increasingly important in the past decade and can now identify over 50% of spectra invisible to traditional search methods^11^. Previous open-mass search analyses of proteomics datasets have found hundreds of yet-to-be-identified modifications in untreated protein samples^11–14,18,19^, but until now there were few pipelines dedicated to discovering exposure related xenobiotic-protein adducts in a background of modifications identified by open searching. The pipeline presented here allows the use of standard shotgun proteomics to identify xenobiotic-protein adducts in an automated manner with no prior knowledge of the adduct mass or the adducted peptides.

Magnum is optimized to detect adduct modified peptides and outperforms existing tools in xenobiotic-protein adduct discovery. Limelight provides specific tools to automatically highlight exposure specific modifications while also fulfilling a major need in the rapidly developing field of open-mass searching, which until now has had no comprehensive data visualization tools. Users are thus empowered to utilize the best algorithm, or combination of algorithms to fulfill their needs.

The tools and methods developed here are transformative in the fields of drug development and environmental toxicology, which until now have been forced to perform laborious and painstaking manual determination of adduct modification masses followed by targeted MS analysis.

## Methods

### Reagents

Human serum albumin (HSA), flucloxacillin sodium, dicloxacillin sodium, and raloxifene were purchased from MilliporeSigma (Burlington, MA). Rat P450 reductase was expressed and purified as described previously^42^. Purified recombinant human CYP3A4 and the liposome stock containing l- α -Dilauroyl-sn-glycero-3-phosphocholine (DLPC), l- α -diloleoyl-sn-glycero-3-phosphocholine (DOPC), and l-α-dilauroyl-sn-glycero-3-phosphoserine (DLPS) (1:1:1, w/w/w per mL) were gifts from Dr. William Atkins, University of Washington. The expression and purification of human recombinant CYP3A4 was performed as described previously^43^.

### Drug Incubations

Human serum albumin (0.5 mg/mL) was incubated with and without 100 μM antibiotics (flucloxacillin and dicloxacillin) in 100 mM potassium phosphate buffer (pH 7.4) at 37°C for 16 hours. The total volume was 200 μL. The control group contained no antibiotics. After incubation unreacted antibiotics were removed by buffer exchange into 100 mM potassium phosphate buffer (pH7.4) using protein desalting spin columns (ThermoFisher part number 89849) according to manufacturer’s instructions. Samples were then flash-frozen using liquid nitrogen and stored at −80°C until processing for LC-MS analysis. Incubations were performed in duplicates.

Recombinant human CYP3A4 (3 μM) was incubated with 6 μM rat P450-reductase, 2 mM NADPH in the absence and presence of 200 μM raloxifene in buffer containing 20 μg/mL liposomes [DLPC, DOPC, DLPS (1:1:1, w/w/w per mL)], 0.1 mg/mL CHAPS, 3 mM glutathione, 30 MgCl2 and 50 mM potassium HEPES buffer (pH 7.4). The control group contained no raloxifene. Incubations were done in singlet at 37°C for 1 hour in a total volume of 200 μL and the reaction was started by adding NADPH. Samples were flash-frozen using liquid nitrogen right after the incubation and stored at −80°C until LC-MS analysis.

### Mass Spectrometry

#### Sample preparation

β-lactam antibiotics and HSA: Aliquots (30 μL) of control or drug treated HSA (0.5 mg/mL) were reduced by adding 10 mM DTT, final concentration, and incubating at 37°C for 30 minutes. The mixture was brought to a final concentration of 16 mM iodoacetamide and alkylation was performed in the dark by incubating at room temperature for 20 minutes. Trypsin was added at an enzyme to substrate ratio of 1:15 and digestion performed at 37°C for 6 hours in an Eppendorf Thermomixer with shaking (1000 rpm) prior to acidification with 250 mM HCl (final concentration). Samples were spun at maximum speed in a benchtop microfuge for 10 minutes and supernatant was transferred to autosampler vials and stored at −80°C until MS analysis.

Raloxifene and CYP3A4: Aliquots (100 μL) of control or drug treated CYP3A4 incubation reaction mixture (63 μg total protein) were reduced alkylated and digested as described above except samples were digested for 4 hours. To increase peptide coverage, a second set of digests were performed (labeled “extra digest” in results): 100 μL aliquots were incubated at 75°C for 30 minutes. Samples were brought to 5.5 mM TCEP and reduced for 1 hour at 60°C, cooled to room temperature, brought to 6 mM iodoacetamide and alkylated by incubating at room temperature in the dark for 30 min prior to a 16-hour trypsin digestion at 37°C. After digestion solid phase extraction was performed on all CYP3A4 samples using Oasis MCX cartridges (1 cc/30 mg cartridges, Waters corporation, product number 186000782) according to manufacturer’s instructions. The resulting eluate was dried in a speed vac and reconstituted into 100 μL 0.1% trifluoroacetic acid, 2% acetonitrile in water before transfer to autosampler vials and storage at −80°C prior to MS analysis.

#### Chromatography

Sample digest (2 μL ~1 μg) was loaded by autosampler onto a 150 μm Kasil fritted trap packed with 2 cm of ReprosilPur C18AQ (3 μm bead diameter, Dr. Maisch) at a flow rate of 2 μL per min. Desalting was performed with 8 μL of 0.1% formic acid plus 2% acetonitrile and the trap was subsequently brought online with a Self-Packed PicoFrit Column (New Objective part number PF360-75-10-N-5, 75 μm i.d.) packed with 30 cm of ReprosilPur C18AQ (3 μm bead diameter, Dr. Maisch) mounted in an in-house constructed microspray source and placed in line with a Waters Nanoacquity binary UPLC pump plus autosampler. Peptides were eluted from the column at 0.25 μL/min using an acetonitrile gradient. The standard gradient used in all runs except those otherwise specified consisted of the following steps: (1) 0-20 mins; 2-7.5% B; flow 0.25 μL/min; (2) 20-100 mins; 7.5-25% B; flow 0.25 μL/min; (3) 100-140 mins; 25-60% B; flow 0.25 μL/min; (4) 140-145 mins; 60% B; flow 0.25 μL/min; (5) 145-146 mins; 60-95% B; flow 0.25 μL/min; (6) 146-151 mins; 95% B; flow 0.45 μL/min; (7) 151-152 mins; 95-2% B; flow 0.45 μL/min; (8) 152-179 mins; 2% B; flow 0.45 μL/min; (9) 179-180 mins; 2% B; flow 0.25 μL/min. A higher acetonitrile gradient was used for select CYP3A4 samples (labeled “highB” in results) that consisted of the following steps: (1) 0-20 mins; 2-15% B; flow 0.25 μL/min; (2) 20-70 mins; 15-60% B; flow 0.25 μL/min; (3) 70-135 mins; 60-95% B; flow 0.25 μL/min; (4) 135-141 mins; 95% B; flow 0.5 μL/min; (5) 141-142 mins; 95-2% B; flow 0.5 μL/min; (6) 142-164 mins; 2% B; flow 0.5 μL/min; (7) 164-165 mins; 2% B; flow 0.25 μL/min. For all gradients, buffer A was: 0.1% formic acid in water and buffer B was 0.1% formic acid in acetonitrile.

#### Data acquisition

A QExactive HF (Thermo Fisher Scientific) was used to perform mass spectrometry in data dependent mode. A maximum of 20 tandem MS (MS/MS) spectra were acquired per MS spectrum (scan range of *m/z* 400–1,600). The resolution for MS and MS/MS was 60,000 and 15,000, respectively, at *m/z* 200. The automatic gain control targets for MS and MS/MS were set to set to a nominal value of 3e6 and 2e5, respectively, and the maximum fill times were 50 and 100 ms, respectively. MS/MS spectra were acquired using an isolation width of 2.5 *m/z* and a normalized collision energy of 27. MS/MS acquisitions were prevented for +1, >+6 or undefined precursor charge states. Dynamic exclusion was set for 5 s. MS spectra were collected in profile mode and MS/MS spectra were centroided.

#### Data processing

Acquired spectra were converted into mzML (for input to all algorithms except MODa) or mzXML (for input to MODa) using ProteoWizard’s msConvert (Chambers et al., 2012). Proteins present in the samples were identified using Comet^44^ by standard closed searching against the entire human proteome, for HSA samples, or the E. coli proteome, for CYP3A4 samples, plus common contaminants (https://www.thegpm.org/crap/). For CYP3A4 samples protein sequences for the heterologously expressed proteins CYP3A4 and P450-reductase (Supplementary Table 13) were appended to the search database. A q-value was assigned to each PSM through analysis of the target and decoy distributions using Percolator^26^. Smaller databases were made for subsequent open searching consisting only of proteins identified in initial comet searches by at least 3 peptides with a Percolator assigned q-value of ≤ 0.01. Decoy databases consisted of the corresponding set of reversed protein sequences and were provided to algorithms requiring pre-generated decoy sequences. All data reported are filtered at a false discovery rate (FDR) of 1% unless otherwise stated.

### Database searching

#### Comet (closed)

Searches were performed on untreated, flucloxacillin and dicloxacillin treated HSA samples for generation of gold standard results (Supplementary Note 5) and as positive controls in dicloxacillin and flucloxacillin searches by defining the known dicloxacillin or flucloxacillin adduct masses as variable modifications. Comet^44^ version 2019.01 rev. 0 was used for these searches configured with a 15 ppm precursor mass tolerance, a fragment bin tolerance of 0.03, and an isotope error of 3. Dicloxacillin or flucloxacillin adducts on lysine were present as defined variable modifications when specified (469.0266 or 453.0561, respectively). Percolator^26^ version 3.02.1 was used to assign q-values to comet generated PSMs.

#### Comet (500 Da wide precursor)

Searches were performed as for closed searches above, but with a 500 amu precursor mass tolerance and an isotope error of 1. Percolator version 3.02.1 was used to assign q-values to comet generated PSMs.

#### Comet-PTM (open)

Searches were done using Comet version “PTM 2016.01 rev. 2”. Precursor mass tolerance and delta_outer_tolerance were set to 500 amu, delta_inner_tolerance was 0.8 and fragment_bin_tol was 0.02. FDR was calculated by the Limelight XML converter as described below.

#### Magnum (open)

Searches were performed using Magnum version 1.0-dev.11 allowing open modifications of 60-500 Da on all amino acids (adduct_sites = ARNDCQEGHILKMFPOSUTWYV), K only (adduct_sites = K), C only (adduct_sites = C), CWY only (adduct_sites = CWY). Defined variable modifications were oxidation of methionine and carbamidomethylation of cysteine. E value depth was set to 10000. Precursor mass tolerance was 15 ppm and isotope error was set to 1. For dicloxacillin and flucloxacillin HSA searches reporter ions were defined as follows, using a reporter ion threshold of 5: 160.04; 311.00; 470.03; 295.03; 454.06. Percolator version 3.05.0 was used to assign q-values to Magnum generated PSMs.

#### MetaMorpheus (open)

Searches were performed using MetaMorpheus version 0.0.308. An initial calibrate task was performed using a precursor mass tolerance of 15 ppm and a product mass tolerance of 25 ppm. Post calibration searches were performed using precursor and product mass tolerances of 5 and 20 ppm, respectively. The modern search option was specified, and open-mass differences of - 187 and up were allowed. Native q-values calculated by MetaMorpheus were used for FDR filtering.

#### MODa (open)

Searches were done using version 1.62 allowing open modifications of 60-500 Da with a precursor mass tolerance of 15 ppm and a product mass tolerance of 0.03 Da. Variable modifications cannot be defined in MODa. A fixed modification for carbamidomethylation of cysteine was used during analysis of dicloxacillin and flucloxacillin HSA samples as this resulted in increased sensitivity in gold standard searches. For raloxifene searches the native mass of cysteine was used (no fixed carbamidomethylation of cysteine) as raloxifene is known to modify cysteine; instead, open modifications were allowed between 55-500 Da to incorporate the possibility of carbamidomethylation of cysteine. For all searches, data were presented for searches where blind mode was set to 1, allowing 1 open modification per peptide. This was found to be much more sensitive than blind mode 2, which allows an arbitrary number of modifications per peptide. Instrument was set to ESI-TRAP and high resolution was set to on. FDR was calculated by the Limelight XML converter as described below.

#### MSFragger (open)

Searches were performed using version 2.3 allowing open modifications between 60 and 500 Da. Precursor and fragment mass tolerance was set to 15 ppm and calibrate mass was set to 2. Isotope error was 0/1 and localize delta mass was 1. MSFragger output was processed with PeptideProphet^27^ and PTMProphet^45^ (TPP v5.2.1-dev Flammagenitus, Build 202003241419-8041) for error rate estimation and open modification localization using the following options for PeptideProphetParser: ACCMASS DECOYPROBS DECOY=random NONPARAM MASSWIDTH=520 MINPROB=0. PTMProphetParser was run using MASSDIFFMODE MINPROB=0. FDR was calculated by the Limelight XML converter as described below.

#### open-pFind (multinotch)

Searches were done using version EVA.3.0.11 with a precursor mass tolerance of 15 ppm and a fragment tolerance of 0.03 Da. Mixture spectra was true, precursor score model was normal and a threshold of −0.5 was used. Peptide mass was between 600 and 6000 and length between 6 and 100. Open search was set to true. FDR was calculated by the Limelight XML converter as described below.

All searches were performed requiring fully tryptic peptides allowing either 2 or 3 missed cleavages and defining variable modifications of oxidation (15.9949) of methionine and carbamidomethylation (57.021464) of cysteine except MODa which does not allow for defined variable modifications. Search databases consisted of all proteins detectable in the sample by searching spectra against whole-proteome databases using comet. PSMs were processed with Percolator and proteins identified by ≥ 3 PSMs with a Percolator assigned q-value ≤ 0.01 were included in a smaller protein database used for open modification searches. Algorithms that required pre-generated decoys sequences were given the reversed sequence of each target sequence. For each search performed, software version numbers and complete configuration files are available for download via Limelight or via ProteomeXchange (see below).

### False Discovery Rate (FDR) calculation

When a software pipeline produces an FDR or q-value associated with a PSM, that value was used to filter the data at a 0.01 FDR or q-value threshold directly. The software pipelines below do not produce a PSM-level score analogous to a FDR or q-value. This value was therefore calculated and associated with every PSM as follows.

#### MSFragger (post processed by TPP)

This pipeline produces a probability score associated with each PSM that represents the probability that the identified peptide is correctly associated with the given spectrum. So, to calculate the predicted FDR an estimation for the number of incorrectly identified PSMs can be calculated as the sum of 1 minus this probability for all PSMs with a given probability or better. This sum can be divided by the total number of PSMs with a given probability or better to obtain an estimate of the FDR. The code may be viewed on GitHub at https://github.com/yeastrc/limelight-import-msfragger-tpp.

#### Comet-PTM

All unique E-value scores (reported by Comet-PTM) are sorted from best to worst. This list is then iterated over and for each score a sum is calculated for the total number of target hits and the total number of decoy hits with that score or better. An FDR is calculated for the given score as the total number of decoy hits divided by the total number of decoy hits plus total number of target hits. Then the FDR for any previously processed score (i.e., better scores) is changed to the minimum of its existing FDR or the current score’s FDR. The code may be viewed on GitHub at https://github.com/yeastrc/limelight-import-cometptm.

#### MODa

All unique probability scores (reported by MODa) are sorted from best to worst. This list is then iterated over and for each score a sum is calculated for the total number of target hits and the total number of decoy hits with that score or better. An FDR is calculated for the given score as the total number of decoy hits divided by the total number of decoy hits plus total number of target hits. Then the FDR for any previously processed score (i.e., better scores) is changed to the minimum of its existing FDR or the current score’s FDR. The code may be viewed on GitHub at https://github.com/yeastrc/limelight-import-moda.

#### Open-pFind

All unique final scores (reported by open-pFind) are sorted from best to worst. This list is then iterated over and for each score a sum is calculated for the total number of target hits and the total number of decoy hits with that score or better. An FDR is calculated for the given score as the total number of decoy hits divided by the total number of decoy hits plus total number of target hits. Then the FDR for any previously processed score (i.e., better scores) is changed to the minimum of its existing FDR or the current score’s FDR. The code may be viewed on GitHub at https://github.com/yeastrc/limelight-import-open-pfind

### Limelight XML generation

Native search results from all search pipelines were converted to Limelight XML prior to uploading to the Limelight web application. The authors of Limelight have written converters for many popular software pipelines including Comet^44^, Percolator^26^, the Trans-Proteomic Pipeline (TPP)^46^, Crux^47^, MSFragger^12^, open-pFind^13^, Comet-PTM^15^, MetaMorpheus^18^, MODa^14^, TagGraph^19^ and Magnum (this paper). A current list of converters is available at our documentation website here: https://limelight-ms.readthedocs.io/. The Limelight XML schema may be seen at https://github.com/yeastrc/limelight-import-api/tree/master/xsd.

### Quantification of CYP3A4 and P450-reductase adducts using Skyline

Peptides were quantified by integrating and summing the area under the curve of the peptide elution for specific transitions using Skyline^35,36^, which automatically extracts and quantifies MS1 and MS2 ion chromatograms. All MS1 transitions corresponding to residues identified in initial Magnum-CWY searches as having a 471 Da modification in ≥ 2 PSMs (Supplementary Table 10) were quantified as previously described^36^. Additionally, 5 unmodified CYP3A4 and 5 unmodified P450-reductase peptides were chosen for normalization of the data. These contained no raloxifene modifiable residues (e.g. C, W, Y) and no tryptic ragged ends (e.g. C-terminal KK, RR, KR or RK). The total extracted MS1 transition area of each normalization peptides within each experiment was summed. For each 471 Da modified transition the total MS1 area for that transition was divided by the normalization peptide sum for that experiment and this normalized ratio was taken as the value for each 471 Da modified transition. Full data plus Limelight links can be found in Supplementary File raloxifene-MagnumCWY.xlsx (InitialReplicates tab). Full Skyline sessions are available on Panorama here: https://panoramaweb.org/CYP3A4-raloxifene.url.

## Code availability

Magnum is written in C++, supports multithreaded computation, is open source and freely available at http://magnum-ms.org. Full source code plus precompiled Magnum binaries are available for Windows and Linux. Docker containers are additionally provided and support all operating systems. Magnum Docker containers include Percolator^26^ to assign q-values to Magnum generated PSMs, plus a Magnum/Percolator to Limelight XML converter for convenience. Docker containers plus associated documentation can be found here: https://limelight-ms.readthedocs.io/en/latest/tutorials/magnum-pipeline.html. Magnum outputs results a simple tab-delimited text format. Output can also be exported in PepXML^48^ format, for potential use with existing software supporting this format. For further details of Magnum see Supplementary Note 1.

Limelight is open source and is freely available at https://limelight-ms.org/. Limelight is written in Java, Typescript, HTML and CSS and makes use of the MySQL relational database management system. In addition to all Limelight source code, preconfigured Docker containers are provided for running Limelight and Limelight XML converters. See the installation guide at the documentation website here: https://limelight-ms.readthedocs.io/. Users not able to run their own Limelight installation may email the authors at with a description of their project to request access to the authors’ installation. For full details of Limelight see Supplementary Note 2.

## Data availability

All raw and processed data discussed in this paper are available via Limelight at: https://limelight.yeastrc.org/limelight/p/adduct-discovery. In addition, complete search algorithm configuration files, fasta search databases, raw search output and raw MS data files were deposited to the ProteomeXchange Consortium via the PRIDE^49^ partner repository with the dataset identifier PXD025019. Full Skyline quantification data of CYP3A4/raloxifene experiments were deposited to the ProteomeXchange Consortium via Panorama Public^50^ and is available at: https://panoramaweb.org/CYP3A4-raloxifene.url with the dataset identifier PXD024932. Data can also be requested directly from the corresponding author.

## Acknowledgements

We thank Jimmy Eng and David D. Shteynberg for help with comet-PTM and PTMProphet, respectively. Purified recombinant human CYP3A4 and the liposome stock were gifts from Dr. William Atkins. Research reported in this publication was supported by the National Institutes of Health, National Institute of General Medical Sciences under Award P41GM103533 (to M.J.M.), and National Institute On Aging under Award Number U19AG023122 (to R.L.M.).

## Author Contributions

A.Z., G.Z., M.R., M.R.H. and N.I. conceived the experiments. G.Z. performed drug-protein incubations. A.Z. carried out the MS experiments and MS data analysis. M.R.H. developed Magnum. M.R. and D.J. developed Limelight. The manuscript was written by A.Z. with contributions from M.R., M.R.H., NI and G.Z. All authors discussed the results and commented on the manuscript.

## References

1. Misra, B. B. The Chemical Exposome of Human Aging. Frontiers in Genetics vol. 11 (2020).

2. Gan, J., Zhang, H. & Humphreys, W. G. Drug-Protein Adducts: Chemistry, Mechanisms of Toxicity, and Methods of Characterization. Chem. Res. Toxicol. 29, 2040–2057 (2016).

3. Whitby, L. R., Obach, R. S., Simon, G. M., Hayward, M. M. & Cravatt, B. F. Quantitative Chemical Proteomic Profiling of the in Vivo Targets of Reactive Drug Metabolites. ACS Chem. Biol. 12, 2040–2050 (2017).

4. Evans, D. C., Watt, A. P., Nicoll-Griffith, D. A. & Baillie, T. A. Drug-Protein Adducts: An Industry Perspective on Minimizing the Potential for Drug Bioactivation in Drug Discovery and Development. Chem. Res. Toxicol. 17, 3–16 (2004).

5. Baer, B. R., Wienkers, L. C. & Rock, D. A. Time-dependent inactivation of P450 3A4 by raloxifene: Identification of Cys239 as the site of apoprotein alkylation. Chem. Res. Toxicol. 20, 954–964 (2007).

6. Lin, H. L., Kenaan, C. & Hollenberg, P. F. Identification of the residue in human CYP3A4 that is covalently modified by bergamottin and the reactive intermediate that contributes to the grapefruit juice effect. Drug Metab. Dispos. 40, 998–1006 (2012).

7. Rock, B. M., Hengel, S. M., Rock, D. A., Wienkers, L. C. & Kunze, K. L. Characterization of ritonavir-mediated inactivation of cytochrome P450 3A4. Mol. Pharmacol. 86, 665–674 (2014).

8. Törnqvist, M. et al. Protein adducts: Quantitative and qualitative aspects of their formation, analysis and applications. J. Chromatogr. B Anal. Technol. Biomed. Life Sci. 778, 279–308 (2002).

9. Sabbioni, G. & Turesky, R. J. Biomonitoring human albumin adducts: The past, the present, and the future. Chem. Res. Toxicol. 30, 332–366 (2017).

10. Eng, J. K., Searle, B. C., Clauser, K. R. & Tabb, D. L. A Face in the Crowd: Recognizing Peptides Through Database Search. Mol. Cell. Proteomics 10, 1–9 (2011).

11. Chick, J. M. et al. A mass-tolerant database search identifies a large proportion of unassigned spectra in shotgun proteomics as modified peptides. Nat. Biotechnol. 33, 743–749 (2015).

12. Kong, A. T., Leprevost, F. V., Avtonomov, D. M., Mellacheruvu, D. & Nesvizhskii, A. I. MSFragger: Ultrafast and comprehensive peptide identification in mass spectrometry-based proteomics. Nat. Methods 14, 513–520 (2017).

13. Chi, H. et al. Comprehensive identification of peptides in tandem mass spectra using an efficient open search engine. Nat. Biotechnol. 36, 1059–1066 (2018).

14. Na, S., Bandeira, N. & Paek, E. Fast Multi-blind Modification Search through Tandem Mass Spectrometry. Mol. Cell. Proteomics 11, M111.010199 (2012).

15. Bagwan, N. et al. Comprehensive Quantification of the Modified Proteome Reveals Oxidative Heart Damage in Mitochondrial Heteroplasmy. Cell Rep. 23, 3685–3697 (2018).

16. Li, Q. et al. Global Post-Translational Modification Discovery. J. Proteome Res. 16, 1383–1390 (2017).

17. David, M., Fertin, G., Rogniaux, H. & Tessier, D. SpecOMS: A Full Open Modification Search Method Performing All-to-All Spectra Comparisons within Minutes. J. Proteome Res. 16, 3030–3038 (2017).

18. Solntsev, S. K., Shortreed, M. R., Frey, B. L. & Smith, L. M. Enhanced Global Post-translational Modification Discovery with MetaMorpheus. J. Proteome Res. 17, 1844–1851 (2018).

19. Devabhaktuni, A. et al. TagGraph reveals vast protein modification landscapes from large tandem mass spectrometry datasets. Nat. Biotechnol. 37, 469–479 (2019).

20. Hoopmann, M. R. et al. Kojak: Efficient analysis of chemically cross-linked protein complexes. J. Proteome Res. 14, 2190–2198 (2015).

21. Noble, W. S. Mass spectrometrists should search only for peptides they care about. Nat. Methods 12, 605–608 (2015).

22. Enoch, S. J., Ellison, C. M., Schultz, T. W. & Cronin, M. T. D. A review of the electrophilic reaction chemistry involved in covalent protein binding relevant to toxicity. Critical Reviews in Toxicology vol. 41 783–802 (2011).

23. Jenkins, R. E. et al. Characterisation of flucloxacillin and 5-hydroxymethyl flucloxacillin haptenated HSA in vitro and in vivo. Proteomics - Clin. Appl. 3, 720–729 (2009).

24. Ariza, A. et al. Protein haptenation by amoxicillin: High resolution mass spectrometry analysis and identification of target proteins in serum. J. Proteomics 77, 504–520 (2012).

25. Paul Zolg, D. et al. Proteometools: Systematic characterization of 21 post-translational protein modifications by liquid chromatography tandem mass spectrometry (lc-ms/ms) using synthetic peptides. Mol. Cell. Proteomics 17, 1850–1863 (2018).

26. Kall, L. et al. Semi-supervised learning for peptide identification from shotgun proteomics datasets. Nat. Methods 4, 923–925 (2007).

27. Shteynberg, D. et al. iProphet: Multi-level Integrative Analysis of Shotgun Proteomic Data Improves Peptide and Protein Identification Rates and Error Estimates. Mol. Cell. Proteomics 10, 1–16 (2011).

28. Rappaport, S. M., Li, H., Grigoryan, H., Funk, W. E. & Williams, E. R. Adductomics: Characterizing exposures to reactive electrophiles. Toxicol. Lett. 213, 83–90 (2012).

29. Parker, C. E., Perkins, J. R. & Tomer, K. B. Nanoscale packed capillary liquid chromatography-electrospray ionization mass spectrometry: analysis of penicillins and cephems. 616, 45–51 (1993).

30. Ariza, A. et al. Hypersensitivity reactions to ß-lactams: Relevance of hapten-protein conjugates. J. Investig. Allergol. Clin. Immunol. 25, 12–25 (2015).

31. Rad, R. et al. Improved Monoisotopic Mass Estimation for Deeper Proteome Coverage. J. Proteome Res. (2020) doi:10.1021/acs.jproteome.0c00563.

32. Creasy, D. M. & Cottrell, J. S. Unimod: Protein modifications for mass spectrometry. Proteomics vol. 4 1534–1536 (2004).

33. Sharma, V., Eng, J. K., Maccoss, M. J. & Riffle, M. A mass spectrometry proteomics data management platform. Mol Cell Proteomics 11, 824–831 (2012).

34. Chen, Q. et al. Cytochrome P450 3A4-mediated bioactivation of raloxifene: Irreversible enzyme inhibition and thiol adduct formation. Chem. Res. Toxicol. 15, 907–914 (2002).

35. MacLean, B. et al. Skyline: An open source document editor for creating and analyzing targeted proteomics experiments. Bioinformatics 26, 966–968 (2010).

36. Schilling, B. et al. Platform-independent and label-free quantitation of proteomic data using MS1 extracted ion chromatograms in skyline: Application to protein acetylation and phosphorylation. Mol. Cell. Proteomics 11, 202–214 (2012).

37. Yu, L. et al. Oxidation of raloxifene to quinoids: Potential toxic pathways via a diquinone methide and o-quinones. Chem. Res. Toxicol. 17, 879–888 (2004).

38. Benkaidali, L. et al. Four Major Channels Detected in the Cytochrome P450 3A4: A Step toward Understanding Its Multispecificity. Int. J. Mol. Sci. 20, 987 (2019).

39. Cojocaru, V., Winn, P. J. & Wade, R. C. The ins and outs of cytochrome P450s. Biochimica et Biophysica Acta - General Subjects vol. 1770 390–401 (2007).

40. VandenBrink, B. M. et al. Cytochrome P450 architecture and cysteine nucleophile placement impact raloxifene-mediated mechanism-based inactivation. Mol. Pharmacol. 82, 835–842 (2012).

41. Wen, B. et al. Cysteine 98 in CYP3A4 contributes to conformational integrity required for P450 interaction with CYP reductase. Arch. Biochem. Biophys. 454, 42–54 (2006).

42. Woods, C. M., Fernandez, C., Kunze, K. L. & Atkins, W. M. Allosteric activation of cytochrome P450 3A4 by α-naphthoflavone: Branch point regulation revealed by isotope dilution analysis. Biochemistry 50, 10041–10051 (2011).

43. Redhair, M., Hackett, J. C., Pelletier, R. D. & Atkins, W. M. Dynamics and Location of the Allosteric Midazolam Site in Cytochrome P4503A4 in Lipid Nanodiscs. Biochemistry 59, 766–779 (2020).

44. Eng, J. K., Jahan, T. A. & Hoopmann, M. R. Comet: an open-source MS/MS sequence database search tool. Proteomics 13, 22–24 (2013).

45. Shteynberg, D. D. et al. PTMProphet: Fast and Accurate Mass Modification Localization for the Trans-Proteomic Pipeline. J. Proteome Res. 18, 4262–4272 (2019).

46. Deutsch, E. W. et al. A guided tour of the Trans-Proteomic Pipeline. Proteomics vol. 10 1150–1159 (2010).

47. McIlwain, S. et al. Crux: Rapid open source protein tandem mass spectrometry analysis. J. Proteome Res. 13, 4488–4491 (2014).

48. Keller, A., Eng, J., Zhang, N., Li, X. jun & Aebersold, R. A uniform proteomics MS/MS analysis platform utilizing open XML file formats. Mol. Syst. Biol. 1, (2005).

49. Vizcaíno, J. A. et al. A guide to the Proteomics Identifications Database proteomics data repository. Proteomics vol. 9 4276–4283 (2009).

50. Sharma, V. et al. Panorama Public: A Public Repository for Quantitative Data Sets Processed in Skyline. Mol. Cell. Proteomics 17, 1239–1244 (2018).

