## Supplemental Material for "Discovery and visualization of uncharacterized drug-protein adducts using mass spectrometry"

Description 18
Experiment Builder 18
*Supplementary Figure 7: Iteratively building an experiment using the experiment builder.* 19
Score filters and cutoffs 19
Single protein view 19
Peptide view 20
Protein view 20
Modification view 20
*Visualizations and Transformations* 21
*Two-tailed test of proportions* 21
Supplementary Note 4: Development and validation of adduct discovery pipeline using dicloxacillin, flucloxacillin and HSA 23
Open modifications unrelated to exposure 23
*Supplementary Figure 8: Open modification masses in untreated, purified human serum albumin as identified by 7 different open search algorithms and visualized by Limelight.* 23
*Supplementary Figure 9: The number of PSMs at 1% FDR resulting from searching 6 untreated and dicloxacillin and flucloxacillin exposed HSA datasets with open and closed search algorithms.* 24
Discovery of dicloxacillin/flucloxacillin adducts in HSA 24
*Supplementary Figure 10: Open modification masses identified by Magnum or MSFragger in untreated, dicloxacillin treated or flucloxacillin treated human serum albumin (HSA).* 25
*Supplementary Figure 11: A two-tailed test of proportions performed within Limelight identifies treatment specific adducts in HSA.* 26
*Supplementary Table 7: Dicloxacillin and flucloxacillin adducted peptides identified by Magnum.* 27
*Supplementary Figure 12: Annotated MS and MS/MS spectrum of flucloxacillin adducted HSA peptide identified by Magnum.* 30
Supplementary Note 5: Localization of dicloxacillin and flucloxacillin adducts 31
*Supplementary Table 8: Dicloxacillin and flucloxacillin adduct localization determined by Magnum or MSFragger plus PTMProphet in 2,979 gold standard MS/MS spectra.* 31
Supplementary Note 6: CYP3A4/Raloxifene analysis 32
*Supplementary Figure 13: Bioactivation of raloxifene by CYP3A4 and representative adducts formed with cysteine, tryptophan or tyrosine.* 32
Initial Experiments and Skyline Quantification 32
*Supplementary Table 9: A two-tailed test of proportions identifies a 471 Da raloxifene specific adduct mass in CYP3A4 and P450-reductase.* 33
*Supplementary Figure 14: Identification of novel raloxifene adducts in CYP3A4 and P450-reductase* 34

### Supplementary Note 1: Magnum

#### Overview

Magnum is a database search algorithm designed to identify adducts of variable mass without *a priori* knowledge. The algorithm has three major functional categories: reading and processing inputs, open modification database searching, and results scoring (main manuscript Figure 1). Output for Magnum is a simple tab-delimited text format. The output is optionally also exported in PepXML[1] format, for potential use with existing software supporting this format. Magnum is written in C++, and is open source and freely available from http://magnum-ms.org/.

#### Reading and Processing Inputs

Magnum requires a protein FASTA sequence file and a spectral data file. The FASTA sequence file is parsed to create a peptide list according to the user-defined proteolytic cleavage rules (see Supplementary Table 1). Peptides containing suspected adduct attachment sites are marked to facilitate downstream analysis of these sequences. The user can also alter the mass of any amino acid, specify novel characters for special-case amino acids, and identify both static and variable modification masses on either amino acids, peptide termini, or protein termini. Decoy protein sequences can be identified with a user-defined text label, and peptides mapping to these protein sequences will be identified as decoy PSMs in the Magnum results, for the purposes of downstream validation at the user's discretion.

*Supplementary Table 1: Select parameters relevant to open modification searches in Magnum*

| Open Modification Search Parameters | | |
|---|---|---|
| Parameter | Values | Description |
| adduct_sites | Uppercase [A-Z],n,c | Identifies one or more sites of open modifications. Must be uppercase letters, except for lowercase 'c' and 'n' to indicate protein C-terminus and protein N-terminus. |
| min_adduct_mass | number | Describes the smallest adduct mass in the open modification search range. |
| max_adduct_mass | number > min_adduct_mass | Describes the largest adduct mass in the open modification search range. |
| **Peptide Sequence Search Rules** | | |
| Parameter | Values | Description |
| enzyme | enzyme cleavage rules | Specifies the peptide sequence cleavage rules to produce peptides. Rules are defined as amino acids where cleavage occurs. See http://magnum-ms.org/param/enzyme.html |
| max_miscleavages | positive number | Maximum number of missed enzyme cleavages to consider when computing the peptide list from the FASTA sequence file. |
| min_peptide_mass | number | Smallest peptide mass allowed in the search space (including the adduct mass). |
| max_peptide_mass | number > min_peptide_mass | Largest peptide mass allowed in the search space (including the adduct mass). |
| min_peptide_length | positive number | Minimum number of amino acids (regardless of mass) in any peptide to search. |
| max_peptide_length | number > min_peptide_length | Maximum number of amino acids (regardless of mass) in any peptide to search. |

#### Select Parameters to Customize Search Results

| Parameter | Values | Description |
|---|---|---|
| modification | amino acid, modification mass | Indicates site and mass for variable modifications of known mass. This parameter can be repeated any number of times, each specifying a novel variable modification to search. |
| max_mods_per_peptide | positive number | Maximum number of variable modifications to consider on a peptide. The larger the number, the slower Magnum performs. |
| fixed_modification | amino acid, modification mass | Indicates fixed amino acid modification mass applied to all instances of the amino acid. Example: "C 57.02146" for carbamidomethyl-cysteine. |
| isotope_error | positive number | Integer value indicating number of carbon atom offsets to consider when evaluating precursor mass predictions. |
| e_value_depth | positive number | Number of decoy peptides per histogram when computing e-values for each PSM. Recommended to have at least 5000. However, larger numbers increase computation time for Magnum. |

The spectral data input file must be one of several supported open formats that include mzML (preferred), mzXML, and MGF. Magnum reads the spectral data file to extract MS/MS scan data, performs refinement steps, and converts the spectra to an internal data structure for rapid cross-correlation analysis[2,3] before storing all spectral data in memory. Refinement consists of two major processes. The first process performs analysis of precursor MS spectra to refine the precursor mass. Precursor refinement attempts to find the elution apex of the selected peptide represented in an MS/MS spectrum, to more accurately predict the monoisotopic precursor mass and charge state, particularly if such information is not provided in the spectral data file. Precursor refinement is skipped if the data contain no precursor MS spectra. Additional functions allow for estimation of isotope mass errors and additional charge state assignments among ambiguous or missing precursor information. For many spectra analyzed by Magnum, more than one candidate precursor mass and charge state may be assigned for database searching. The second major refinement process consists of MS/MS peak refinement. Here, isotope clusters are collapsed to their monoisotopic peak, summing the intensities of each peak in the cluster. Subsequent fragment ion matching (see below) is therefore performed on the monoisotopic mass. Additional, optional processing includes reducing the MS/MS spectra to a fixed, user-defined maximum number of peaks. These steps are repeated on all MS/MS spectra before proceeding to the database searching procedures.

##### Open Modification Database Searching

Magnum attempts to identify peptide sequences from MS/MS spectra allowing for an open modification mass with a user-defined range, referred to as an adduct mass. Database searching is performed by matching theoretical fragment ions (*a*, *b*, *c*, *x*, *y*, or *z*, user-defined) for every peptide sequence parsed from the FASTA file to every spectrum for which the peptide mass falls within the precursor mass tolerance. Which spectra fall within this mass range is defined as:

Supplementary Equation 1: $p + v + a_{min} \leq s_{pre} \leq p + v + a_{max}$

Where $p$ is the peptide mass, $v$ is the sum of the variable modification masses (if any), and $a_{min}$ and $a_{max}$ are the smallest and largest adduct masses defined by the user. $s_{pre}$ is a precursor mass assigned to an MS/MS spectrum. For peptides without an adduct binding site, $a_{min} = a_{max} = 0$, defining a narrow mass range with a user-defined ppm tolerance around the precursor mass for spectra to search. For peptides with an adduct site, the mass range may span hundreds of daltons and require searching several thousand spectra. The adduct mass is different for each spectrum, and defined as:

Supplementary Equation 2: $a_{pre} = s_{pre} - p - v$

Where $a$ is the adduct mass for a predicted precursor (*pre*) of spectrum $s$, and $p$ and $v$ are the peptide and variable modification masses, respectively. If the peptide contains more than one adduct site, the adduct mass is iteratively scored at each site and the highest scoring orientation is kept. In this manner, it is possible to localize the adduct on the peptide, however, no probability is assigned. Thus, the localization is simply the highest scoring orientation without validating the likeliness that this orientation is correct given all available options. The adduct mass is never divided among multiple sites, and therefore only a single adduct of variable mass is ever scored per peptide.

#### Results Scoring

Peptide-spectrum matches (PSMs) are initially scored using a cross-correlation scoring method (Xcorr). However, Xcorr values are not generally comparable between PSMs, as the Xcorr values for longer peptide sequences tend to be higher than Xcorr values for shorter peptide sequences. This is because longer peptide sequences contain more theoretical fragment ion masses to match to a MS/MS spectrum, giving them a higher potential score. A solution to this problem is to compute an expect value (e-value) for the PSM with the highest Xcorr value for each spectrum[4], using the histogram of all PSM Xcorr values to that spectrum. This generally works well if the assumption that all PSMs for a given spectrum are approximately the same length. However, for open modification searches, the PSMs for a spectrum have a much larger range of peptide lengths when considering a short peptide with a large adduct vs. a long peptide without any adduct. Therefore, an alternative method was used to compute e-values for all PSMs for a spectrum (not just the highest Xcorr value), then re-rank the PSMs by e-value and return the PSM with the lowest e-value to the user.

To more accurately compute an e-value from an Xcorr value for a PSM of a given length, a histogram of Xcorr values of random peptide sequences of equal length was generated for each spectrum. Thus, the e-value for a short peptide of 10 amino acids with a large adduct was generated using a histogram of Xcorr values from random peptides of 10 amino acids in length. For the same spectrum, the e-value for a different peptide of 14 amino acids with a small adduct is then computed using a histogram of Xcorr values from random peptides of 14 amino acids in length. By using this approach, the 10-amino acid PSM may produce a lower e-value than the 14-amino acid PSM, despite having a lower Xcorr value. The e-values normalize the effects of peptide length represented by Xcorr values, allowing peptides of very disparate lengths to be compared for the same spectrum. This step of the results scoring is very computationally intensive, which is mitigated by pre-computing the histograms prior to the database search. The pre-computation process is made efficient by taking all the histogram Xcorr values for peptides of length *n*, and extending them by one additional fragment ion to produce histograms of length *n+1*. This process is repeated for the range of all expected peptide lengths for a given MS/MS spectrum after considering all possible adduct sizes.

#### MS-labile versus non-labile adducts

MS-labile modifications are prone to dissociation during peptide fragmentation preceding MS/MS acquisition and may result in a strong unmodified ion series. In contrast, non-labile adducts remain attached to the peptide through the fragmentation process and result in an adduct-modified ion series. To optimally account for both situations Magnum calculates an unmodified ion series as well as a modified ion series localizing the open modification to the position being tested. These are scored independently, and the highest scoring result kept. Peptide sequence and adduct mass identification is therefore possible with or without adduct localization.

#### Restriction of open modification mass to specific residues in Magnum

Magnum optionally allows adduct localization to be restricted to specific amino acids. The reactivity of xenobiotics may be known or hypothesized based on the chemistry of the compound of interest, or detection of glutathione or other conjugates to reactive intermediates. Increased search sensitivity and statistical power can be gained by restricting the open modification search space to specific amino acids[5].

#### Adduct reporter ions

We implemented the ability of Magnum to flag peptide spectrum matches (PSMs) that contained user-defined reporter ion masses and of Limelight to filter data using this information and to annotate reporter ions when viewing individual spectra using Lorikeet[6]. This is useful to identify MS/MS spectra that contain peptides modified with labile adducts whose fragment masses will be independent of the peptides to which they are adducted. Details of this parameter can be found on the Magnum website here.

### Supplementary Note 2: Gold standard dataset for evaluation of xenobiotic-protein adduct discovery

As the focus of the current study was to accurately detect unknown xenobiotic-protein adducts related to specific exposures, the assignment of correct open modification masses is critical. We therefore created three gold standard datasets to allow evaluation of both the accuracy (precision) and sensitivity (recall) of, Magnum, the search algorithm presented here, as well as previously published open search algorithms, within the context of our xenobiotic-protein adduct discovery pipeline. These datasets were derived from four dicloxacillin and flucloxacillin treated HSA samples and the raw MS data files (Supplementary Table 2) were deposited to the ProteomeXchange Consortium via the PRIDE[7] partner repository with the dataset identifier PXD025019. These data consist of 307,652 MS/MS scans. We derived known correct open modification masses for 2,979 unique MS/MS spectra from these data using two methods.

*Supplementary Table 2: Raw files used for creation of gold standard data.*

| Sample Treatment | Untreated (rep 1) | Untreated (rep 2) | Flucloxacillin (rep 1) | Flucloxacillin (rep 2) | Dicloxacillin (rep 1) | Dicloxacillin (rep 2) |
|---|---|---|---|---|---|---|
| Filename | QEP2_2018_0812_AZ_024_az732_AZ.mzML | QEP2_2018_0812_AZ_025_az733_AZ.mzML | QEP2_2018_0812_AZ_028_az734_AZ.mzML | QEP2_2018_0812_AZ_029_az735_AZ.mzML | QEP2_2018_0812_AZ_029_az735_AZ.mzML | QEP2_2018_0812_AZ_033_az736_AZ.mzML |
| Used in gold standard dataset | no | no | yes | yes | yes | yes |

Both methods made use of the fact that β-lactam antibiotics and their adducts fragment in MS/MS giving rise to known reporter ions[8,9] (Supplementary Figure 1, green boxes).

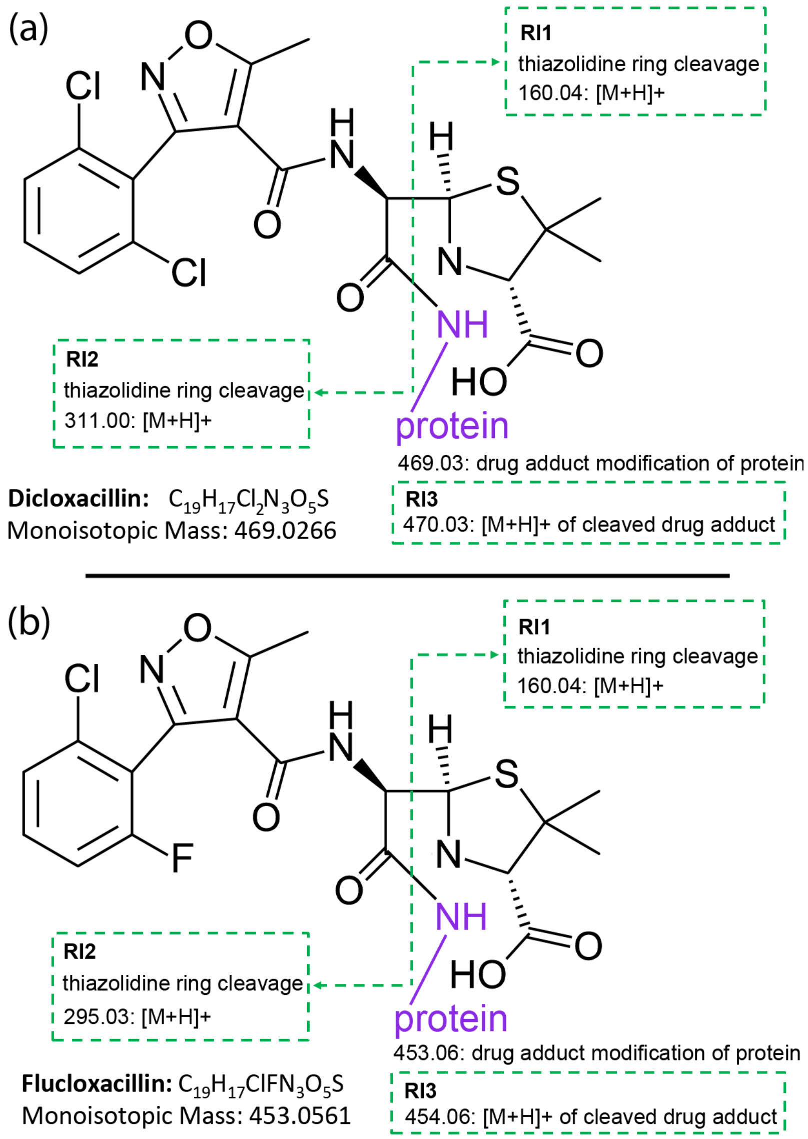

*Supplementary Figure 1: The structure of (a) dicloxacillin and (b) flucloxacillin after forming adducts on a lysine primary amine. The monoisotopic mass and chemical formular of the original antibiotics are listed. For both antibiotics, the adduct is covalently bound to the lysine primary amine (purple). Dicloxacillin and flucloxacillin form MS-labile adducts. During peptide fragmentation the adduct itself is cleaved at the thiazolidine ring (green dotted line) releasing reporter ion 1 (RI1) and reporter ion 2 (RI2). The entire adduct can also be cleaved from the peptide releasing reporter ion 3 (RI3). The masses of these ions are independent of the peptide to which the drug is adducted as the ions are derived from adduct fragmentation. The adduct masses and those of the reporter ions have been previously characterized[8,9].*

These reporter ions are present within any MS/MS spectrum that contains a β-lactam antibiotic adduct, and their masses are unrelated to the peptide to which the adduct is attached. We wrote a simple program called ScanFinder, available at http://magnum-ms.org, which searches raw spectra for signals at a defined m/z and intensity (percent of the base peak). Raw MS data acquired from untreated, flucloxacillin and dicloxacillin treated HSA were searched for both dicloxacillin and flucloxacillin reporter ions at the specified intensities. The results of these searches are summarized in Supplementary Table 3 and show that zero spectra from untreated HSA samples contained the combination of reporter ions searched for at the required intensities. Likewise, zero spectra in flucloxacillin treated samples contained dicloxacillin reporter ions and zero dicloxacillin treated samples contained flucloxcillin reporter ions at the specified intensities. This is important as it shows that spectra picked out in the treated samples are specific to their respective treatments.

*Supplementary Table 3: Number of MS/MS spectra that contain β-lactam antibiotic reporter ions at the defined m/z and intensities.*

| Sample Treatment | Untreated (rep 1) | Untreated (rep 2) | Flucloxacillin (rep 1) | Flucloxacillin (rep 2) | Dicloxacillin (rep 1) | Dicloxacillin (rep 2) |
|---|---|---|---|---|---|---|
| Number of MS/MS spectra containing dicloxacillin reporter ions<br>160.04 m/z ≥ 80% intensity<br>311.00 m/z ≥ 5% intensity<br>470.03 m/z ≥ 5% intensity | 0 | 0 | 0 | 0 | 654 | 587 |
| Number of MS/MS spectra containing flucloxacillin reporter ions<br>160.04 m/z ≥ 80% intensity<br>311.00 m/z ≥ 5% intensity<br>470.03 m/z ≥ 5% intensity | 0 | 0 | 798 | 758 | 0 | 0 |

In Gold Standard Method 1, the 1,241 MS/MS spectra from the dicloxacillin treated HSA samples that were found to contain dicloxacillin specific reporter ions at the intensity threshold of 80% 160.04 m/z, 5% 311.00 m/z and 5% 470.03 m/z, were evaluated to confirm the presence of a 469 Da adduct modification in the spectrum. As there were too many spectra to manually solve each of the 1,241 spectra individually, the following method was used to create a list of scan numbers representing spectra resulting from a peptide with a single 469 Da mass modification:

- 1,241 MS/MS spectra were found to contain dicloxacillin specific reporter ions with a minimum of the following intensities: 80% 160.04 m/z, 5% 311.00 m/z and 5% 470.03 m/z
- The targeted m/z of each MS/MS scan's precursor ion was noted and rounded down to the nearest integer yielding 145 distinct targeted m/z's.
- MS/MS spectra were extracted from the original list of 1,241 only for targeted m/z's that occurred at least 10 times resulting in 926 scans with 33 distinct m/z's. This constituted 75% of the original spectra (946/1,241).
- For each of the 33 distinct sets of targeted precursor ions one representative spectrum was manually evaluated to confirm the peptide contained a single 469 Da mass modification. If this could be confirmed, the entire set of scans at that targeted m/z was added to the gold standard list, if this could not be confirmed the entire set was excluded from the list.

**Using this completely algorithm free method, Gold Standard Method 1 resulted in 763 MS/MS spectra of dicloxacillin treated HSA that were confirmed to contain a single 469 Da dicloxacillin adduct modification.** These scans represented 22 distinct m/z species (Supplementary Figure 2a).

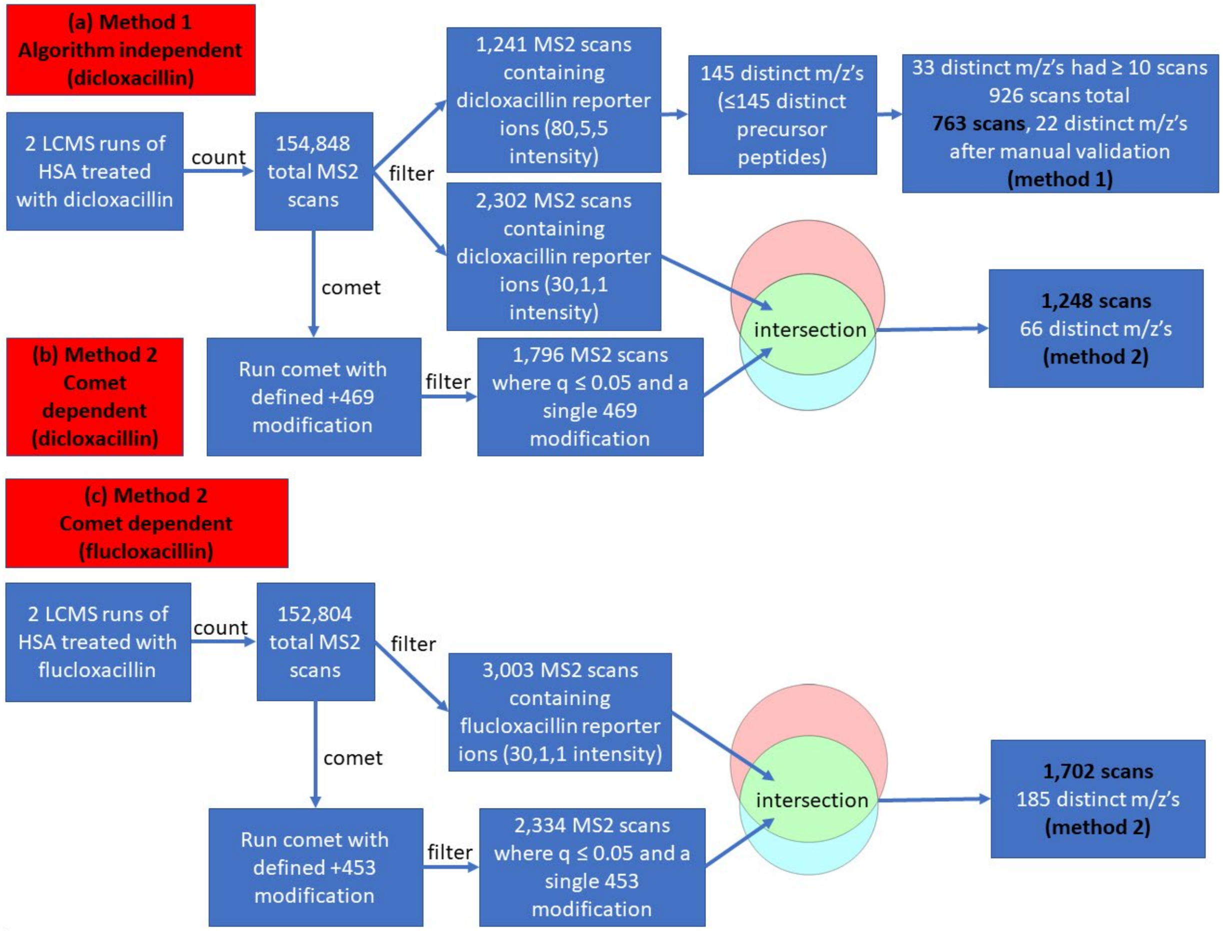

*Supplementary Figure 2: Workflow for the creation of 3 gold standard datasets. (a) Method 1 was fully manual, relied on no algorithms and resulted in 763 MS/MS scans with a known 469 Da dicloxacillin modification. (b,c) Method 2 relied on the presence of known β-lactam antibiotic reporter ions plus confident adduct modification mass identifications using a comet closed search. This method resulted in 1,248 and 1,702 MS/MS scans with known dicloxacillin or flucloxacillin adduct modifications, respectively.*

In Gold Standard Method 2, each of the 6 raw files described in Supplementary Table 2 were searched using ScanFinder for dicloxacillin and flucloxacillin reporter ions similarly as for method 1, but using the lower intensities stated in Supplementary Table 4.

*Supplementary Table 4: Number of MS/MS spectra that contain β-lactam antibiotic reporter ions at the defined m/z and lower intensities than searched for in Supplementary Table 3.*

| Sample Treatment | Untreated (rep 1) | Untreated (rep 2) | Flucloxacillin (rep 1) | Flucloxacillin (rep 2) | Dicloxacillin (rep 1) | Dicloxacillin (rep 2) |
|---|---|---|---|---|---|---|
| Dicloxacillin reporter ions<br>160.04 m/z ≥ 30% intensity<br>311.00 m/z ≥ 1% intensity<br>470.03 m/z ≥ 1% intensity | 0 | 0 | 0 | 0 | 1217 | 1085 |
| Flucloxacillin reporter ions<br>160.04 m/z ≥ 30% intensity<br>311.00 m/z ≥ 1% intensity<br>470.03 m/z ≥ 1% intensity | 0 | 0 | 1526 | 1477 | 13 | 5 |

The ScanFinder results in Supplementary Table 4 show that zero spectra from untreated HSA samples contained the combination of reporter ions searched for at the required lower intensities. Likewise, zero spectra in flucloxacillin treated samples contained dicloxacillin reporter ions at the specified lower intensities. 18 out of 154,848 spectra from dicloxacillin treated samples contained flucloxacillin specific reporter ions at the lower intensities used for this second set of ScanFinder searches. This is likely due to carryover from the flucloxacillin samples, which were run before the dicloxacillin samples on the same column. These data thus show that spectra picked out in the treated samples are specific to their respective treatments even at these lower intensity thresholds.

In addition to searching for MS/MS scans containing dicloxacillin and flucloxacillin reporter ions, we ran a closed comet search on each dataset, allowing for a defined variable modification of 469.0266 (the known dicloxacillin adduct mass) or 453.0561 (the known flucloxacillin adduct mass) on lysines. Scan numbers resulting in a confident (Percolator assigned $q \leq 0.05$) PSM containing either a single dicloxacillin or flucloxacillin adduct were noted (Supplementary Table 5).

*Supplementary Table 5: Number of MS/MS spectra that contain a single defined 469.0266 (dicloxacillin) or 453.0561 (flucloxacillin) adduct at a Percolator $q \leq 0.05$ based on a comet closed search.*

| Sample Treatment | Untreated (rep 1) | Untreated (rep 2) | Flucloxacillin (rep 1) | Flucloxacillin (rep 2) | Dicloxacillin (rep 1) | Dicloxacillin (rep 2) |
|---|---|---|---|---|---|---|
| Spectra containing a single 469.0266 modification (comet) | 140 | 119 | 124 | 184 | 916 | 880 |
| Spectra containing a single 453.0561 modification (comet) | 84 | 91 | 1149 | 1185 | 188 | 224 |

To create the Method 2 Gold Standard datasets we listed the scan numbers for the lower intensity ScanFinder search plus the scan numbers for the comet search specific to each antibiotic and extracted the intersection of those scan numbers. In other words, only scan numbers that showed the required reporter ions AND resulted in a confident comet PSM yielding a single antibiotic adduct of the specified mass were added to the gold standard list (Supplementary Figure 2b and c). **Gold Standard Method 2 resulted in 1,248 MS/MS spectra (66 distinct m/z's) of dicloxacillin treated HSA confirmed to contain a single 469 Da dicloxacillin adduct modification. The same method yielded 1,702 MS/MS spectra (185 distinct m/z's) of flucloxacillin treated HSA confirmed to contain a single 453 Da flucloxacillin adduct modification.**

The procedure above resulted in 3 sets of gold standard scan numbers: (1) dicloxacillin gold standard method 1 (763 spectra with known +469 Da single modifications); (2) dicloxacillin gold standard method 2 (1,248 spectra with known +469 Da single modifications); and (3) flucloxacillin gold standard method 2 (1,702 spectra with known +453 Da single modifications). These 3 sets of scan numbers constituted a combined total of 2,979 unique MS/MS spectra all of which were modified by a single xenobiotic-protein adduct of known mass. Various searches were then run on the complete datasets and only the open modification masses for these specific spectra were extracted for the gold standard comparisons. Precision and recall could then be calculated for each method for any search performed, based on the fraction of correct answers and the total number of correct answers.

The data presented in the main manuscript combines results from all three methods for brevity (Main manuscript Figure 2), however results from the individual methods are presented below (Supplementary Figures 3, 4, 5 and 6), illustrating that PSMs derived from each individual method produced similar results.

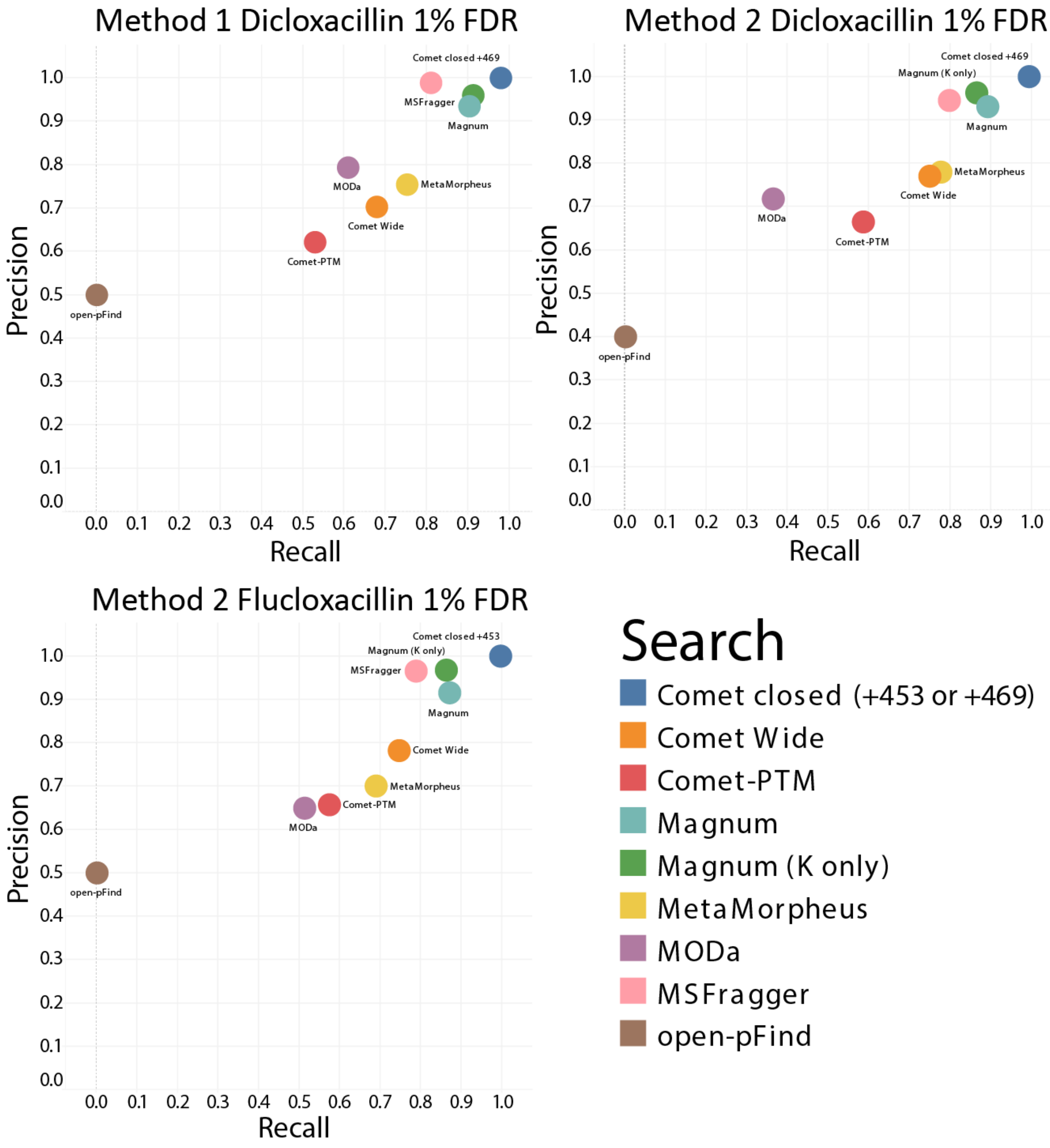

*Supplementary Figure 3: Precision recall plot of adduct masses reported at 1% FDR by 7 open search algorithms for MS/MS spectra definitively determined to result from a peptide containing a single +469 Da (dicloxacillin) or +453 Da (flucloxacillin) modification. Results are shown separately for each of the gold standards datasets described above. Results from closed comet searches using defined modifications of 469 or 453 were included as a positive control. An open modification mass returned by an algorithm is defined as correct if it is within ±3 Da of the known correct modification mass (469 or 453). Magnum was run allowing for open masses on any amino acid (Magnum) or restricted to lysines only (Magnum K), the previously published residue modified by dicloxacillin and flucloxacillin adducts.*

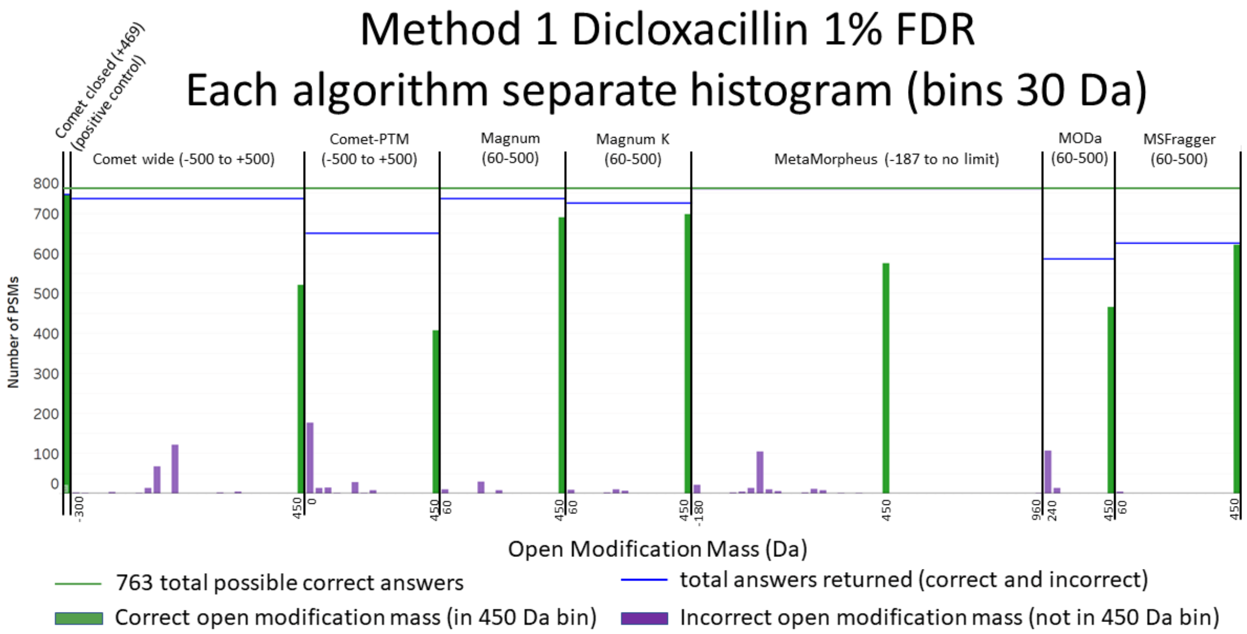

*Supplementary Figure 4: Histograms showing the distribution of open modification masses reported by each algorithm for the 763 gold standard spectra generated from dicloxacillin treated samples using method 1, described above. Results are shown at 1% FDR and are distributed between incorrect masses (purple bars, >3 Da from known correct modification mass) and correct masses (green bars, within ±3 Da of the known correct modification mass). The first and last mass bin are labeled on the x axis. The open mass range searched by each algorithm is shown in parenthesis above each plot. Bins are 30 Da wide and all correct answers fall within the 450 Da bin. Incorrect masses within the 450 Da bin are shaded purple.*

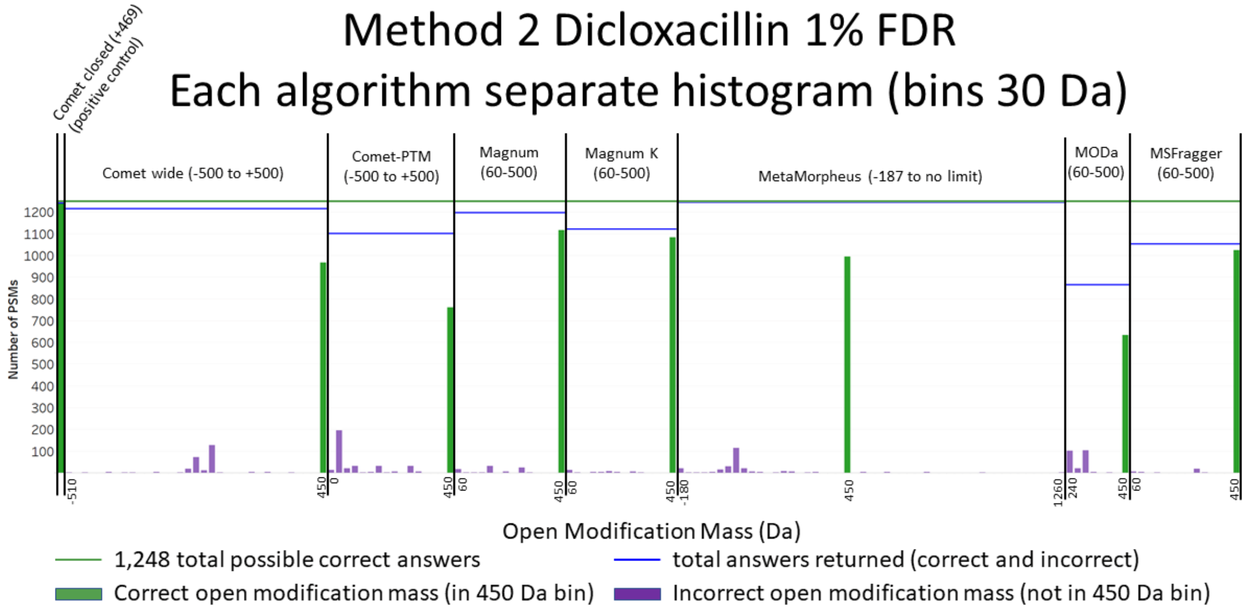

*Supplementary Figure 5: Histograms showing open modification masses reported by each algorithm for the 1,248 gold standard spectra generated from dicloxacillin treated samples by method 2, described above. Results at 1% FDR are distributed between incorrect masses (purple bars, >3 Da from known correct mass) and correct masses (green bars, within ±3 Da of the known correct modification mass). The first and last mass bin are labeled on the x axis. The open mass range searched by each algorithm is shown in parenthesis above each plot. Bins are 30 Da wide and all correct answers fall within the 450 Da bin. Incorrect masses within the 450 Da bin are shaded purple.*

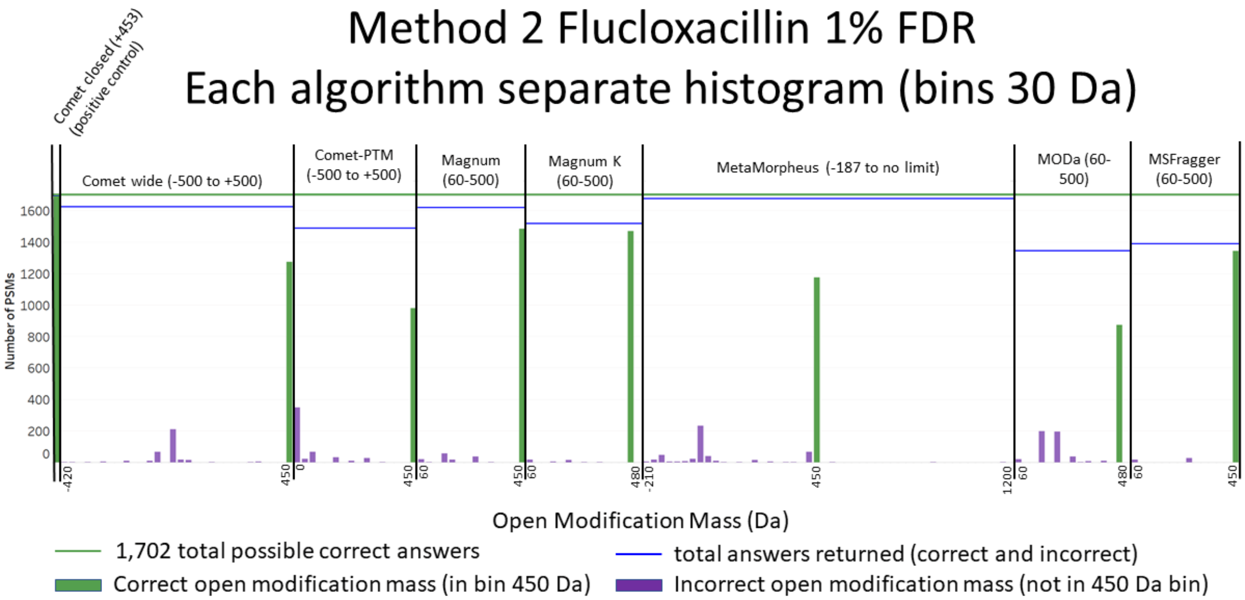

*Supplementary Figure 6: Histograms showing the distribution of open modification masses reported by each algorithm for the 1,702 gold standard spectra generated from flucloxacillin treated samples using method 2, described above. Results are shown at 1% FDR and are distributed between incorrect masses (purple bars, >3 Da from known correct modification mass) and correct masses (green bars, within ±3 Da of the known correct modification mass). The first and last mass bin are labeled on the x axis. The open mass range searched by each algorithm is shown in parenthesis above each plot. Bins are 30 Da wide and all correct answers fall within the 450 Da bin. Incorrect masses within the 450 Da bin are shaded purple.*

*Supplementary Table 6: Gold standard results returned by open-pFind at 1% FDR. Of the 9 results returned, 4 had masses within the tolerance deemed correct (flucloxacillin 453 ± 3 Da; dicloxacillin 469 ± 3Da ) for the purposes of our gold standard analysis. The masses matched by open-pFind correspond to the following three Unimod[10] entries: (1) Accession #: 409; Interim Name: FMN; Monoisotopic Mass: 454.088965. (2) Accession #: 1431; Interim Name: Hex(1)NeuAc(1); Monoisotopic Mass: 453.148240. (3) Accession #: 1375; Interim Name: dHex(1)Hex(2); Monoisotopic Mass: 470.163556. As open-pFind is unable to assign the real masses of flucloxacillin (453.0561) and dicloxacillin (469.0266) adducts, most spectra do not result in correct or confident assignments. This issue will exist for any modifications not in the Unimod database prior to searching.*

| URL to Annotated Spectrum on Limelight | Open Mod Mass | Reported Peptide | q-value |
|---|---|---|---|
| **HSA + flucloxacillin - replicate 1** | | | |
| https://limelight.yeastrc.org/limelight/d/pg/spectrum-viewer/ps/1893/psm/78163210 | 454.08897 | YKAAFTEC[454.09]CQAADK | 0.0031088 |
| https://limelight.yeastrc.org/limelight/d/pg/spectrum-viewer/ps/1893/psm/78172798 | 38.930957 | MPC[57.02]AED[37.95]YLSVVLN[0.98]QLCVLHEK | 0.0073529 |
| **HSA + flucloxacillin - replicate 2** | | | |
| https://limelight.yeastrc.org/limelight/d/pg/spectrum-viewer/ps/1894/psm/78198359 | 453.14824 | RYKAAFT[453.15]ECC[57.02]QAADK | 0.0010267 |
| https://limelight.yeastrc.org/limelight/d/pg/spectrum-viewer/ps/1894/psm/78210881 | 324.03587 | n[324.04]KASSAKQR | 0.0036443 |
| **HSA + dicloxacillin - replicate 1** | | | |
| https://limelight.yeastrc.org/limelight/d/pg/spectrum-viewer/ps/1895/psm/67218827 | 38.01565 | DEGK[38.02]ASSAKQR | 0.0011351 |
| https://limelight.yeastrc.org/limelight/d/pg/spectrum-viewer/ps/1895/psm/67219788 | 0 | AVM[15.99]DDFAAFVEK | 0.012616 |
| https://limelight.yeastrc.org/limelight/d/pg/spectrum-viewer/ps/1895/psm/67222933 | 470.16356 | AT[470.16]EEQLK | 0.0011351 |
| **HSA + dicloxacillin - replicate 2** | | | |
| https://limelight.yeastrc.org/limelight/d/pg/spectrum-viewer/ps/1896/psm/71787832 | 37.946941 | M[15.99]AAQGE[37.95]PGYLAAQSDPGSNSER | 0.014199 |
| https://limelight.yeastrc.org/limelight/d/pg/spectrum-viewer/ps/1896/psm/71789821 | 470.16356 | RPC[57.02]FSALEVDET[470.16]YVPK | 0.0072029 |

### Supplementary Note 3: Limelight

#### Description

Limelight is a web application built to analyze, visualize, and share bottom-up MS proteomics data. It is open source and freely available at http://limelight-ms.org.

Central to its design is the separation of data analysis and visualization from the software pipeline that generated the data. Limelight's design makes as few assumptions about the data as possible, providing a generalized platform that fully and equally supports data generated by any MS database search pipeline while providing access to the full stack of proteomics data.

All the native results of each pipeline (e.g. p-values, q-values, Xcorrs, etc.) are available within Limelight, and may be used as filtering and analysis criteria for viewing single searches or combining multiple searches even if those searches used different pipelines. Limelight achieves pipeline-independence via an XML schema, called Limelight XML, that encodes all the results and scores in a generalized way. Limelight XML encodes which programs and versions were run, which scores are present from each program and, importantly, how those scores are to be treated (i.e. are larger or smaller numbers more significant). Not only can the output of any pipeline be represented, but this provides a level of provenance for the data that enhances reproducibility. Once data are represented as Limelight XML, they may be imported into Limelight via the web interface or via an upload API.

The authors of Limelight have written Limelight XML converters for many popular software pipelines including Comet[4], Percolator[11], the Trans-Proteomic Pipeline (TPP)[12], Crux[13], MSFragger[14], open-pFind[15], Comet-PTM[16], MetaMorpheus[17], MODa[18], TagGraph[19] and Magnum (this paper).

Limelight is written in Java and other standard web technologies, including Javascript, HTML and CSS. Our GitHub repository resides at: https://github.com/yeastrc/limelight-core and a current list of Limelight XML importers may be seen by visiting https://limelight-ms.readthedocs.io/. The Limelight XML schema may be seen at https://github.com/yeastrc/limelight-import-api/tree/master/xsd.

#### Experiment Builder

Limelight includes a novel interface to defining the experimental conditions for a set of separate mass spectrometry searches. This is referred to in Limelight as the "experiment builder". Using the experiment builder, users may iteratively build up an experimental design by adding condition groups and then conditions within those condition groups. For example, a user may designate that they have two technical replicates in their experimental design. Then a user may designate they have 6 timepoints in their experimental design. Then a user may add a condition group for "tissue type" and add 3 conditions in this condition group: "heart", "lung", and "liver." (See Supplementary Figure 7, below). Once an experiment has been structured and searches have been assigned to each cell in the resulting grid, users may view the experiment to automatically compare searches and groups of searches according to their experimental design.

(A)

**Experiment Layout**

| Tech Rep 1 | Tech Rep 2 |
|---|---|
| Empty | Empty |

(B)

**Experiment Layout**

| | T1 | T2 | T3 | T4 | T5 | T6 |
|---|---|---|---|---|---|---|
| Tech Rep 1 | Empty | Empty | Empty | Empty | Empty | Empty |
| Tech Rep 2 | Empty | Empty | Empty | Empty | Empty | Empty |

(C)

**Experiment Layout**

| | | T1 | T2 | T3 | T4 | T5 | T6 |
|---|---|---|---|---|---|---|---|
| Heart | Tech Rep 1 | Empty | Empty | Empty | Empty | Empty | Empty |
| | Tech Rep 2 | Empty | Empty | Empty | Empty | Empty | Empty |
| Lung | Tech Rep 1 | Empty | Empty | Empty | Empty | Empty | Empty |
| | Tech Rep 2 | Empty | Empty | Empty | Empty | Empty | Empty |
| Liver | Tech Rep 1 | Empty | Empty | Empty | Empty | Empty | Empty |
| | Tech Rep 2 | Empty | Empty | Empty | Empty | Empty | Empty |

*Supplementary Figure 7: Iteratively building an experiment using the experiment builder. (A) The user has added two technical replicates. (B) The user has added six time points. (C) The user has added three tissue types. The user may continue to iteratively add condition groups and conditions to a level of infinite complexity to fully represent their experimental design. Once done, the user may click in the empty cells to add searches from the experiment to each cell to organize the data.*

#### Score filters and cutoffs

On all pages (peptide, protein, and modification views), users may filter the data present on the page according to any score present in the respective software pipeline that was used to search the data. For example, E-value, Xcorr, or any other score native to Comet may be used for filtering; and q-value, posterior error probability, or any other score native to Percolator maybe used for filtering. If the pipeline was a multistep pipeline, any score from any step may be used.

The currently-used filters are shown at the top of the page, and may be easily changed by clicking on any of the displayed filters to bring up an interface for changing the cutoffs. This interface includes a text box for every type of score present in the native pipeline and entering new values and clicking “Save” will result in those filters being applied.

#### Single protein view

Wherever protein names are displayed in Limelight, they be clicked to view the single protein view. This view provides data visualizations for a single protein identified in the experiment. This includes the name and description of the protein, its sequence coverage, and a list of all peptides localized to this protein. This list of peptides may be expanded to view all PSMs for each peptide, and each PSM will include native scores and links to view spectra.

A critical aspect of the single protein view is the ability to apply advanced filtering to the peptide list to focus on peptides relevant to a specific question. This peptide list may be filtered by which modifications were identified, whether it was uniquely identified in this protein, peptide sequence, and whether it overlapped specific positions in the protein sequence. The sequence coverage map is interactive, and by clicking positions users may filter for only peptides covering that position (control-click to select multiple positions).

#### Peptide view

The peptide view provides a peptide-focused view of the experimental results. This view shows all the peptides identified in the experiment, given the current cutoffs (see above). Each listed peptide includes that peptide's sequence, whether it uniquely matched a single protein, in which proteins it was localized, and the spectral count. Each peptide may be clicked on and expanded to reveal all PSMs associated with that peptide, including links to view the annotated spectrum and native scores and annotations from the respective software pipeline.

The peptide list may be filtered according to which modification masses were observed, whether the peptide is unique to a single protein, peptide sequence, or a set of specific proteins (and positions within those proteins). This enables users to perform filtering such as listing only the peptides that contain a phosphorylation and localize to the C-terminal region of all the variants of a given protein identified in the experiment.

#### Protein view

The protein view provides a protein-focused view of the experimental results. It lists all the proteins identified in the experiment, given the current cutoffs (see above). Each row lists the protein's name, description, sequence coverage, number of peptides, number of unique peptides, and number of PSMs (that meet the current cutoffs). Each row may be clicked to expand to view peptides, and each peptide may be clicked to expand to view PSMs.

#### Modification view

The modification list view provides a modification-focused view of the experimental results. All modifications identified in the search are displayed in two ways. First the modifications are displayed as a heatmap, where the modification masses are displayed on the x-axis, the currently-shown searches are displayed on the y-axis, and the matrix is shaded according to statistics associated with a given modification mass in a given search. These statistics may be PSM count, scan count, ratio of all PSMs or scans that have that modification mass, or a statistical transformation (described below). The heatmap is interactive, and users may click (and optionally drag) within the visualization to filter which modifications are displayed below in the modification table. The heatmap (and table below) may be further filtered and customized by choosing how to scale the colors, the minimum and maximum modification mass to display, statistical transformations, and filtering based on specific proteins (and positions in those protein to which the modifications must localize).

The modification table below lists each modification on separate rows and may be filtered by interacting with the visualization above. Each row includes the modification mass, links to external modification annotation resources, and the value of the current statistic being displayed in the heatmap for that modification in each of the currently shown searches. Each row may be clicked and expanded to view all proteins, positions in those proteins this given mod localizes, which residues in that protein are modified, and the PSM count for the given modification mass in the respective protein. Each protein may be expanded to view the list of peptides that contain that modification mass for this protein. Associated with each listed peptide are the N- and C-terminal residues in the protein that flank the peptide, the number of PSMs for that peptide, the positions in the respective protein covered by this modification mass in this peptide, and a list of residues (amino acid codes) modified by this modification mass in this peptide. Each peptide may be expanded to view all PSMs (and associated scores) associated with this peptide, including links to view underlying spectra.

#### Visualizations and Transformations

The following statistical transformations are available in the modification view data visualization:

- Scaled mean difference: For each mod mass and search display: $(x - \mu) / \mu$, where x is the count or ratio for a mod mass in a search and μ is the mean for that mod mass across all searches.
- Per-mod Z-score: For each mod mass and search display: $(x - \mu) / s$, where x is the count or ratio for a mod mass in a search, μ is the mean for a mod mass across all searches, and s is the standard deviation for this mod mass across all searches.
- Global Z-score: For each mod mass and search display: $(x - \mu) / s$, where x is the count or ratio for a mod mass in a search, μ is the mean for all mod masses across all searches, and s is the standard deviation across all mod masses in all searches.
- Global P-value: For each mod mass and search display: p, where p is the Bonferroni-corrected p-value associated with the global Z-score (the probability of observing a z-score of that magnitude or greater by chance given a normal distribution with the observed mean and standard deviation).
- Global Q-value: For each mod mass and search display: q, where q is the Benjamini-Hochberg q-value associated with the global Z-score (the probability of observing a z-score of that magnitude or greater by chance given a normal distribution with the observed mean and standard deviation).

#### Two-tailed test of proportions

Users of Limelight may download a report that attempts to identify the most statistically significant modifications in one set of searches versus another. This is done by calculating the ratio of PSMs (or scans) that have the given modification mass and dividing by the total number of PSMs. This ratio is calculated for a given set of searches (e.g., the two biological replicates for a control) and compared against another set of searches (e.g., the two biological replicates for a treatment) and a Z-score is calculated using the canonical test of proportions (see Supplementary Equation 3). In this equation, x1 is the number of PSMs or scans with the modification mass in the first set of searches, x2 is the number of PSMs or scans with the modification mass in the second set of searches, n1 is the total PSM count for the first set of searches, and n2 is the total PSM count for the second set of searches. This calculation produces a Z-statistic which may be used to compare modification masses for significance and may be converted to a p-value using a lookup table. The p-values in the report are then Bonferroni-corrected to account for multiple hypothesis tests (i.e., the number of modification masses tested).

$$Z = \frac{\frac{x_1}{n_1} - \frac{x_2}{n_2}}{\sqrt{p(1-p)\left(\frac{1}{n_1} + \frac{1}{n_2}\right)}} \; and \;\; p = \frac{x_1 + x_2}{n_1 + n_2}$$

*Supplementary Equation 3: Canonical test of proportions*

### Supplementary Note 4: Development and validation of adduct discovery pipeline using dicloxacillin, flucloxacillin and HSA

#### Open modifications unrelated to exposure

Estimates suggest over 50% of spectra remain invisible to traditional "closed" search methods[20]. The prevailing hypothesis is that unidentified spectra constitute peptides not represented in a typical search space: they have undefined post-translational modifications (PTMs), chemical modifications, variant protein sequences or unpredictable cleavage aberrations. Several "open" search strategies have been developed to shed light on these "dark" spectra. These strategies enable mass tolerant database searching and have allowed peptide spectrum matches (PSMs) to be made from a large proportion of previously unassigned spectra in shotgun proteomics data[14–22]. Past open search publications have focused largely on global open modification analyses of large and complex proteomics datasets and show increased numbers of PSMs as well as complex modification landscapes not previously accessible via traditional closed searching.

Limelight is designed to support all bottom-up MS proteomics pipelines and is fully capable of analyzing and visualizing open modification data from unexposed samples. We performed open searching, using 7 open search algorithms, on MS data we acquired from unexposed human serum albumin (HSA) samples (Supplementary Figure 8).

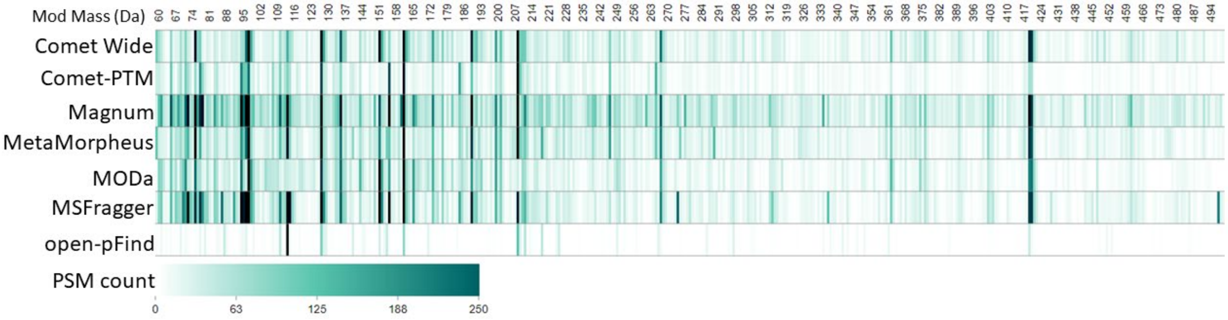

*Supplementary Figure 8: Open modification masses in untreated, purified human serum albumin as identified by 7 different open search algorithms and visualized by Limelight. Open modifications identified in the range of 60 to 500 Da are shown at 1% FDR. A live view of these data is available here: https://limelight.yeastrc.org/limelight/go/Vc6Z8ppwpl*

These data show that even in a purified protein sample, open searching results in PSMs containing open modification masses across the entire range of masses available to the search algorithms based on the search parameters they were given.

We then performed both open and closed searches on LC-MS/MS data from 6 untreated and flucloxacillin or dicloxacillin exposed HSA samples. In agreement with previous open search publications, open search analyses resulted in more than twice the number of PSMs as closed searching performed on the same data (Supplementary Figure 9).

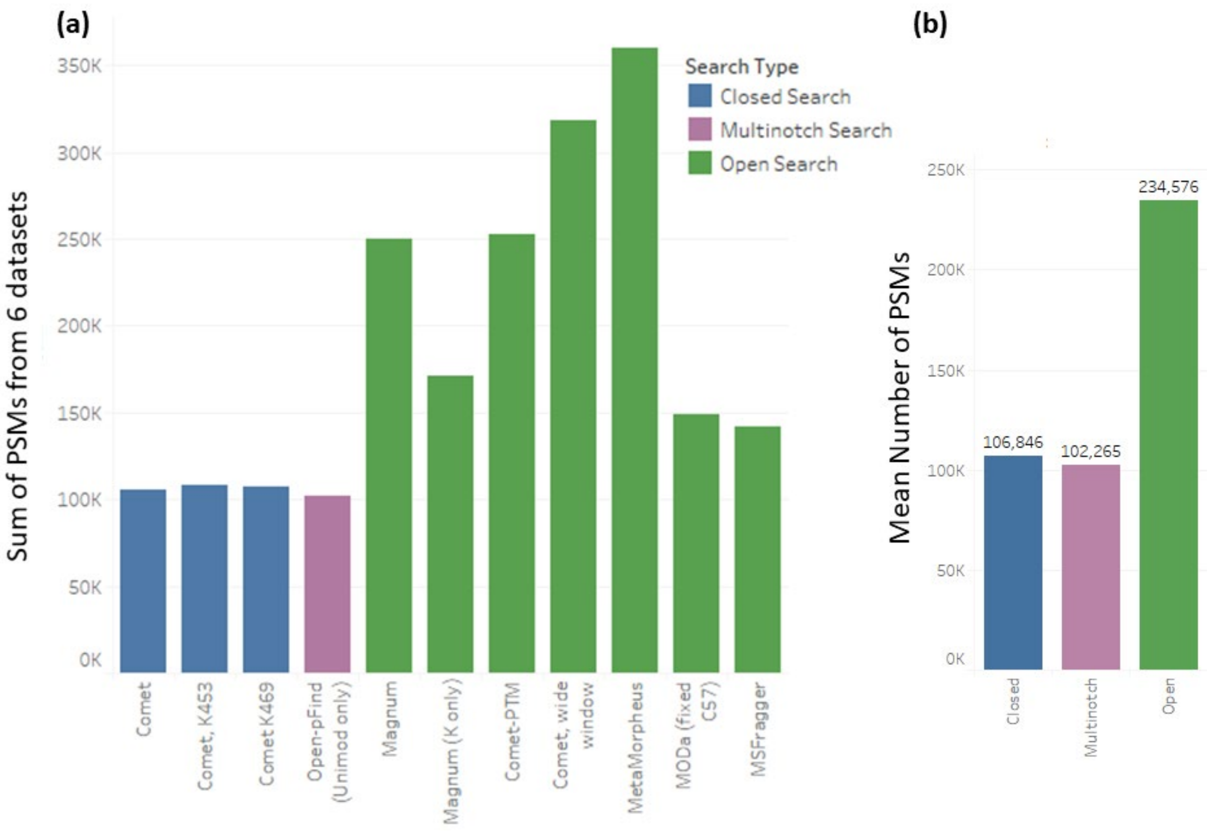

*Supplementary Figure 9: The number of PSMs at 1% FDR resulting from searching 6 untreated and dicloxacillin and flucloxacillin exposed HSA datasets with open and closed search algorithms. (a) The number of PSMs shown is the sum of all 6 datasets for each algorithm. (b) The mean number of PSMs for each algorithm type was calculated from the data in (a). All search engines were configured with 57.02146 on C and 15.99490 on M as variable modifications except MODa which does not allow variable modifications to be defined. Magnum was run allowing open modifications on all residues (Magnum) or restricted to lysines only (Magnum K only). Closed comet searches were performed with the variable modifications previously stated (comet), plus defined variable modifications of the previously published adduct masses of dicloxacillin or flucloxacillin allowed on lysines (comet, K469 and K453).*

#### Discovery of dicloxacillin/flucloxacillin adducts in HSA

Previously published studies[8,9] used multiple methods to determine that dicloxacillin and flucloxacillin produce 469 Da and 453 Da adduct modifications, respectively, on HSA lysine residues (Supplementary Figure 1).

We acquired untargeted MS data of unexposed, dicloxacillin, and flucloxacillin exposed human serum albumin (HSA) and searched the resulting data using 7 different algorithms. Initially, we compared the open modification masses identified by Magnum in 2 untreated, 2 dicloxacillin treated, and 2 flucloxacillin treated samples and found that the same set of 441 open modification masses (rounding to the nearest integer) were identified in all 6 samples. These masses constituted all the open modification masses available to Magnum based on the search parameters it was given. To reduce this apparent noise, we compared open modification masses identified by ≥10 PSMs in untreated versus treated samples, however none of the adduct masses known to result from dicloxacillin or flucloxacillin exposure were found exclusively in their respective treatment groups. An analysis of the same spectra with an alternative open search tool, MSFragger[14], likewise found at least one PSM for each of the 441 open modification masses allowed and did not find known exposure related adduct masses uniquely in treated samples using either a 1 or 10 PSM cutoff (Supplementary Figure 10). These observations are therefore likely inherent to open searching in general and highlight the difficulty in identifying exposure-specific adducts against the background produced by open modification searching.

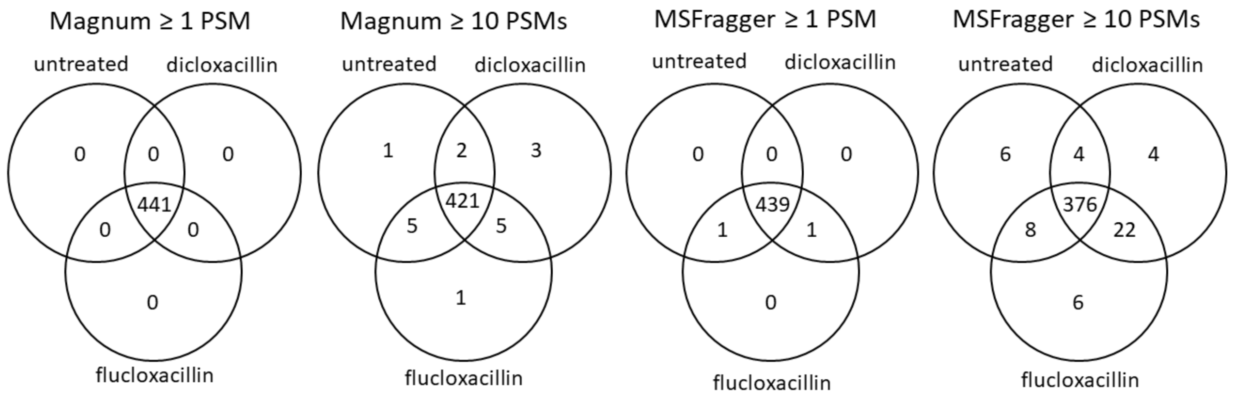

| Search Engine/PSM cutoff | Magnum/ none | Magnum/10 PSMs | MSFragger/ none | MSFragger/ 10PSMs |
|---|---|---|---|---|
| Num. mod masses common to all treatments | 441 | 421 | 439 | 376 |
| Num. mod masses unique to untreated samples | 0 | 1 | 0 | 6 |
| Num. mod masses unique to dicloxacillin treated samples | 0 | 3 | 0 | 4 |
| Num. mod masses unique to flucloxacillin treated samples | 0 | 1 | 0 | 6 |
| Treatment specific mod masses unique to treated samples? | no | no | no | no |

*Supplementary Figure 10: Open modification masses identified by Magnum or MSFragger in untreated, dicloxacillin treated or flucloxacillin treated human serum albumin (HSA). The number of open modification masses common or unique to the different treatments is shown. Modification masses known to result from dicloxacillin and flucloxacillin treatment were observed but were not unique to treated samples. Data are filtered at a 1% FDR. The full data presented here can be found at: https://limelight.yeastrc.org/limelight/go/45kI4Zi9cE (Magnum) and https://limelight.yeastrc.org/limelight/go/tTBnwmSdzk (MSFragger).*

We overcame these difficulties using the two-tailed test of proportions described in the main manuscript and Supplementary Note 3. We built this method into Limelight and were able to use it to highlight exposure related modifications rapidly and sensitively by comparing untreated with treated samples (or sample groups). These data are presented in the main manuscript for PSMs generated by Magnum. We also searched the same MS data using 6 other open search algorithms for comparison with Magnum. PSMs were imported into Limelight for downstream analysis and a two-tailed test of proportions comparing untreated with dicloxacillin or flucloxacillin treated HSA was performed with PSMs generated by each algorithm.

These analyses resulted in 469 Da (the correct mass) or 471 Da (a monoisotopic mass misassignment of 2 Da) being the most significantly enriched mass for all algorithms tested (Supplementary Figure 11) except open-pFind which could not identify dicloxacillin or flucloxacillin adducts for the reasons outlined in the main manuscript. Equivalent comparisons with flucloxacillin treated samples resulted in 453 Da being the most significantly enriched mass for all algorithms except open-pFind. Complete data is shown in **Supplementary File penicillin_ twotail.xlsx** along with links to all data in Limelight.

##### HSA ± Dicloxacillin (replicate 1): 469 Da

| Search Algorithm | Mod Mass (Da) | Untreated PSM count | Treated PSM count | P value | Z Score |
|---|---|---|---|---|---|
| Comet Wide | 471 | 5 | 342 | 0 | -18.421 |
| Comet-PTM | 469 | 1 | 120 | 0 | -11.166 |
| Magnum | 469 | 11 | 387 | 0 | -19.440 |
| Magnum K | 469 | 7 | 375 | 0 | -19.638 |
| MetaMorpheus | 469 | 3 | 353 | 0 | -18.500 |
| MODa | 471 | 0 | 181 | 0 | -14.816 |
| MSFragger | 471 | 7 | 309 | 0 | -17.184 |
| open-pFind | 156 | 4 | 88 | 0 | -9.702 |

##### HSA ± Dicloxacillin (replicate 2): 469 Da

| Search Algorithm | Mod Mass (Da) | Untreated PSM count | Treated PSM count | P value | Z Score |
|---|---|---|---|---|---|
| Comet Wide | 471 | 4 | 306 | 0 | -16.79 |
| Comet-PTM | 469 | 0 | 121 | 0 | -10.93 |
| Magnum | 469 | 16 | 364 | 0 | -17.83 |
| Magnum K | 469 | 9 | 344 | 0 | -17.68 |
| MetaMorpheus | 469 | 4 | 345 | 0 | -18.06 |
| MODa | 471 | 0 | 162 | 0 | -13.14 |
| MSFragger | 471 | 3 | 281 | 0 | -16.29 |
| open-pFind | 209 | 115 | 49 | 5.11154e-05 | 5.17 |

##### HSA ± Flucloxacillin (replicate 1): 453 Da

| Search Algorithm | Mod Mass (Da) | Untreated PSM count | Treated PSM count | P value | Z Score |
|---|---|---|---|---|---|
| Comet Wide | 453 | 25 | 561 | 0 | -22.78 |
| Comet-PTM | 453 | 10 | 304 | 0 | -17.28 |
| Magnum | 453 | 18 | 726 | 0 | -26.82 |
| Magnum K | 453 | 7 | 713 | 0 | -27.93 |
| MetaMorpheus | 453 | 15 | 686 | 0 | -25.70 |
| MODa | 453 | 6 | 378 | 0 | -20.72 |
| MSFragger | 453 | 15 | 583 | 0 | -23.60 |
| open-pFind | 128 | 98 | 37 | 0.000973345 | 4.59 |

##### HSA ± Flucloxacillin (replicate 2): 453 Da

| Search Algorithm | Mod Mass (Da) | Untreated PSM count | Treated PSM count | P value | Z Score |
|---|---|---|---|---|---|
| Comet Wide | 453 | 21 | 572 | 0 | -21.87 |
| Comet-PTM | 453 | 6 | 289 | 0 | -16.12 |
| Magnum | 453 | 15 | 726 | 0 | -25.71 |
| Magnum K | 453 | 7 | 710 | 0 | -25.67 |
| MetaMorpheus | 453 | 24 | 658 | 0 | -23.97 |
| MODa | 453 | 4 | 357 | 0 | -18.89 |
| MSFragger | 453 | 14 | 597 | 0 | -23.24 |
| open-pFind | 156 | 0 | 80 | 0 | -9.03 |

*Supplementary Figure 11: A two-tailed test of proportions performed within Limelight identifies treatment specific adducts in HSA. Tests were done on PSMs identified by 7 different open search algorithms. Results were sorted on the absolute value of the Z score (large to small) followed by the magnitude of the P value (small to large). The top result is shown for each algorithm, representing the most significantly enriched mass found by each algorithm. Magnum was run allowing open modifications on all residues (Magnum) or restricted to lysines only (Magnum K). Data are shown at 1% FDR. Full data is available in Supplementary File penicillin_twotail.xlsx.*

In all cases Magnum identified the most treatment related PSMs as well as resulting in the largest Z Score for the correct mass compared to other algorithms. These data show a two-tailed test of proportions comparing treated versus untreated samples, is effective in highlighting exposure specific adducts from PSMs generated by several open search algorithms and that Magnum is the most sensitive algorithm.

The complete list of +469 Da and +453 Da modified HSA peptides identified by Magnum at 1% FDR is shown in Supplementary Table 7. All spectra can be manually inspected via Limelight's built in spectrum viewer, Lorikeet[6], using the links in the table caption. A representative, manually annotated spectrum of a dicloxacillin adducted peptide is depicted in the main manuscript Figure 5 and a similar annotated spectrum of a flucloxacillin adducted peptide is depicted in the Supplementary Figure 12.

*Supplementary Table 7: Dicloxacillin and flucloxacillin adducted peptides identified by Magnum. Peptides with at least one dicloxacillin (160.04, 311.00 or 470.03) or flucloxacillin (160.04 or 295.03 or 454.06) reporter ion are in black. Peptides with no identified reporter ions are in blue. The 46 dicloxacillin adducted peptides can be viewed on Limelight here: https://limelight.yeastrc.org/limelight/go/avyNUyhojE. The 55 flucloxacillin adducted peptides can be viewed here: https://limelight.yeastrc.org/limelight/go/XFLoAxI0Uw. Equivalent views additionally filtering for presence of reporter ions can be found here https://limelight.yeastrc.org/limelight/go/coQIdEjkRI (for dicloxacillin plus reporter ions) and here https://limelight.yeastrc.org/limelight/go/5aPYwCwMLc (for flucloxacillin plus reporter ions). Data are shown at a percolator q ≤ 0.01.*

| Dicloxacillin Adducted Peptide Sequence | PSM Count ( run 2195) | PSM Count (run 2197) |
|---|---|---|
| AFKAWAVAR-(469) | 11 | 12 |
| ASSAKQR-(469) | 78 | 81 |
| ATKEQLK-(469) | 97 | 94 |
| CCAAADPHEC[57]YAK-(469) | 0 | 2 |
| CCAAADPHECYAK-(469) | 0 | 1 |
| D[469]EGKASSAK | 1 | 0 |
| DEGKASSAK-(469) | 6 | 6 |
| EQLKAVMDDFAAFVEK-(469) | 1 | 1 |
| FGERAFK-(469) | 10 | 15 |
| FKDLGEENFK-(469) | 0 | 2 |
| KQTALVELVK-(469) | 2 | 7 |
| KYLYEIAR-(469) | 1 | 0 |
| L[469]VNEVTEFAKTCVADESAENCDK | 1 | 0 |
| LAKTYETTLEK-(469) | 30 | 25 |
| LC[57]TVATLR-(469) | 1 | 1 |
| LDELRDEGK[469]ASSAK | 0 | 1 |
| LDELRDEGKASSAK-(469) | 14 | 8 |
| LKC[57]ASLQK-(469) | 9 | 6 |
| LKCASLQK-(469) | 4 | 5 |
| MPC[57]AEDYLSVVLNQLC[57]VLHEK-(469) | 2 | 0 |
| NECFLQHKDDNPNLPR-(469) | 5 | 4 |

| | | |
|---|---|---|
| NLGKVGSK-(469) | 61 | 49 |
| RHPDYSVVLLLR-(469) | 0 | 2 |
| RPC[57]F[469]SALEVDETYVPK | 0 | 1 |
| RPC[57]FS[469]ALEVDETYVPK | 6 | 4 |
| RPC[57]FSAL[469]EVDETYVPK | 2 | 1 |
| RPC[57]FSALEVDETYVPK-(469) | 0 | 1 |
| RPCFSALEVDETYVPK-(469) | 0 | 1 |
| RYKAAFTEC[57]C[57]QAADK-(469) | 8 | 1 |
| RYKAAFTEC[57]CQAADK-(469) | 4 | 1 |
| RYKAAFTECCQAADK-(469) | 3 | 0 |
| SHC[57]IAEVE[469]NDEM[16]PADLPSLAADFVESK | 1 | 0 |
| SHC[57]IAEVENDEMPADLPSL[469]AADFVESK | 0 | 1 |
| SHC[57]IAEVENDEMPADLPSLAADFVESK-(469) | 0 | 2 |
| TC[57]VADESAENC[469]DK | 1 | 2 |
| VGSKC[57]C[57]K-(469) | 14 | 11 |
| VGSKCCK-(469) | 2 | 1 |
| YKAAFTEC[57]C[57]QAADK-(469) | 6 | 10 |
| YKAAFTEC[57]CQAADK-(469) | 1 | 0 |
| YKAAFTECC[57]QAADK-(469) | 1 | 3 |
| YKAAFTECCQAADK-(469) | 1 | 1 |
| **Flucloxacillin Adducted Peptide Sequence** | **PSM Count (run 2191)** | **PSM Count (run 2193)** |
| A[453]FKAWAVAR | 11 | 4 |
| AA[453]FTEC[57]C[57]QAADK | 1 | 0 |
| AAFTECC[57]QAADK-(453) | 1 | 4 |
| AAFTECCQAADK-(453) | 2 | 1 |
| AFK[453]AWAVAR | 0 | 1 |
| AFKAWAVAR-(453) | 64 | 67 |
| ASSAKQR-(453) | 154 | 154 |
| ATKEQLK-(453) | 107 | 99 |
| AVMD[453]DFAAFVEK | 2 | 2 |
| AVMDD[453]FAAFVEK | 0 | 1 |
| AVMDDFAAFV[453]EK | 0 | 1 |
| AVMDDFAAFVEK-(453) | 2 | 1 |
| AWAVARLSQR-(453) | 1 | 3 |
| CC[57]AAADPHEC[57]YAK-(453) | 0 | 1 |
| CCAAADPHEC[57]YAK-(453) | 2 | 0 |
| DEGKASSAK-(453) | 9 | 9 |
| EFNAETFTFHADICTLSEK-(453) | 0 | 2 |
| EQLKAVMDDFAAFVEK-(453) | 1 | 0 |
| FGERAFK-(453) | 9 | 19 |

| | | |
|---|---|---|
| **KQTALVELVK-(453)** | 24 | 24 |
| **KVPQVSTPTLVEVS[453]RNLGK** | 0 | 1 |
| **KVPQVSTPTLVEVSR-(453)** | 1 | 0 |
| **KVPQVSTPTLVEVSRNLG[453]K** | 2 | 4 |
| **L[453]KECCEKPLLEKSHCIAEVENDEM[16]PADLPSLAADFVESK** | 1 | 0 |
| **LAKTYETTLEK-(453)** | 89 | 80 |
| **LDELRDEGK-(453)** | 0 | 1 |
| **LDELRDEGK[453]ASSAK** | 0 | 2 |
| **LDELRDEGKASSAK-(453)** | 71 | 63 |
| **LKC[57]ASLQK-(453)** | 64 | 71 |
| **LKCASLQK-(453)** | 1 | 1 |
| **LVRPEV[453]DVMC[57]TAFHDNEETFLK** | 1 | 0 |
| **MPC[57]AEDYLSVVLNQLC[57]VLHEK-(453)** | 4 | 3 |
| **MPC[57]AEDYLSVVLNQLCVLHEK-(453)** | 3 | 3 |
| **NLGKVGSK-(453)** | 34 | 40 |
| **QNC[57]ELFEQLGEYKFQNALLVR-(453)** | 0 | 1 |
| **RPC[57]F[453]SALEVDETYVPK** | 4 | 2 |
| **RPC[57]FS[453]ALEVDETYVPK** | 6 | 9 |
| **RPC[57]FSA[453]LEVDETYVPK** | 3 | 1 |
| **RPC[57]FSAL[453]EVDETYVPK** | 0 | 1 |
| **RPC[57]FSALEVDETYVPK-(453)** | 7 | 3 |
| **RPCF[453]SALEVDETYVPK** | 0 | 1 |
| **RPCFSALEVDETYVPK-(453)** | 1 | 1 |
| **RYK[453]AAFTEC[57]C[57]QAADK** | 0 | 1 |
| **RYKAAFTEC[57]C[57]QAADK-(453)** | 0 | 2 |
| **RYKAAFTECC[57]QAADK-(453)** | 2 | 1 |
| **RYKAAFTECCQAADK-(453)** | 1 | 1 |
| **SHC[57]IAEVE[453]NDEMPADLPSLAADFVESK** | 1 | 0 |
| **SHC[57]IAEVENDE[453]MPADLPSLAADFVESK** | 1 | 0 |
| **VFDEFKPLVE[453]EPQNLIK** | 2 | 0 |
| **VGSKC[57]C[57]K-(453)** | 22 | 24 |
| **VGSKCCK-(453)** | 0 | 1 |
| **VHTEC[57]C[57]HGDLLEC[57]ADDR-(453)** | 0 | 2 |
| **YKAAFTEC[57]C[57]QAADK-(453)** | 3 | 3 |
| **YKAAFTEC[57]CQAADK-(453)** | 3 | 2 |
| **YKAAFTECCQAADK-(453)** | 4 | 6 |

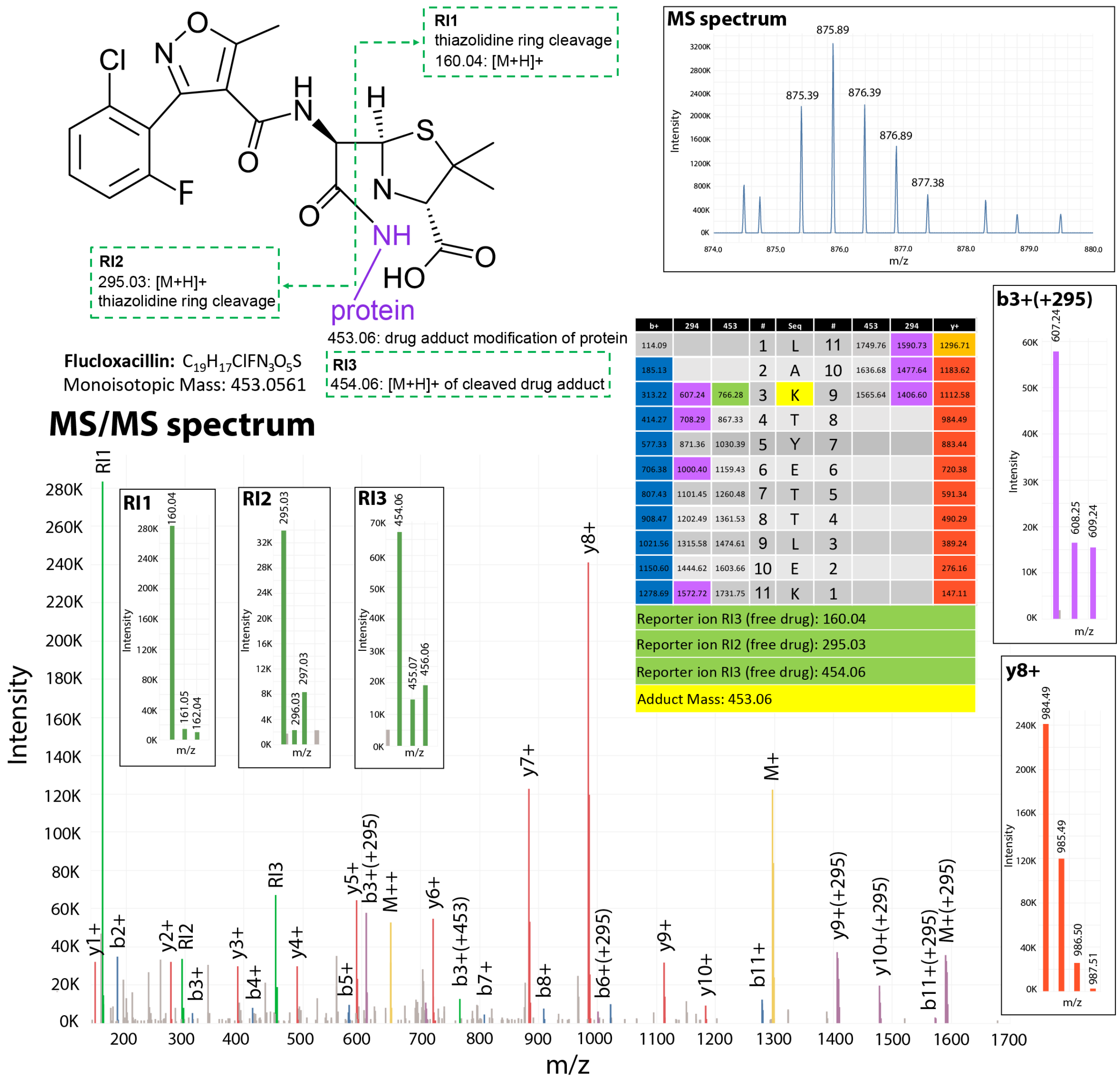

| b+ | 294 | 453 | # | Seq | # | 453 | 294 | y+ |
|---|---|---|---|---|---|---|---|---|
| 114.09 | | | 1 | L | 11 | 1749.76 | 1590.73 | 1296.71 |
| 185.13 | | | 2 | A | 10 | 1636.68 | 1477.64 | 1183.62 |
| 313.22 | 607.24 | 766.28 | 3 | K | 9 | 1565.64 | 1406.60 | 1112.58 |
| 414.27 | 708.29 | 867.33 | 4 | T | 8 | | | 984.49 |
| 577.33 | 871.36 | 1030.39 | 5 | Y | 7 | | | 883.44 |
| 706.38 | 1000.40 | 1159.43 | 6 | E | 6 | | | 720.38 |
| 807.43 | 1101.45 | 1260.48 | 7 | T | 5 | | | 591.34 |
| 908.47 | 1202.49 | 1361.53 | 8 | T | 4 | | | 490.29 |
| 1021.56 | 1315.58 | 1474.61 | 9 | L | 3 | | | 389.24 |
| 1150.60 | 1444.62 | 1603.66 | 10 | E | 2 | | | 276.16 |
| 1278.69 | 1572.72 | 1731.75 | 11 | K | 1 | | | 147.11 |
| Reporter ion RI3 (free drug): 160.04 | | | | | | | | |
| Reporter ion RI2 (free drug): 295.03 | | | | | | | | |
| Reporter ion RI3 (free drug): 454.06 | | | | | | | | |
| Adduct Mass: 453.06 | | | | | | | | |

*Supplementary Figure 12: Annotated MS and MS/MS spectrum of flucloxacillin adducted HSA peptide identified by Magnum. The adduct (structure, top left) is covalently bound to the lysine primary amine (purple). Fluoxacillin is an MS-labile adduct. During peptide fragmentation the adduct itself is cleaved at the thiazolidine ring (green dotted line) releasing reporter ion 1 (RI1) and reporter ion 2 (RI2). The entire adduct can also be cleaved from the peptide releasing reporter ion 3 (RI3). The masses of these ions are independent of the peptide to which the drug is adducted as the ions are derived from adduct fragmentation. For this precursor ion, the MS spectrum indicated an observed mass of 875.39 m/z at charge 2 ($M+2H^{2+}$). The observed precursor mass is thus 1,748.76 Da. The theoretical mass of peptide LAKTYETTLEK is 1,295.697 Da. The adduct modification mass, equal to the mass unexplained by the predicted peptide, is thus 453.063 Da. The peptide sequence and calculated ion series are displayed in the table (center right). The annotated MS/MS spectrum is shown below with inset zoomed panels depicting RI1, RI2, RI3, b3(+295) which is the b3+ ion plus the flucloxacillin adduct minus the thiazolidine ring and is the dominant drug modified ion series, and y8+. Note that the R2, RI3 and b3+(+295) all have clear chlorine isotope signatures due to the chlorine in flucloxacillin. R1 has no chlorine signature as the thiazolidine ring has no chlorine. Also note that the dominant b and y ion series are unmodified as the adduct is cleaved off the peptide during the fragmentation step. This prevents correct localization of the adduct by all open search algorithms tested (discussed in Supplementary Note 5). The original identification can be viewed on limelight here: https://limelight.yeastrc.org/limelight/d/pg/spectrum-viewer/ps/2323/psm/138936595*

### Supplementary Note 5: Localization of dicloxacillin and flucloxacillin adducts

The main manuscript does not discuss the algorithm-determined localization of dicloxacillin and flucloxacillin HSA adducts. This was done because (1) not all algorithms are able to localize adducts and (2) dicloxacillin and flucloxacillin form labile adducts which typically break off during peptide fragmentation leaving unmodified fragment ions. This is illustrated in the main manuscript Figure 5 and Supplementary Figure 12. As a result of the adduct being cleaved from the peptide the unmodified ion series is typically more intense (and thus higher scoring) than the adduct modified ion series. As a result of adduct fragmentation there is generally insufficient spectral evidence to localize dicloxacillin and flucloxacillin adducts accurately. Magnum accounts for both labile and non-labile adducts as described above under Magnum development. In addition, Limelight was designed to simultaneously work with peptides containing both localized and unlocalized open modifications. In the case of labile adducts an unlocalized modification allowing for an unmodified ion series will typically score better than a localized modification within Magnum. For example, 98% of the gold standard PSMs identified by Magnum searches (Supplementary Note 2) scored better as an unlocalized modification even when allowing only unlocalized or lysine localized (the known correct localization) adducts (Supplementary Table 8). If adducts were allowed on any residue, 91% still scored better as an unlocalized modification within Magnum. Adduct localization performed by MSFragger in combination with PTMProphet[23] was incorrect (not on a lysine) 82% of the time, however 74% of the time the adduct was localized to the first residue of the peptide, which results in an unmodified theoretical y ions series. These results do not reflect the localization capabilities of the algorithms tested but reflect the intrinsic properties of labile adducts, which break off during peptide fragmentation. In addition to being able to handle unlocalized and localized modifications, Limelight incorporates the ability to manually move or set the position of any PTM via the spectral viewer while updating the annotated ions in real time. This allows the user to manually evaluate and define the most likely adduct localization on the peptide if desired.

*Supplementary Table 8: Dicloxacillin and flucloxacillin adduct localization determined by Magnum or MSFragger plus PTMProphet in 2,979 gold standard MS/MS spectra. Correct adduct localization is known to be lysine. Magnum (K only) allowed adducts to be unlocalized or lysine localized. Magnum (all localizations) allows adducts to be unlocalized, or, localized to any amino acid in the matched peptide. MSFragger allows adducts to be localized to any amino acid. Results from comet closed searches allowing adduct localization on lysines only are shown as a positive control. Data are shown at 1% FDR.*

| Algorithm | % open mods localized to lysine | % open mods localized to N terminal | % open mods unlocalized |
|---|---|---|---|
| Comet closed (K only) | 100 | 4.10 | n/a |
| Magnum (K or unlocalized) | 1.58 | 0.08 | 98.4 |
| Magnum (all amino acids or unlocalized) | 0.21 | 7.52 | 91.1 |
| MSFragger (all amino acids) | 17.74 | 73.74 | n/a |

#### Supplementary Note 6: CYP3A4/Raloxifene analysis

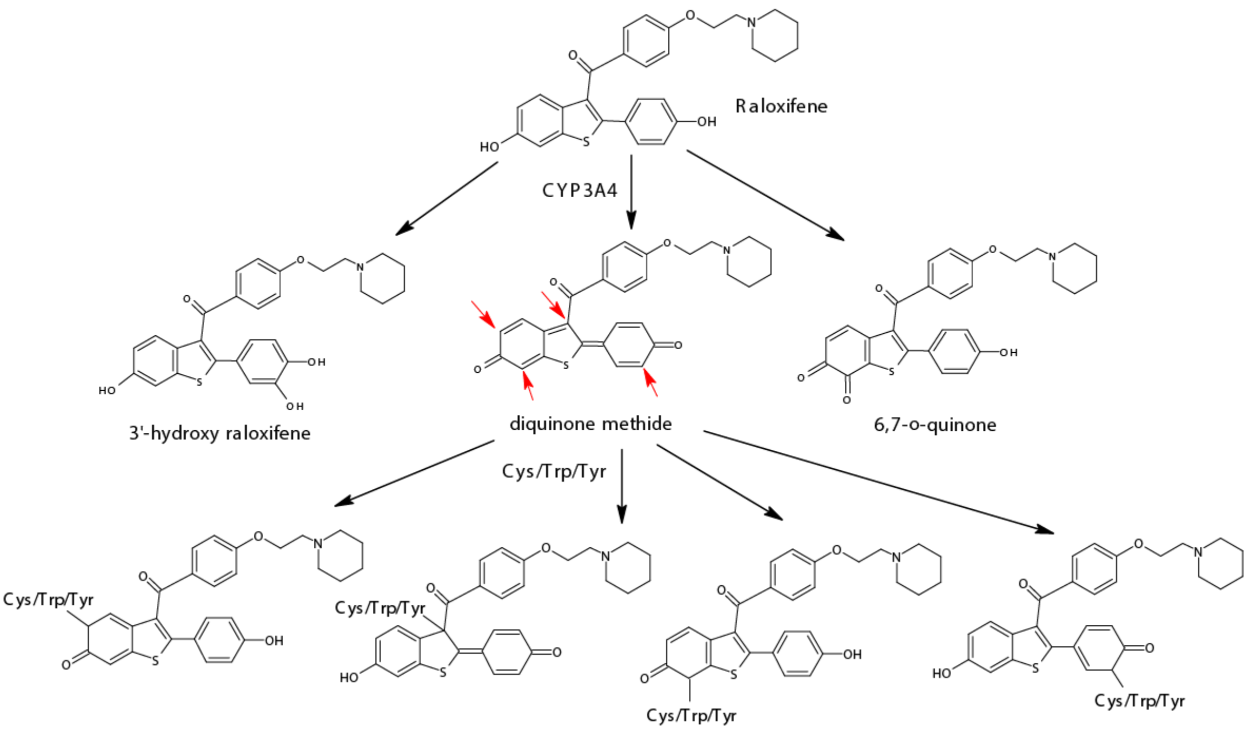

*Supplementary Figure 13: Bioactivation of raloxifene by CYP3A4 and representative adducts formed with cysteine, tryptophan or tyrosine. Red arrows indicate possible sites of diquinone methide which can undergo nucleophilic attack by amino acids to form adducts with a modification mass of 471 Da[27].*

##### Initial Experiments and Skyline Quantification

LC-MS/MS data of raloxifene treated and untreated purified CYP3A4 plus P450-reductase were acquired, searched using Magnum, and analyzed using Limelight. Comparisons of individual raloxifene treated versus untreated replicates, using a two-tailed tests of proportions, resulted in several masses reported as significantly enriched in either untreated or treated samples.

To decrease the significance of masses that varied between replicates while increasing the significance of masses that were constant across replicates, we developed a method able to combine multiple replicates into treatment groups within Limelight (see Experiment Builder, Supplementary Note 3). A two-tailed test of proportions analyzing Magnum data on untreated versus raloxifene treated groups resulted in 471 Da being the most significantly enriched mass in the raloxifene treated sample group (Supplementary Table 9).

*Supplementary Table 9: A two-tailed test of proportions identifies a 471 Da raloxifene specific adduct mass in CYP3A4 and P450-reductase. PSMs were generated using Magnum and analyzed using Limelight. Results were sorted on the absolute value of the Z score (large to small) followed by the magnitude of the P value (small to large). The top 5 masses are shown, representing the most significantly enriched masses found in either the treated sample groups (negative Z scores) or the untreated sample group (positive Z scores). Magnum was run allowing for open masses on any amino acid. Data are shown at 1% FDR. Full data is available in Supplementary File raloxifene_twotail.xlsx and a live view the data is available at: https://limelight.yeastrc.org/limelight/go/wwa741ZTTE (click [View Replicate ZScore Report] to view table).*

| mod mass | untreated PSM count | treated PSM count | z-score | p-value |
|---|---|---|---|---|
| **471** | **14** | **146** | **-10.485762** | **0** |
| 293 | 367 | 183 | 7.78277453 | 3.13349E-12 |
| 298 | 594 | 375 | 6.95595554 | 1.54427E-09 |
| 72 | 490 | 306 | 6.44581951 | 5.07051E-08 |
| 115 | 174 | 79 | 5.92416284 | 1.38427E-06 |

We searched the same MS data using 6 other open search algorithms. PSMs were imported into Limelight and a two-tailed test of proportions comparing untreated versus raloxifene treated sample groups was performed with PSMs generated by each algorithm. Of the other algorithms used, MetaMorpheus also found 471 Da as being the most significantly enriched mass in the raloxifene treated sample group (Supplementary Figure 14a). However, Magnum identified twice as many PSMs with that modification mass in treated samples than MetaMorpheus (146 versus 72) and fewer 471 Da masses in untreated samples (14 versus 16). The treated:untreated ratio of 471 Da PSMs for MetaMorpheus was 5:1 indicating about 80% of the 72 PSMs with a 471 Da modification in treated samples were treatment specific. In contrast an equivalent Magnum search had a 10:1 ratio as described in the main manuscript. Other algorithms identified a range of masses as being significantly enriched in treated or untreated samples, however there were typically many PSMs for each of these masses in both treated and untreated samples and the resulting data did not indicate clear treatment specific adduct masses. For example, Magnum and MSFragger reported a similar number of PSMs containing 471 Da modification masses in treated samples, however 471 Da modification masses were also common in PSMs identified by MSFragger in untreated samples leading to a treated to untreated ratio of 3:1. This suggests that 1 in 3 PSMs containing a 471 Da modification mass identified by MSFragger were likely unrelated to exposure. For this experiment, therefore, the advantages of Magnum, which was designed specifically for xenobiotic protein-adduct detection, were required to detect and distinguish raloxifene adducts in a background of other masses. A complete summary of results from all algorithms and links to all data in Limelight can be found in Supplementary File raloxifene_twotail.xlsx.

(a)

| Search Algorithm | Test Rank | Mod Mass (Da) | PSM count (untreated) | PSM count (treated) | z-score | p-value |
|---|---|---|---|---|---|---|
| Comet Wide | 1 | 88 | 137 | 386 | -11.4 | 0 |
| Comet-PTM | 1 | 88 | 160 | 369 | -9.70 | 0 |
| Magnum | 1 | 471 | 14 | 146 | -10.5 | 0 |
| Magnum C | 1 | 471 | 5 | 67 | -7.50 | 2.4E-11 |
| Magnum CWY | 1 | 471 | 3 | 157 | -12.362 | 0 |
| Meta-Morpheus | 3 | 471 | 16 | 72 | -6.08 | 5.4E-07 |
| MODa | 6 | 248 | 9 | 34 | -3.87 | 0.04 |
| MSFragger | 6 | 88 | 126 | 281 | -7.16 | 3.7E-10 |
| open-pFind | n/a | n/a | n/a | n/a | n/a | n/a |

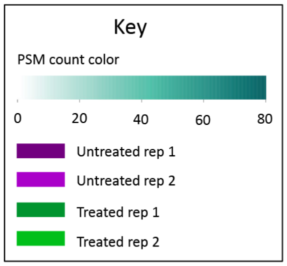

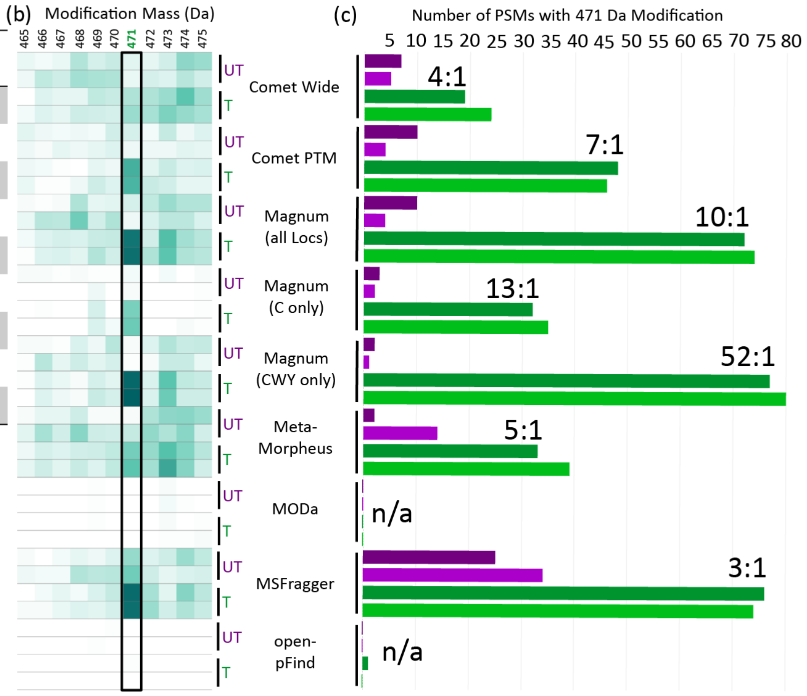

*Supplementary Figure 14: Identification of novel raloxifene adducts in CYP3A4 and P450-reductase. (a) A two-tailed test of proportions performed within Limelight on treatment groups from PSMs identified by 7 open search algorithms. Results were sorted on the absolute value of the Z score (large to small) followed by the magnitude of the P value (small to large). The top significant (p≤0.05) negative Z score is shown for each algorithm, representing the most significantly enriched mass found by each algorithm in the raloxifene treated samples. Magnum was run allowing for open masses on any amino acid (Magnum), restricted to cysteines only (Magnum C), or restricted to cysteine, tryptophan, and tyrosine (Magnum CWY). Data are shown at 1% FDR. Full data is available in Supplementary File raloxifene_twotail.xlsx and a live view of (b) is available at: https://limelight.yeastrc.org/limelight/go/N8sMlKwtQm.*

All 146 PSMs containing a 471 Da mass returned by a fully unrestricted (modifications allowed on any amino acid) Magnum search were manually evaluated (Supplementary File raloxifene_Manual_Eval.xlsx) and it was observed that close to 90% of 471 Da adducts were on cysteine, tryptophan or tyrosine. Further Magnum searches were thus performed restricting open modifications to cysteines (C), tryptophan (W), and tyrosine (Y).

Supplementary Table 10 shows the initial set of residues identified by Magnum-CWY searches as having a 471 Da modification in ≥ 2 PSMs. The number of PSMs at each location is recorded as well as the ion signal quantified in in Skyline (see below). Full data plus Limelight links can be found in Supplementary File raloxifene-MagnumCWY.xlsx (InitialReplicates tab).

Supplementary Table 10: All locations identified as modified by 471 Da adduct masses by Magnum-CWY in CYP3A4 and P450-reductase in initial experiments by ≥ 2 PSMs. PSM counting and Skyline quantification of extracted ion chromatograms were performed to estimate the degree of modification at each location. Full data is available in Supplementary File raloxifene_Manual_Eval.xlsx) and data is plotted in Supplementary Figures 16 and 17 below. Data is shown at a Percolator calculated PSM level q≤0.01.

| Protein Name | PDB numbering modification location | Fasta numbering modification location | Total PSMs at that location | Skyline Quantified Ion Signal at that location |
|---|---|---|---|---|
| **CYP3A4** | Y53 | Y43 | 2 | 0.0000111848 |
| **CYP3A4** | C58 | C48 | 7 | 0.000408657 |
| **CYP3A4** | Y75 | Y65 | 3 | 0.0000385597 |
| **CYP3A4** | C98 | C88 | 20 | 0.001456899 |
| **CYP3A4** | W126 | W116 | 65 | 0.003233199 |
| **CYP3A4** | Y152 | Y142 | 8 | 0.000158244 |
| **CYP3A4** | Y430 | Y420 | 2 | 0.0000243249 |
| **CYP3A4** | Y432 | Y422 | 8 | 0.000229608 |
| **CYP3A4** | C468 | C458 | 19 | 0.001078295 |
| **Reductase** | C472 | C472 | 8 | 0.000118627 |
| **Reductase** | C630 | C630 | 4 | 0.000186243 |

Data were quantified by spectral counting and additionally, extracted ion chromatograms (XICs) were produced and quantified in Skyline[24–26]. Full Skyline sessions are available on Panorama here: https://panoramaweb.org/CYP3A4-raloxifene.url. Skyline XICs showed treatment specific signal for all peptides identified by Magnum as exclusively in treated samples. Only one peptide, VWGFYDGQQPVLAITDPDM[+16]IKTVLVKEC[+471.2]YSVFTNR, was found to be unrelated to raloxifene treatment. This peptide, identified in a single untreated PSM, was found by both PSM counting and Skyline quantification to be unrelated to raloxifene treatment (Supplementary Figure 15) and was included as the single non-raloxifene specific peptide had the same 471 Da adduct location (C98) as multiple treatment-specific identifications in separate peptides. All other peptides showed treatment specific 471 Da modifications.

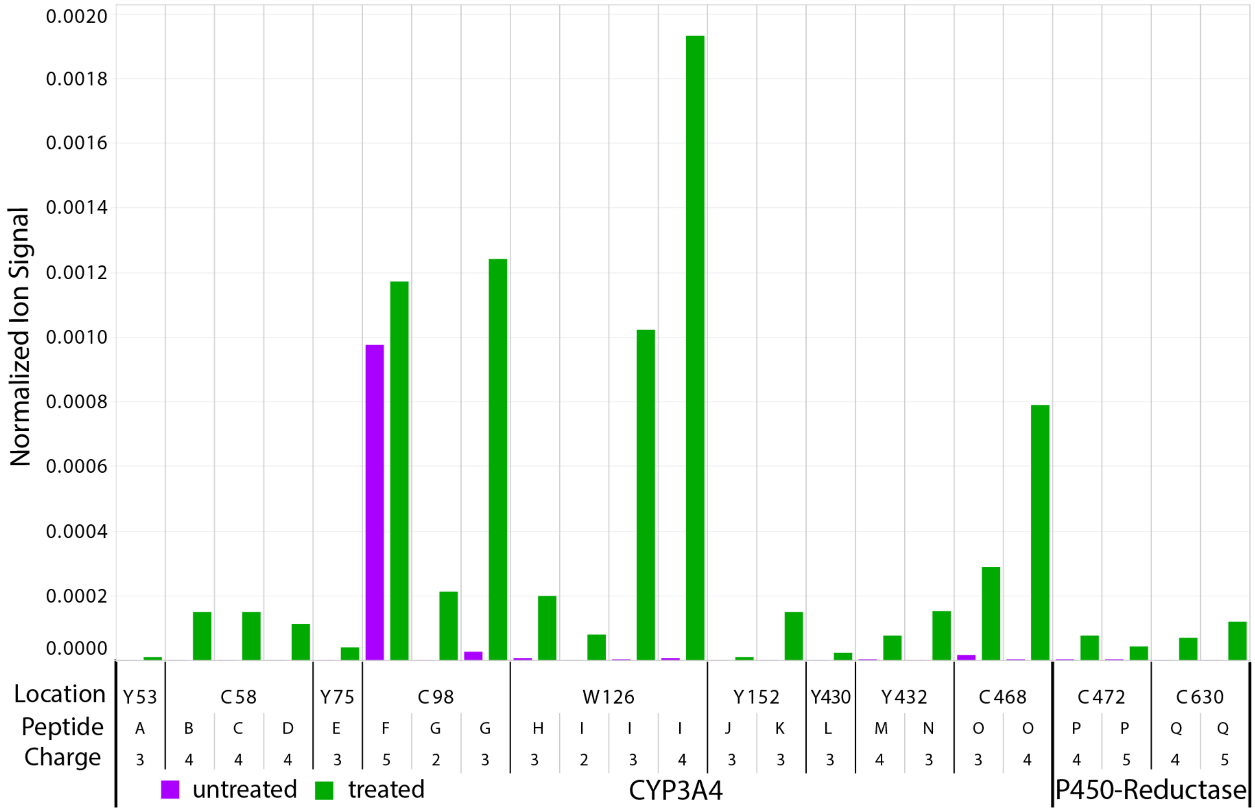

Supplementary Figure 15: Total normalized ion signal in treated and untreated samples, quantified in Skyline, for individual precursor ions corresponding to peptides identified in CYP3A4 and P450-reductase at each location found by Magnum-CWY searches as modified by a 471 Da adduct mass in initial experiments by ≥ 2 PSMs. Full data are available in Supplementary File raloxifene-MagnumCWY.xlsx, SkylineQuant tab. Full Skyline sessions are available on Panorama here: https://panoramaweb.org/CYP3A4-raloxifene.url. Note that peptide F (VWGFYDGQQPVLAITDPDM[+16]IKTVLVKEC[471]YSVFTNR) is not expected to be treatment specific and is only included as the residue (C98) predicted to be modified by 471 Da in one untreated 1 PSM was also identified as modified by 471 Da in 20 other treatment specific PSMs in a different peptide (peptide G: EC[471]YSVFTNR) covering the same residue. The modifications in the other peptide are clearly treatment specific. Supplementary Table 11 shows the full peptide sequence corresponding to the peptide letter abbreviations used in this figure.

Supplementary Table 11: Full peptide sequences corresponding to the peptide letter abbreviations used in Supplementary Figure 15.

| Protein | Peptide Modified Sequence | Peptide | Precursor Charge | PDB numbering modification location | Fasta numbering modification location |
|---|---|---|---|---|---|
| **CYP3A4** | LGIPGPTPLPFLGNILSY[+471.2]HK | A | 3 | 53 | 43 |
| **CYP3A4** | GFC[+471.2]M[+16]FDMEC[+57]HKK | B | 4 | 58 | 48 |
| **CYP3A4** | GFC[+471.2]MFDM[+16]EC[+57]HKK | C | 4 | 58 | 48 |
| **CYP3A4** | GFC[+471.2]M[+16]FDM[+16]EC[+57]HKK | D | 4 | 58 | 48 |
| **CYP3A4** | VWGFY[+471.2]DGQQPVLAITDPDM[+16]IK | E | 3 | 75 | 65 |
| **CYP3A4** | VWGFYDGQQPVLAITDPDM[+16]IKTVLVKEC[+471.2]YSVFTNR | F | 5 | 98 | 88 |
| **CYP3A4** | EC[+471.2]YSVFTNR | G | 2 | 98 | 88 |
| **CYP3A4** | EC[+471.2]YSVFTNR | G | 3 | 98 | 88 |
| **CYP3A4** | SAISIAEDEEW[+471.2]K | H | 3 | 126 | 116 |
| **CYP3A4** | SAISIAEDEEW[+471.2]KR | I | 2 | 126 | 116 |
| **CYP3A4** | SAISIAEDEEW[+471.2]KR | I | 3 | 126 | 116 |
| **CYP3A4** | SAISIAEDEEW[+471.2]KR | I | 4 | 126 | 116 |
| **CYP3A4** | EMVPIIAQY[+471.2]GDVLVR | J | 3 | 152 | 142 |
| **CYP3A4** | EM[+16]VPIIAQY[+471.2]GDVLVR | K | 3 | 152 | 142 |
| **CYP3A4** | DNIDPY[+471.2]IYTPFGSGPR | L | 3 | 430 | 420 |
| **CYP3A4** | NKDNIDPYIY[+471.2]TPFGSGPR | M | 4 | 432 | 422 |
| **CYP3A4** | DNIDPYIY[+471.2]TPFGSGPR | N | 3 | 432 | 422 |
| **CYP3A4** | VLQNFSFKPC[+471.2]K | O | 3 | 468 | 458 |
| **CYP3A4** | VLQNFSFKPC[+471.2]K | O | 4 | 468 | 458 |
| **P450-Reductase** | VHPNSVHIC[+471.2]AVAVEYEAK | P | 4 | 472 | 472 |
| **P450-Reductase** | VHPNSVHIC[+471.2]AVAVEYEAK | P | 5 | 472 | 472 |
| **P450-Reductase** | LIHEGGAHIYVC[+471.2]GDAR | Q | 4 | 630 | 630 |
| **P450-Reductase** | LIHEGGAHIYVC[+471.2]GDAR | Q | 5 | 630 | 630 |

Quantification of 471 Da modifications by spectral counting (Supplementary Figure 16) and by Skyline quantification of XICs (Supplementary Figure 17) produced similar results and thus spectral counting was performed when combining data from all raloxifene adduct experiments.

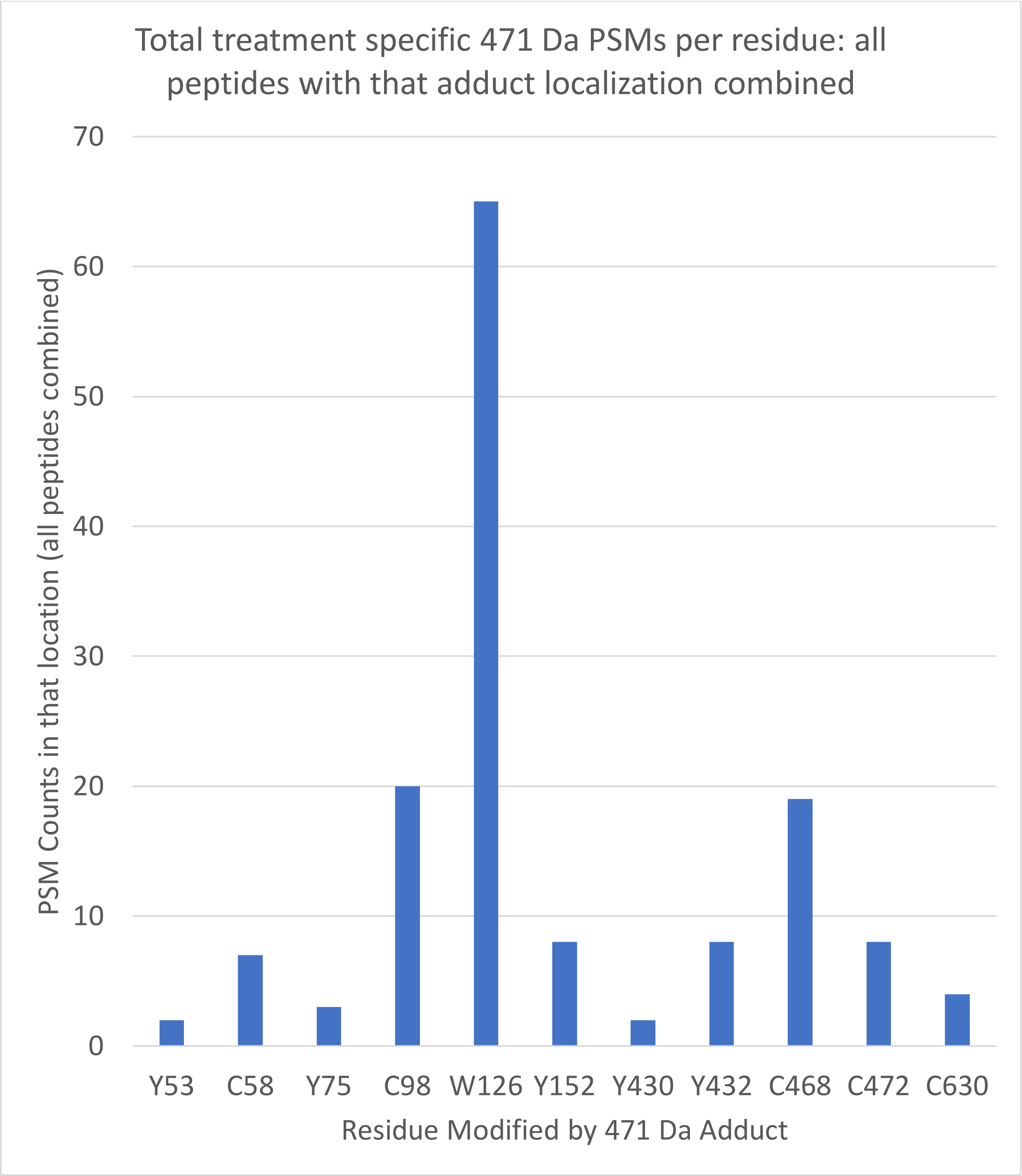

*Supplementary Figure 16: Total PSMs identified in CYP3A4 and P450-reductase at each location found by Magnum-CWY searches as modified by a 471 Da adduct mass in initial experiments by ≥ 2 PSMs. Data is shown at a Percolator calculated PSM level q≤0.01. Full data is available in Supplementary File raloxifene-MagnumCWY.xlsx, InitialReplicates tab.*

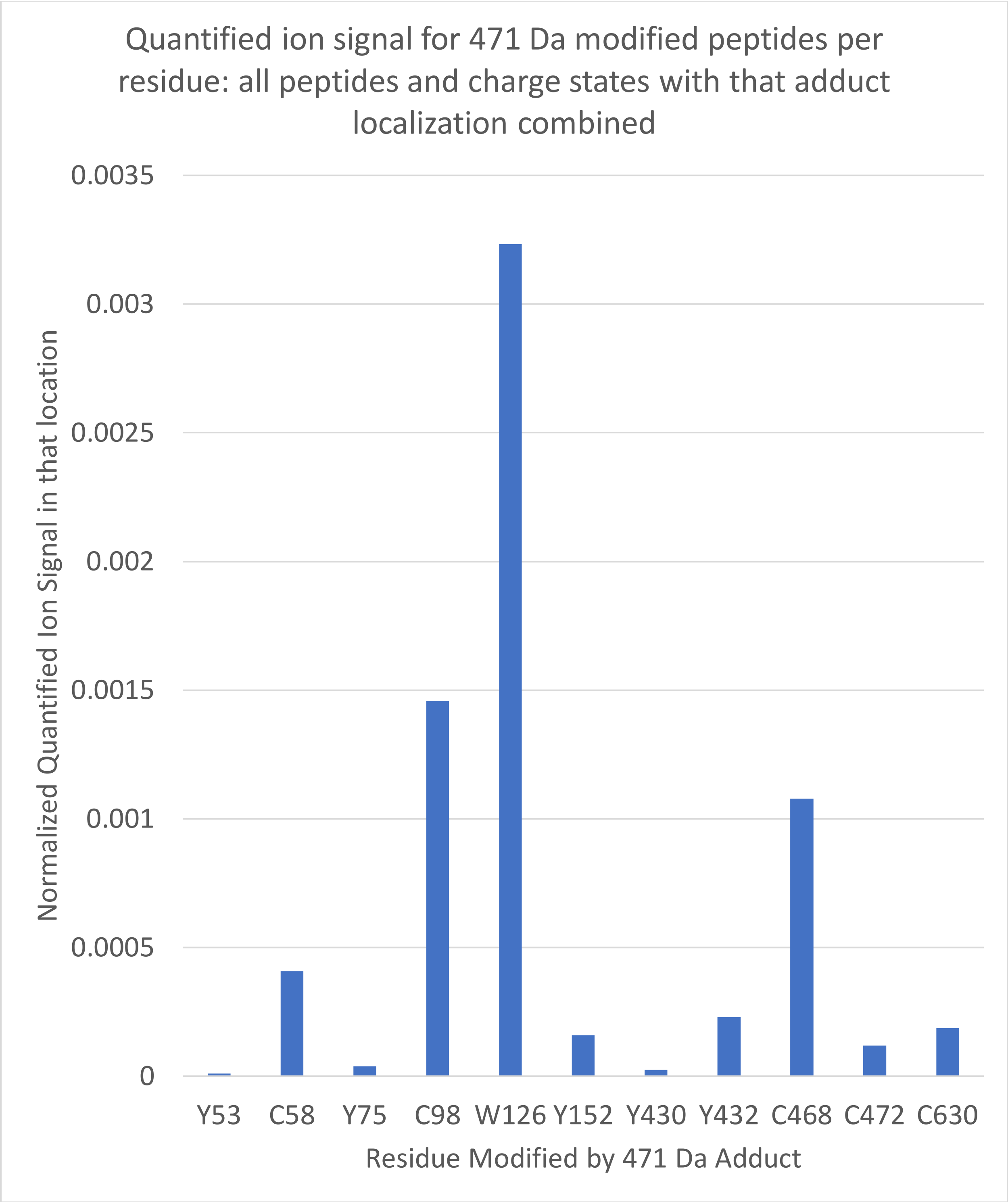

*Supplementary Figure 17: Total normalized ion signal quantified in Skyline for all locations identified in CYP3A4 and P450-reductase at each location found by Magnum-CWY searches as modified by a 471 Da adduct mass in initial experiments by ≥ 2 PSMs. Full data is available in Supplementary File raloxifene-MagnumCWY.xlsx, SkylineQuant tab. Full Skyline sessions are available on Panorama here: [https://panoramaweb.org/CYP3A4-raloxifene.url](https://panoramaweb.org/CYP3A4-raloxifene.url). The final results, combining all raloxifene replicates, are presented in Figure 6 in the main manuscript and are presented in*

*tabular form below (Supplementary Table 12). The complete analysis is available in Supplementary File raloxifene-MagnumCWY.xlsx.*

#### Several raloxifene adducts result in multiple distinct chromatographic peaks

During quantification of raloxifene specific precursor ions, we observed that several raloxifene (471 Da) adducted peptides were associated with distinct chromatographic peaks. The three most obvious examples were W126, C98 and C468, which were also the ions with the largest signals (Supplementary Figures 18-20). The following CYP3A4 peptides were observed to produce >1 chromatographic peak: K.ECYSVFTNR.R (3 peaks); K.SAISIAEDEEWKR.L (4 peaks); K.NKDNIDPYIYTPFGSGPR.N (2 peaks); K.DNIDPYIYTPFGSGPR.N (2 peaks); K.DNIDPYIYTPFGSGPR.N (2 peaks); R.VLQNFSFKPCK.E (2 peaks). P450-reductase peptide K.LIHEGGAHIYCCGDAR.N also produced 2 peaks. All other peptides (e.g. Supplementary Figure 21) produced 1 chromatographic peak under the experimental conditions used.

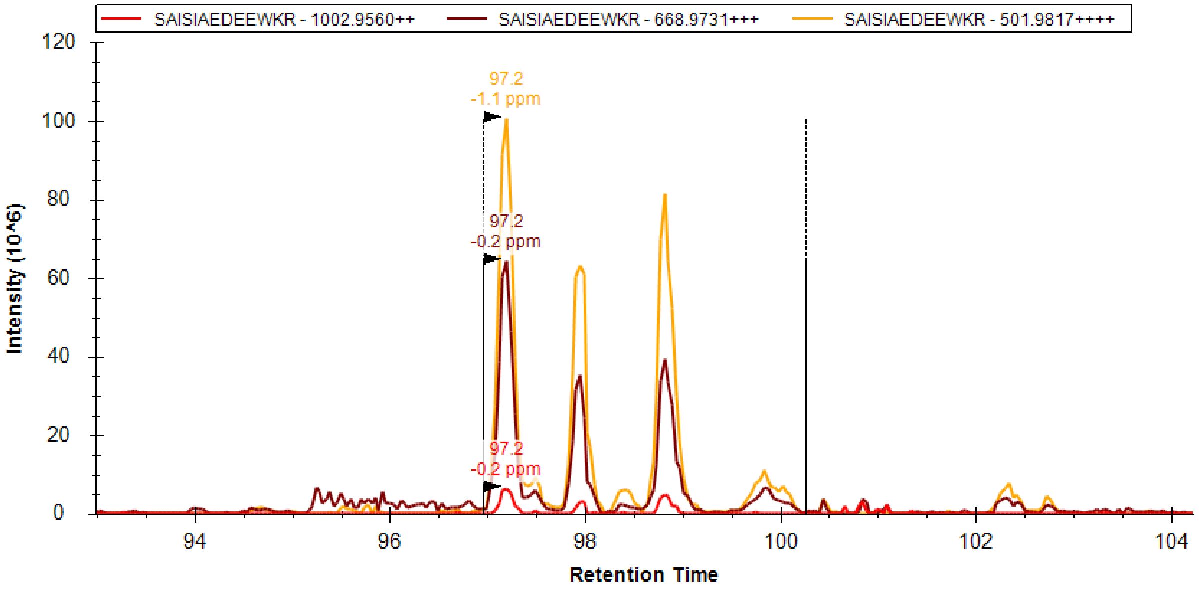

*Supplementary Figure 18: Precursor ions resulting corresponding to the W126 peptide (SAISIAEDEEW[471]KR) elute as 4 distinct chromatographic peaks. Skyline derived extracted ion chromatograms are shown of 2+, 3+ and 4+ precursor ions. Full skyline sessions are available on Panorama here:* [https://panoramaweb.org/CYP3A4-raloxifene.url](https://panoramaweb.org/CYP3A4-raloxifene.url)

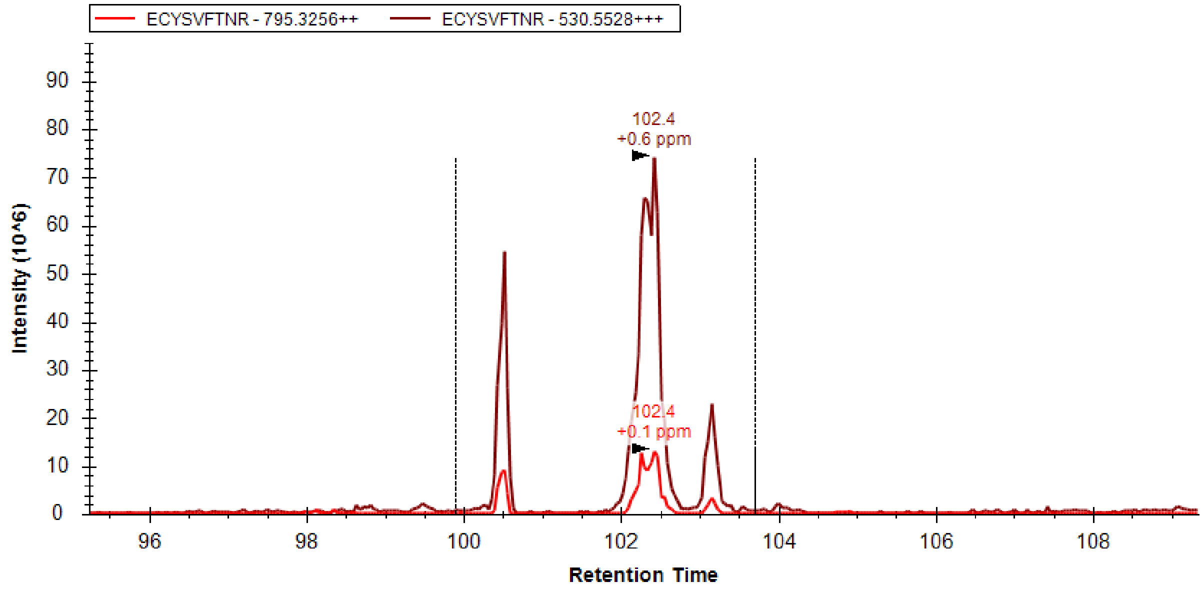

*Supplementary Figure 19: Precursor ions resulting corresponding to the C98 peptide (EC[471]YSVFTNR) elute as 3 distinct chromatographic peaks. Skyline derived extracted ion chromatograms are shown of 2+ and 3+ precursor ions. Full skyline sessions are available on Panorama here: [https://panoramaweb.org/CYP3A4-raloxifene.url](https://panoramaweb.org/CYP3A4-raloxifene.url)*

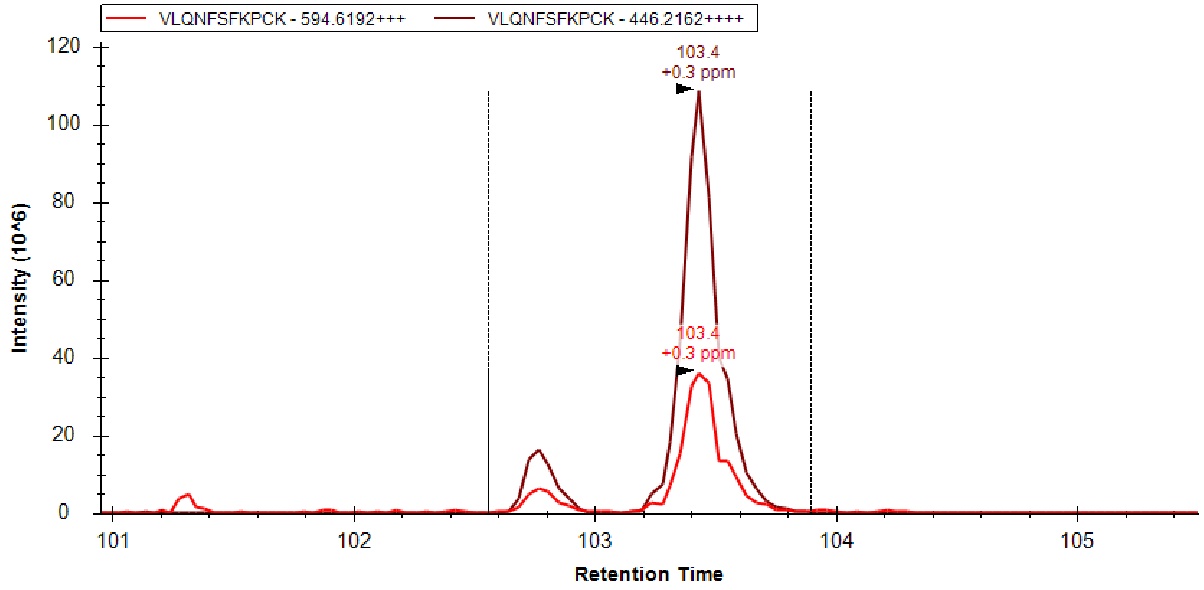

*Supplementary Figure 20: Precursor ions resulting corresponding to the C468 peptide (VLQNFSFKPC[471]K) elute as 2 distinct chromatographic peaks. Skyline derived extracted ion chromatograms are shown of 3+ and 4+ precursor ions. Full skyline sessions are available on Panorama here: [https://panoramaweb.org/CYP3A4-raloxifene.url](https://panoramaweb.org/CYP3A4-raloxifene.url)*

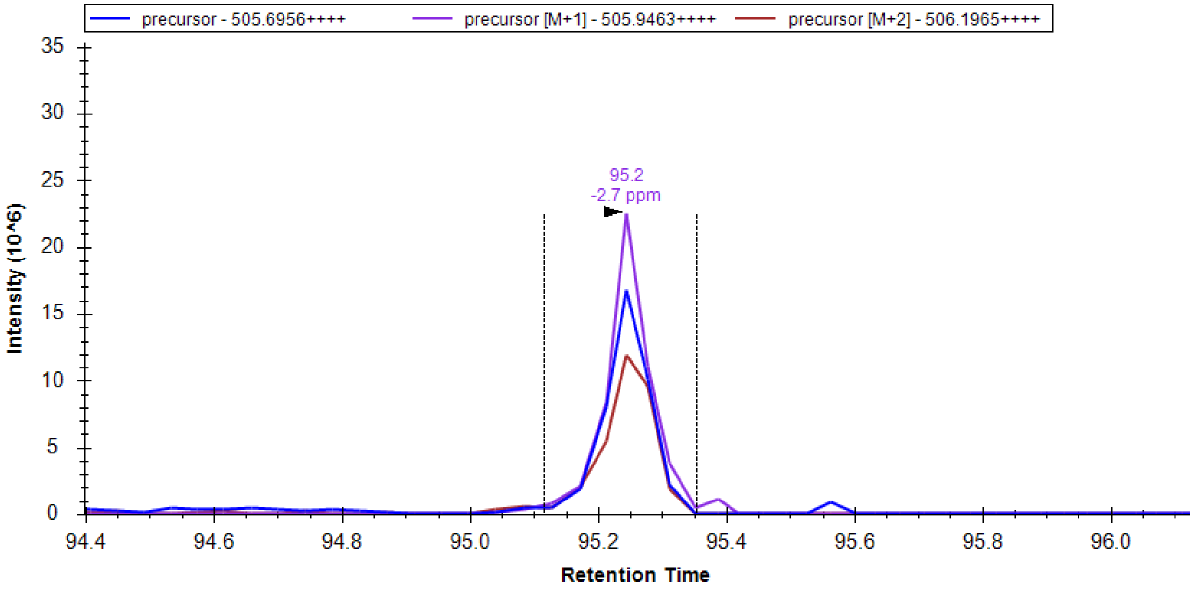

*Supplementary Figure 21: Precursor ions resulting corresponding to the C58 peptide (GFC[471]M[16]FDMEC[57]HKK) elute as 1 single distinct chromatographic peak. Skyline derived extracted ion chromatograms are shown of the 4+ precursor ion. Full skyline sessions are available on Panorama here:* https://panoramaweb.org/CYP3A4-raloxifene.url

Previously published work[27] identified 4 different positions in the raloxifene metabolite, diquinone methide, subject to nucleophilic attack and the presence of multiple distinct chromatographic peaks could be explained by adducts on these differing positions (Supplementary Figure 13).

#### Further raloxifene experiments

To increase peptide the depth of our CYP3A4/raloxifene analysis we performed two additional sets of MS analyses on our raloxifene treated and untreated purified CYP3A4 plus P450-reductase samples. Firstly, a longer tryptic digest was performed (see “extra digest” in Materials and Methods) and secondly a higher acetonitrile LC gradient was used for select CYP3A4 samples (see “highB” in Materials and Methods). Results from all CYP3A4 plus P450-reductase experiments combined are presented below.

*Supplementary Table 12: Magnum identifies multiple 471 Da protein adducts in CYP3A4 and P450-reductase after exposure to raloxifene. Abundance of all modifications is shown relative to W126, which was observed in 121 PSMs. All results shown have a Percolator calculated PSM level q≤0.01 and were identified by ≥3 PSMs. Raw and data and visualizations are available on Limelight at: https://limelight.yeastrc.org/limelight/go/0UjwIJNz45 (CYP3A4) and https://limelight.yeastrc.org/limelight/go/ypAqoCB3IE (reductase).*

| Protein | PDB numbering modification location | Fasta numbering modification location (used in Limelight) | Total treatment specific PSMs in that location | Relative Abundance |
|---|---|---|---|---|
| CYP3A4 | **Y53** | 43 | 18 | 0.14876 |
| CYP3A4 | **C58** | 48 | 8 | 0.06612 |
| CYP3A4 | **Y75** | 65 | 9 | 0.07438 |
| CYP3A4 | **C98** | 88 | 30 | 0.24793 |
| CYP3A4 | **W126** | 116 | 121 | 1 |
| CYP3A4 | **Y152** | 142 | 76 | 0.6281 |
| CYP3A4 | **Y399** | 389 | 40 | 0.33058 |
| CYP3A4 | **Y407** | 397 | 23 | 0.19008 |
| CYP3A4 | **Y430** | 420 | 21 | 0.17355 |
| CYP3A4 | **Y432** | 422 | 25 | 0.20661 |
| CYP3A4 | **C442** | 432 | 3 | 0.02479 |
| CYP3A4 | **C468** | 458 | 49 | 0.40496 |
| Reductase | **Y84** | 84 | 11 | 0.09091 |
| Reductase | **Y178** | 178 | 7 | 0.05785 |
| Reductase | **Y259** | 259 | 4 | 0.03306 |
| Reductase | **Y269** | 269 | 13 | 0.10744 |
| Reductase | **Y373** | 373 | 13 | 0.10744 |
| Reductase | **Y374** | 374 | 9 | 0.07438 |
| Reductase | **Y387** | 387 | 5 | 0.04132 |
| Reductase | **C472** | 472 | 8 | 0.06612 |
| Reductase | **Y564** | 564 | 3 | 0.02479 |
| Reductase | **C630** | 630 | 3 | 0.02479 |
| Reductase | **Y672** | 672 | 8 | 0.06612 |

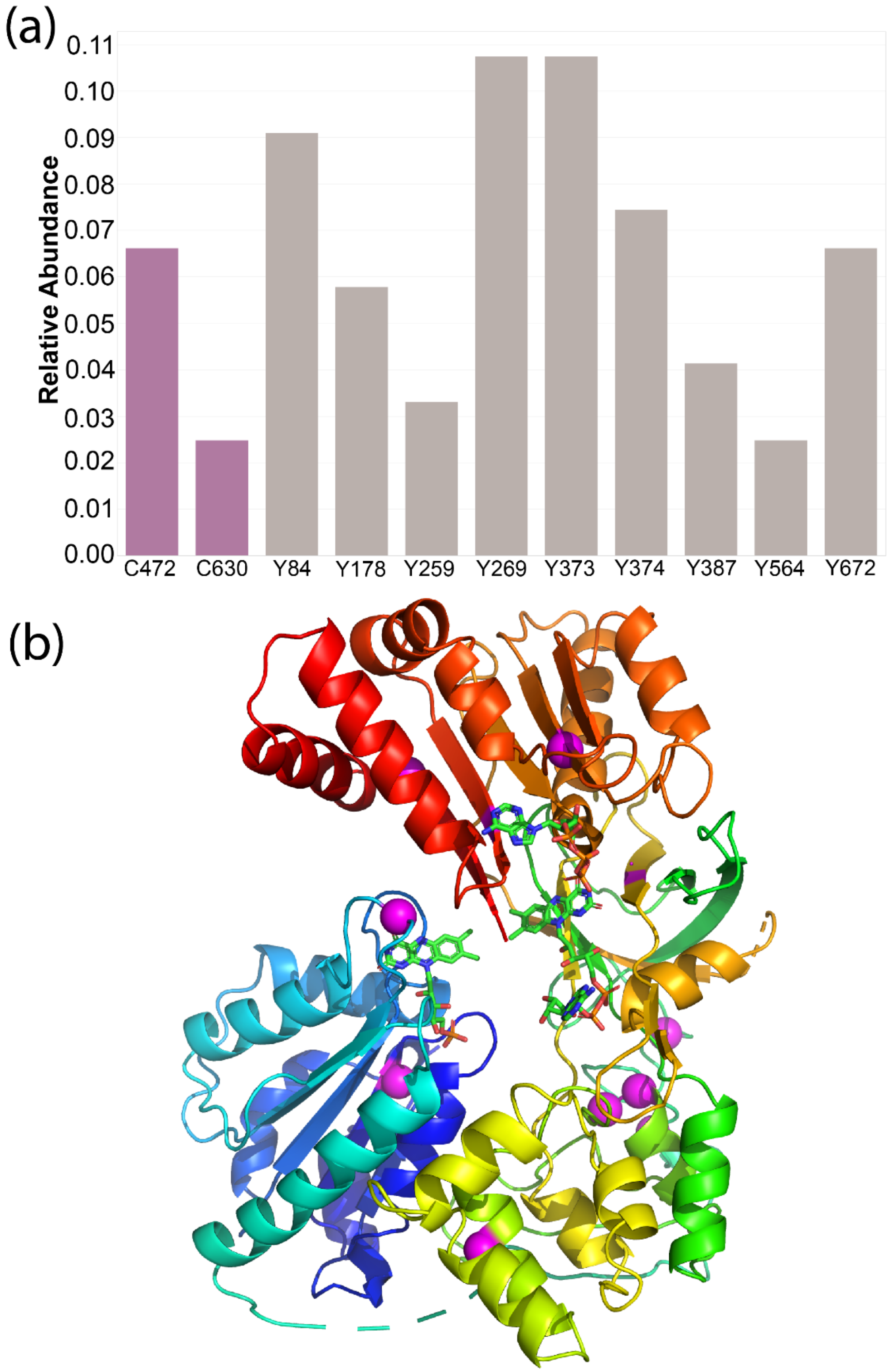

*Supplementary Figure 22: Identification of novel raloxifene adducts in P450-reductase. (a) Abundance of all modifications is shown based on the number of PSMs identified in each location relative to CYP3A4 W126, which was observed in 121 PSMs (main manuscript Figure 6, Supplementary Table 12 above). (b) Observed 471 Da modifications are shown on the structure of P450-reductase (PDB: 1AMO) as magenta spheres. The size of the magenta spheres is proportional to the relative abundance reported in (a). Supplemental File 1AMO-raloxifene.pse is a pymol session of this image. All results shown have a Percolator calculated PSM level q≤0.01 and were identified by ≥3 PSMs. Raw data are available on Limelight at: [https://limelight.yeastrc.org/limelight/go/ypAqoCB3IE](https://limelight.yeastrc.org/limelight/go/ypAqoCB3IE)*

### Key protein sequences:

Supplementary Table 13: Protein sequences of human serum albumin (HSA) plus the heterologously expressed proteins CYP3A4 and rat P450 reductase proteins.

```
>sp|P02768|ALBU_HUMAN Serum albumin OS=Homo sapiens GN=ALB PE=1 SV=2
MKWVTFISLLFLFSSAYSRGVFRRDAHKSEVAHRFKDLGEENFKALVLIAFAQYLQQCPFEDHVKLVNEVTEFAKTCVADESAENC
DKSLHTLFGDKLCTVATLRETYGEMADCCAKQEPERNECFLQHKDDNPNLPRLVRPEVDVMCTAFHDNEETFLKKYLYEIARRHPY
FYAPELLFFAKRYKAAFTECCQAADKAACLLPKLDELRDEGKASSAKQRLKCASLQKFGERAFKAWAVARLSQRFPKAEFAEVSKL
VTDLTKVHTECCHGDLLECADDRADLAKYICENQDSISSKLKECCEKPLLEKSHCIAEVENDEMPADLPSLAADFVESKDVCKNYA
EAKDVFLGMFLYEYARRHPDYSVVLLLRLAKTYETTLEKCCAAADPHECYAKVFDEFKPLVEEPQNLIKQNCELFEQLGEYKFQNA
LLVRYTKKVPQVSTPTLVEVSRNLGKVGSKCCKHPEAKRMPCAEDYLSVVLNQLCVLHEKTPVSDRVTKCCTESLVNRRPCFSALE
VDETYVPKEFNAETFTFHADICTLSEKERQIKKQTALVELVKHKPKATKEQLKAVMDDFAAFVEKCCKADDKETCFAEEGKKLVAA
SQAALGL

>cu|hCYP3A4pCWOri|hCYP3A4pCWOri Cytochrome P450 3A4 cloned in pCWOri OS=Homo sapiens
OX=9606
MALLLAVFLVLLYLYGTHSHGLFKKLGIPGPTPLPFLGNILSYHKGFCMFDMECHKKYGKVWGFYDGQQPVLAITDPDMIKTVLVK
ECYSVFTNRRPFGPVGFMKSAISIAEDEEWKRLRSLLSPTFTSGKLKEMVPIIAQYGDVLVRNLRREAETGKPVTLKDVFGAYSMD
VITSTSFGVNIDSLNNPQDPFVENTKKLLRFDFLDPFFLSITVFPFLIPILEVLNICVFPREVTNFLRKSVKRMKESRLEDTQKHR
VDFLQLMIDSQNSKETESHKALSDLELVAQSIIFIFAGYETTSSVLSFIMYELATHPDVQQKLQEEIDAVLPNKAPPTYDTVLQME
YLDMVVNETLRLFPIAMRLERVCKKDVEINGMFIPKGVVVMIPSYALHRDPKYWTEPEKFLPERFSKKNKDNIDPYIYTPFGSGPR
NCIGMRFALMNMKLALIRVLQNFSFKPCKETQIPLKLSLGGLLQPEKPVVLKVESRDGTVSGASTHHHHHH

>sp|P00388|NCPR_RAT NADPH--cytochrome P450 reductase OS=Rattus norvegicus OX=10116
GN=Por PE=1 SV=3
MGDSHEDTSATMPEAVAEEVSLFSTTDMVLFSLIVGVLTYWFIFRKKKEEIPEFSKIQTTAPPVKESSFVEKMKKTGRNIIVFYGS
QTGTAEEFANRLSKDAHRYGMRGMSADPEEYDLADLSSLPEIDKSLVVFCMATYGEGDPTDNAQDFYDWLQETDVDLTGVKFAVFG
LGNKTYEHFNAMGKYVDQRLEQLGAQRIFELGLGDDDGNLEEDFITWREQFWPAVCEFFGVEATGEESSIRQYELVVHEDMDVAKV
YTGEMGRLKSYENQKPPFDAKNPFLAAVTANRKLNQGTERHLMHLELDISDSKIRYESGDHVAVYPANDSALVNQIGEILGADLDV
IMSLNNLDEESNKKHPFPCPTTYRTALTYYLDITNPPRTNVLYELAQYASEPSEQEHLHKMASSSGEGKELYLSWVVEARRHILAI
LQDYPSLRPPIDHLCELLPRLQARYYSIASSSKVHPNSVHICAVAVEYEAKSGRVNKGVATSWLRAKEPAGENGGRALVPMFVRKS
QFRLPFKSTTPVIMVGPGTGIAPFMGFIQERAWLREQGKEVGETLLYYGCRRSDEDYLYREELARFHKDGALTQLNVAFSREQAHK
VYVQHLLKRDREHLWKLIHEGGAHIYVCGDARNMAKDVQNTFYDIVAEFGPMEHTQAVDYVKKLMTKGRYSLDVWS
```

#### Supplementary References:

1. Keller, A., Eng, J., Zhang, N., Li, X. jun & Aebersold, R. A uniform proteomics MS/MS analysis platform utilizing open XML file formats. *Mol. Syst. Biol.* **1**, (2005).

2. Eng, J. K. *et al.* A Deeper Look into Comet — Implementation and Features. 1865–1874 (2015) doi:10.1007/s13361-015-1179-x.

3. Hoopmann, M. R. *et al.* Kojak: Efficient analysis of chemically cross-linked protein complexes. *J. Proteome Res.* **14**, 2190−2198 (2015).

4. Eng, J. K., Jahan, T. A. & Hoopmann, M. R. Comet: an open-source MS/MS sequence database search tool. *Proteomics* **13**, 22–24 (2013).

5. Noble, W. S. Mass spectrometrists should search only for peptides they care about. *Nat. Methods* **12**, 605–608 (2015).

6. Sharma, V., Eng, J. K., Maccoss, M. J. & Riffle, M. A mass spectrometry proteomics data management platform. *Mol Cell Proteomics* **11**, 824–831 (2012).

7. Vizcaíno, J. A. *et al.* A guide to the Proteomics Identifications Database proteomics data repository. *Proteomics* vol. 9 4276–4283 (2009).

8. Jenkins, R. E. *et al.* Characterisation of flucloxacillin and 5-hydroxymethyl flucloxacillin haptenated HSA in vitro and in vivo. *Proteomics - Clin. Appl.* **3**, 720–729 (2009).

9. Parker, C. E., Perkins, J. R. & Tomer, K. B. Nanoscale packed capillary liquid chromatography- electrospray ionization mass spectrometry : analysis of penicillins and cephems. **616**, 45–51 (1993).

10. Creasy, D. M. & Cottrell, J. S. Unimod: Protein modifications for mass spectrometry. *Proteomics* vol. 4 1534–1536 (2004).

11. Kall, L. *et al.* Semi-supervised learning for peptide identification from shotgun proteomics datasets. *Nat. Methods* **4**, 923–925 (2007).

12. Deutsch, E. W. *et al.* A guided tour of the Trans-Proteomic Pipeline. *Proteomics* vol. 10 1150–1159 (2010).

13. McIlwain, S. *et al.* Crux: Rapid open source protein tandem mass spectrometry analysis. *J. Proteome Res.* **13**, 4488–4491 (2014).

14. Kong, A. T., Leprevost, F. V., Avtonomov, D. M., Mellacheruvu, D. & Nesvizhskii, A. I. MSFragger: Ultrafast and comprehensive peptide identification in mass spectrometry-based proteomics. *Nat. Methods* **14**, 513–520 (2017).

15. Chi, H. *et al.* Comprehensive identification of peptides in tandem mass spectra using an efficient open search engine. *Nat. Biotechnol.* **36**, 1059–1066 (2018).

16. Bagwan, N. *et al.* Comprehensive Quantification of the Modified Proteome Reveals Oxidative Heart Damage in Mitochondrial Heteroplasmy. *Cell Rep.* **23**, 3685–3697 (2018).

17. Solntsev, S. K., Shortreed, M. R., Frey, B. L. & Smith, L. M. Enhanced Global Post-translational Modification Discovery with MetaMorpheus. *J. Proteome Res.* **17**, 1844–1851 (2018).

18. Na, S., Bandeira, N. & Paek, E. Fast Multi-blind Modification Search through Tandem Mass Spectrometry. *Mol. Cell. Proteomics* **11**, M111.010199 (2012).

19. Devabhaktuni, A. *et al.* TagGraph reveals vast protein modification landscapes from large tandem mass spectrometry datasets. *Nat. Biotechnol.* **37**, 469–479 (2019).

20. Chick, J. M. *et al.* A mass-tolerant database search identifies a large proportion of unassigned spectra in shotgun proteomics as modified peptides. *Nat. Biotechnol.* **33**, 743–749 (2015).

21. Li, Q. *et al.* Global Post-Translational Modification Discovery. *J. Proteome Res.* **16**, 1383–1390 (2017).

22. David, M., Fertin, G., Rogniaux, H. & Tessier, D. SpecOMS: A Full Open Modification Search Method Performing All-to-All Spectra Comparisons within Minutes. *J. Proteome Res.* **16**, 3030–3038 (2017).

23. Shteynberg, D. D. *et al.* PTMProphet: Fast and Accurate Mass Modification Localization for the Trans-Proteomic Pipeline. *J. Proteome Res.* **18**, 4262–4272 (2019).

24. Pino, L. K. *et al.* The Skyline ecosystem: Informatics for quantitative mass spectrometry proteomics. *Mass Spectrom. Rev.* (2017) doi:10.1002/mas.21540.

25. MacLean, B. *et al.* Skyline: An open source document editor for creating and analyzing targeted proteomics experiments. *Bioinformatics* **26**, 966–968 (2010).

26. Schilling, B. *et al.* Platform-independent and label-free quantitation of proteomic data using MS1 extracted ion chromatograms in skyline: Application to protein acetylation and phosphorylation. *Mol. Cell. Proteomics* **11**, 202–214 (2012).

27. Baer, B. R., Wienkers, L. C. & Rock, D. A. Time-dependent inactivation of P450 3A4 by raloxifene: Identification of Cys239 as the site of apoprotein alkylation. *Chem. Res. Toxicol.* **20**, 954–964 (2007).
